# The balance between stability and plasticity of the Visual Word Form Area in dyslexia

**DOI:** 10.1101/2025.01.14.632854

**Authors:** Jamie L. Mitchell, Maya Yablonski, Hannah L. Stone, Mia Jimenez, Megumi E. Takada, Kenny A. Tang, Jasmine E. Tran, Clementine Chou, Jason D. Yeatman

## Abstract

Understanding the balance between plastic and persistent traits in the dyslexic brain is critical for developing effective interventions. This longitudinal intervention study examines the Visual Word Form Area in dyslexic and typical readers, exploring how this key component of the brain’s reading circuitry changes with learning. We find that children with dyslexia show significant differences in Visual Word Form Area presence, size, and tuning properties compared to typical readers. While reading intervention improves reading skills and increases Visual Word Form Area size, disparities persist a year later, suggesting that Visual Word Form Area abnormalities are enduring traits of dyslexia. Our results reveal long-term neural and behavioral changes, while also elucidating stable differences in the functional architecture of the dyslexic brain. This work provides comprehensive insights into the potential and limitations of short-term learning-induced plasticity in human visual cortex.

## Introduction

The Visual Word Form Area (VWFA) is a region of high-level visual cortex that is tuned to the visual features of written language^1,2^ and is thought to be directly linked to reading ability^3,4^. VWFA consists of at least 2 distinct sub-regions - the more posterior VWFA-1 which responds selectivity to the visual features of text and the more anterior VWFA-2 which is also sensitive to the linguistic properties of text^5,6^. This text-selective area of cortex is localized to the occipitotemporal sulcus (OTS)^7,8^, in the posterior and lateral portion of left ventral occipitotemporal cortex (VOTC). Because of its close relationship to reading ability, the VWFA is often only detected in literate adults^9^ and children who have already learned how to read^10–12^. Furthermore, VOTC is one of the most common anatomical regions cited in relation to reading disabilities (i.e., dyslexia)^13^. Previous research revealed that people with dyslexia display patterns of underactivation^14–16^ and decreased text-selectivity^11^ in VOTC compared to typical readers. Some suggest that this is due to disruption of typical function in this region for children with dyslexia^17,18^, and that this disruption is due to lack of functional specificity and increased sensitivity to non-text stimuli^19^. However, it remains unclear whether differences in this region are a stable trait of dyslexia or if text-selectivity in VOTC can change as struggling readers improve their reading ability through targeted intervention.

Understanding the dynamic interplay between plasticity and stability is essential for translating research findings into impactful clinical, educational, and policy applications^20^. This understanding is particularly crucial when evaluating the persistence of biological markers even as behavioral improvements are observed. To date, while there has been extensive research on the effects of reading interventions on the brain’s reading circuitry^21,22^, gaps remain in understanding which functional properties are malleable and which are stable traits distinguishing dyslexic from typical readers. Recent evidence suggests that effective reading intervention drives structural changes in white matter tracts involved in reading^23,24^. However, findings regarding the intervention-driven changes in the functional properties of VOTC have been mixed. While some studies report that intervention increases VOTC activation^25–27^, others do not^21,28^ Further, the long-term effects of short-term interventions are largely unknown^25^, making the timescale of plasticity and the balance between plasticity and stability over different timescales an open question.

To address these gaps, in the present study we assessed the functional properties of VOTC in a large group of children with dyslexia who participated in a targeted reading intervention to increase reading ability using a precision neuroimaging approach where regions are localized on each individual’s cortical surface. Many previous intervention studies have asked questions about what changes (broadly construed) but no previous study has specifically targeted the function of the VWFA by localizing the region using a task that was specifically designed to drive the response of this region. Our study targeted this specific question and we used a task that was developed around a computational model of VWFA tuning properties^30^. We used dense neuroimaging sampling (up to 5 times over the course of one year) to detail the short- and long-term learning-induced effects on the functional properties of the VWFA. We use longitudinal modeling - specifically models that separate changes that are linked to behavior (state) from differences that are stable over the 1 year study timeline (trait) - to characterize relationships between intervention, VWFA function, and behavioral outcomes of reading ability. We ask whether differences in the VWFA are a stable trait of dyslexia that persists after the intervention or are ameliorated as reading skills improve.

## Results

Forty-four children with dyslexia who participated in a reading intervention, and 43 controls with (*n*=19) and without dyslexia (*n*=24) underwent functional magnetic resonance imaging (fMRI) and behavioral assessments for up to 5 time points over the course of a year. Participants (ages 7-13 years at the start of the study) completed an fMRI experiment (Figure 1a) consisting of a child-friendly adaptation of the localizer developed by White et al, ^31^ which examines fMRI responses to high and low frequency words, pseudowords, consonant strings, pseudo fonts, faces, limbs and objects under different task demands^32^.

**Figure 1.**
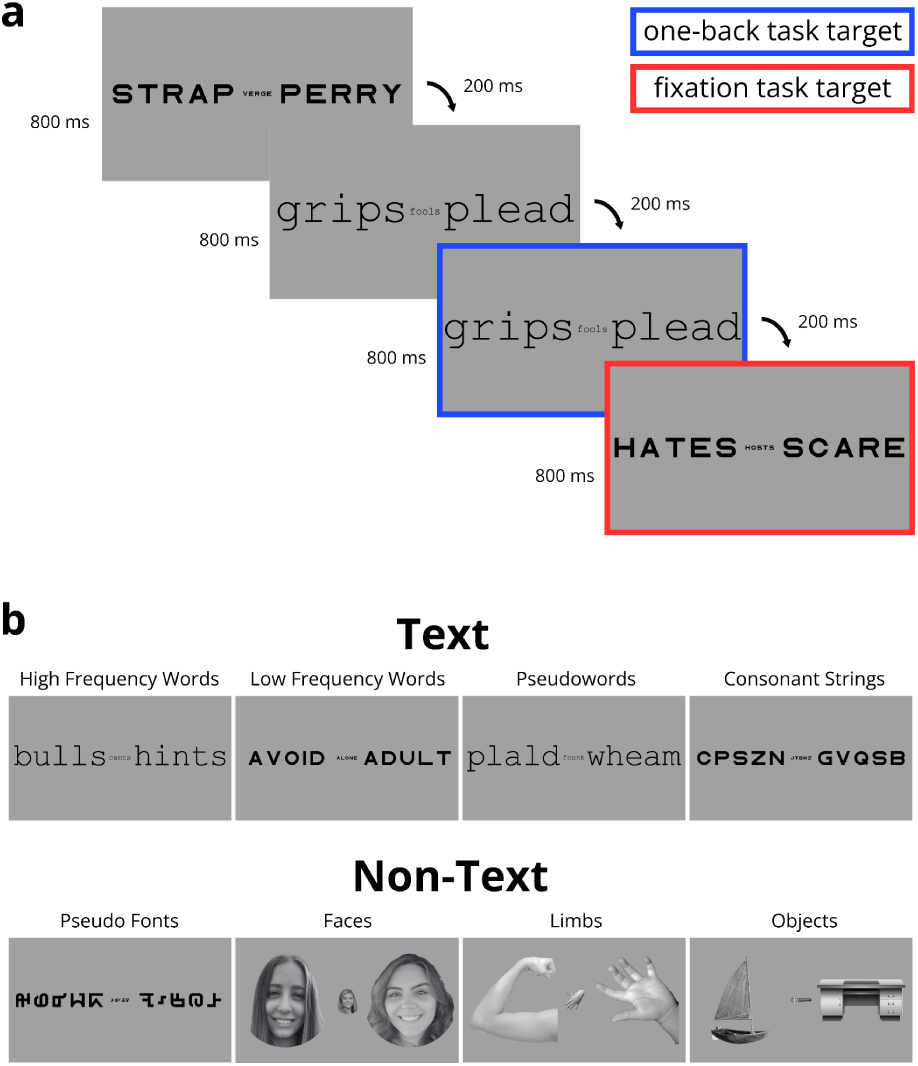
Functional Localizer Experimental Design. **a,** Example of a single trial of the experiment. Each frame is presented for 800 ms followed by a blank fixation screen (not shown in figure) for 200 ms. Each trial contains 4 frames of the same stimulus category and lasts for 4 seconds. Targets for the one-back repetition are highlighted in blue and targets for the fixation color change task are highlighted in red (no colors were used in the actual experiment). **b**, Sample stimuli from each sub-category. The text category (top) is comprised of high- and low-frequency words, pseudo words, and consonant strings. The non-text category (bottom) is comprised of pseudo fonts, faces (male and female), limbs (hands, arms, feet, and legs), and objects. Shown in this illustration are faces of co-authors, but the actual experiment presented faces of people unfamiliar to participants.

Left and right Visual Word Form Area (VWFA) 1 and 2 (posterior to anterior) and Fusiform Face Area (FFA) 1 and 2 (posterior to anterior) were defined on the native cortical surface (see Figure 2a) of each participant at every time point based on the text>non-text and faces>other contrasts (threshold of *t*>3 within the appropriate anatomical boundaries (see Methods for exclusion criteria). For longitudinal analyses of functional selectivity, additional ROIs were defined by combining all timepoints available for each subject, to increase SNR as well as ensure a consistent ROI across timepoints. Posterior and anterior portions of VWFAs and FFAs were defined separately to explore the nuanced differences of functional properties of these distinct regions and how they might differentially relate to learning. To determine any differences in these functionally-defined ROIs, we compared the presence, size, and functional selectivity of each ROI both at the baseline across participants with different levels of reading ability, and longitudinally as reading skills improved. FFAs were included in the analysis as a control region, as it does not respond selectively to text and would not change with increases in reading ability.

**Fig. 2.**
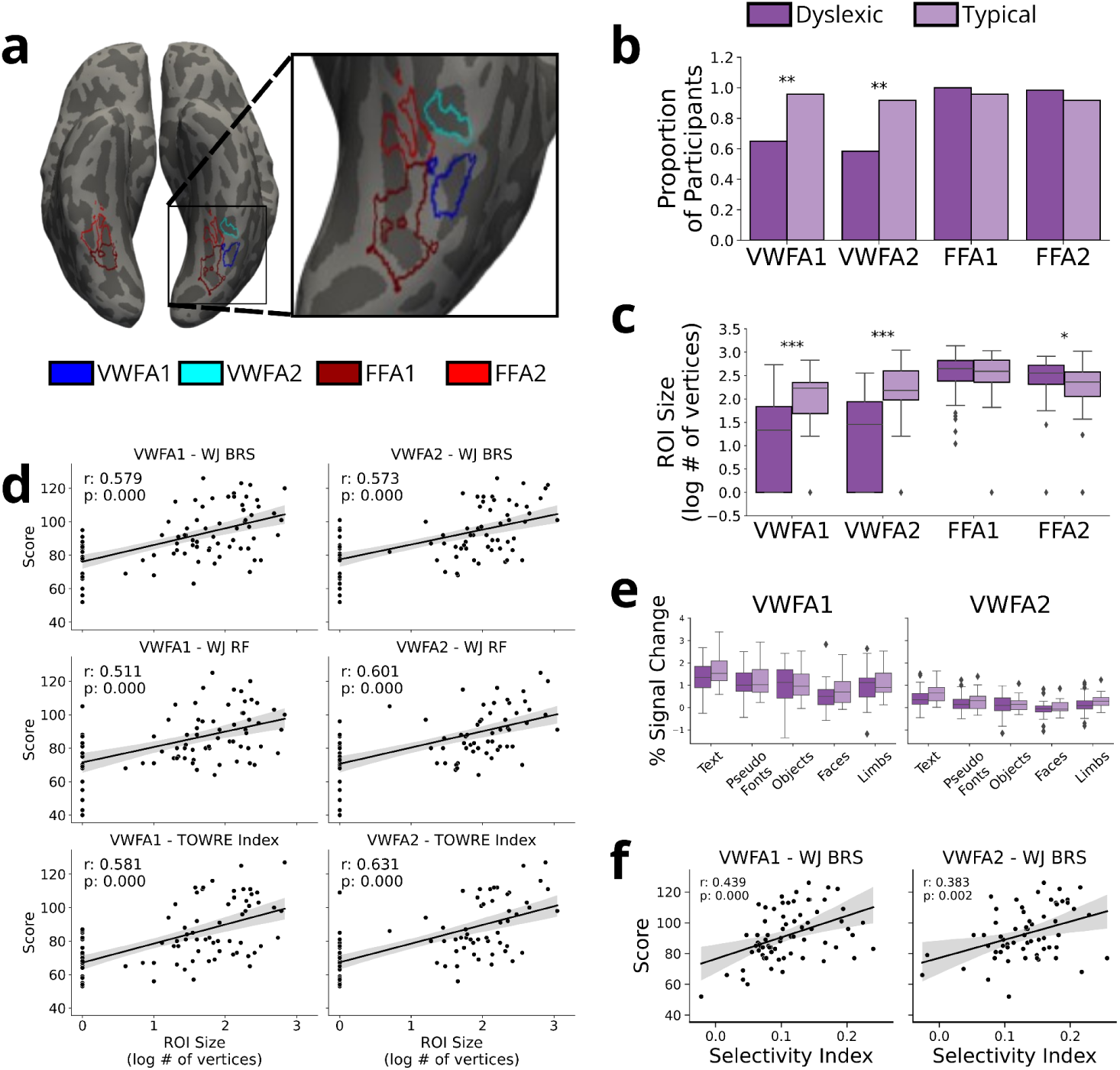
Visual Word Form Area Size and Selectivity Is Related to Reading Ability at Baseline. **a,** Example regions of interest (ROIs) on the inflated surface of one example subject. Visual Word Form Area 1 (VWFA-1; blue) & VWFA-2 (cyan) fall on the left occipitotemporal sulcus (OTS) and Fusiform Face Area 1 (FFA-1; dark red) & FFA-2 (red) fall on the fusiform gyrus. **b**, Proportion of participants with usable data who had a VWFA or FFA of any size present at baseline in typical (light purple) and dyslexic (purple) readers for each ROI. VWFA1: p=0.009; VWFA2: p=0.007; FFA1: p=0.286; FFA2: p=0.403. **c**, Box plot of log transformed size (in number of vertices) at baseline in typical (n=24) and dyslexic readers (n=60) for each ROI. VWFA1: p=1.50E-05; VWFA2: p=1.56E-05; FFA1: p = 0.706; FFA2: p=0.026. **d**, Results from a two-sided Pearson correlation of VWFA size and reading assessment scores: Woodcock-Johnson Basic Reading Skill score (WJ BRS), Reading Fluency score (WJ RF), and Test of Word Reading Efficiency Index (TOWRE). **e**, Activation (in units of percent signal change) for each category of visual stimuli (text, pseudo fonts, objects, face, & limbs), within VWFA-1 (left; n=47 dyslexic & 23 typical) and VWFA-2 (right; n=43 dyslexic & 21 typical). Outliers in the data are individually plotted above and below box whiskers. **f**, Results from a two-sided Pearson correlation of selectivity index in VWFA-1 & VWFA-2 and BRS scores across all participants. Asterisks in panels b and c indicate the degree of significance derived from a two-sides *t* test (p < 0.001: ***, p < 0.01: **, p < 0.05: *, p < 0.1: .) Box plots in panels c and e display the median indicator, the box displays the interquartile range (IQR; 25%-75% range), and whiskers represent +/− 1.5 times the IQR. Outliers in the data are individually plotted above and below box whiskers. Error bands in panels d and f represent that 95% confidence interval. Source data are provided in a public data repository.

### VWFA Presence, Size, and Selectivity is related to Reading Ability

We first tested the hypothesis that the VWFA is more likely to be absent (or harder to detect) in children with dyslexia^11^. To this end, we focused on the baseline (pre-intervention) time point, and compared all participants with dyslexia (*n*=59) to all typical readers (*n*=24). A *χ*^2^ test of independence confirmed that, compared to typical readers, in this specific timepoint, a smaller proportion of dyslexic participants had a detectable VWFA-1 (dys=65%, typ=95.83%; *χ*^2^ = 6.911, *p* = 0.009) and VWFA-2 (dys=58.33%, typ=91.66%; *χ*^2^ = 7.272, *p* = 0.007) at baseline (Figure 2b). There was no difference in the presence of FFAs between dyslexic and typical readers at baseline (FFA1: *χ*^2^ = inf, *p* = 0.286; FFA2: *χ*^2^ = 0.700, *p* = 0.403) confirming the specific role of the VWFA as opposed to a general difference extending across VOTC regions. Note that as a general rule, in all analyses of ROI presence and size we take into account ROIs defined on each specific time point. However when we compare measurements of activation amplitude (including text-selectivity), we extract session specific activations from ROIs defined on data from all time points. This is done to avoid conflating the ROI size in any specific time point with response amplitude (see Methods for more detail).

We next sought to determine if the VWFA is smaller in participants with dyslexia compared to typical controls. With this analysis, we wanted to determine if there is a difference to the amount of cortical territory devoted to text-processing for struggling readers. To do so, we used individually defined ROIs in the baseline timepoint for each participant to determine the size of a given region in units of surface mesh vertices. A *t*-test (Table S1) revealed that participants with dyslexia had significantly smaller ROIs than typical readers for both VWFA-1 (mean dys= 58.833 vertices, mean typ = 189.542 vertices; *t*(82) = −4.603, *p* = 1.50E-05, %CI = (−1.348,-0.534)) and VWFA-2 (mean dys= 62.650 vertices, mean typ = 285.042 vertices; *t*(82) = −4.593, *p* = 1.65E-05, %CI = (−1.491, −0.590)) but not for FFA-1 (mean dys= 471.600 vertices, mean typ = 471.417 vertices; *t*(82) = 0.378, *p* = 0.706, %CI = (−0.190, 0.280)). Size differences were present in FFA-2 (mean dys= 362.333 vertices, mean typ = 275.8775 vertices; *t*(82) = 2.262, *p* = 0.026, %CI = (0.037, 0.577), see Figure 2c), though not to the same extent as the VWFAs. Furthermore, these findings regarding both VWFAs held true even when we excluded individuals without a VWFA from the analysis (See Supplementary Table S2).To better understand the relationship between VWFA size and reading ability, we next calculated the correlation between ROI size and reading ability as a continuous measure. We used three different reading assessments to capture different aspects of reading: Woodcock-Johnson Basic Reading Skills score (WJ BRS), Woodcock-Johnson Reading Fluency score (WJ RF), and Test of Word Reading Efficiency index (TOWRE). We found that the size of both VWFA-1 and VWFA-2 was positively correlated with reading ability for each of the three reading measures (*p* < 0.000; See Figure 2d & Table S3). In contrast, there was no significant correlation between reading ability and FFA size (all *p* > 0.1). To ensure that this relationship was not driven solely by the individuals who were missing an ROI (i.e., participants with ROI size of 0), we re-ran the analysis excluding these individuals and found that the significant relationship persisted even within the smaller sample (Supplementary Figure S1). This robust individual approach to defining ROIs helps to clarify known differences in tuning properties of the VOTC for dyslexic and typical readers in a novel way by describing the differences in the amount of cortical territory that is selective for reading.

We then tested the hypothesis that neural tuning properties of the VWFA differ in participants with dyslexia compared to typical controls. Specifically, we calculated response amplitudes to each stimulus condition within each participant’s individually-defined, VWFA ROIs. Note that for this analysis we used ROIs defined on data from all timepoints, to avoid conflating the ROIs size with response amplitude (see Methods for more detail). To determine any difference in activation we ran a linear mixed effects model (LME) looking at the interaction between group and stimulus category, (Supplementary Table S4). We found a group effect (dyslexic < typical) on the mean percent signal change to text in VWFA-1 (*β*(87) = 0.353, *p* = 0.046, SE = 0.174) and in VWFA-2 (*β*(79) = 0.213, *p* = 0.0047, SE = 0.105; Figure 2e). Of note, since our analyses were not pre-registered, these marginal effects should be interpreted with caution. However, combined with our previous analyses, these findings support past work suggesting that dyslexic participants have weaker text-evoked activation in these regions compared to typical peers^17,18^. Interestingly, there was a significant interaction showing that the difference between text activation and object activation was larger in the typical reader group compared with the dyslexic group (VWFA-1: *β*(272) = −0.380, *p* = 0.2.88E-04, SE = 0.103; VWFA-2: *β*(248) = −0.206, *p* = 9.57E-04, SE = 0.062). This suggests that children with dyslexia have relatively stronger responses to objects, in line with Kubota et al.^11^, in addition to weaker responses to text.

We then calculated a selectivity index for each participant in the sample, defined as the difference between activation to text versus non text, divided by the sum of activation to all stimuli^11^. We calculated the correlations between text selectivity and assessment score (Table S5) as a complement to the above percent signal change analysis and to narrow in on text-specific differences to activation patterns. We found a significant positive relationship between all the reading scores and selectivity in VWFA-1 (WJ BRS: *r*(67) = 0.439, *p* = 1.61E-04, %CI = (0.226, 0.612); WJ RF: *r*(66) = 0.403, *p* = 6.53E-04, %CI = (0.182, 0.585); TOWRE: *r*(64) = 0.356, *p* = 0.003, %CI = (0.125, 0.551)) and with selectivity in VWFA-2 (WJ BRS: *r*(60) = 0.383, *p* = 0.002, %CI = (0.147, 0.578); WJ RF: *r*(59) = 0.400, p = 0.001, %CI = (0.165, 0.592); TOWRE: *r*(57) = 0.397, *p* = 0.002, %CI = (0.157, 0.593); Figure 2f). This confirms findings from previous studies that better reading is associated with greater selectivity for text in the VWFA^11^.

### Reading Improvement Drives Changes in VWFA

To first determine if reading ability improved as a result of the intervention, we fit a LME looking at assessment score as a function of time (in days from baseline) with a random intercept by participant for each study group (Table S6). We confirmed that the reading intervention successfully improved reading ability in the intervention group (WJ BRS: *β*(168) = 0.021, *p* = 1.61E-11, SE = 0.003; WJ RF: *β*(163) = 0.020, *p* = 1.77E-13, SE = 0.002; TOWRE: *β*(165) = 0.026, *p* = 5.74E-25, SE = 0.002 - Figure 3a). As expected, the dyslexic control (WJ BRS: *β*(42) = −0.004, *p* = 0.487, SE = 0.006; TOWRE: *β*(42) = 0.003, *p*= 0.513, SE = 0.005) and typical control (WJ BRS: *β*(50) = −0.006, *p* = 0.173, SE = 0.005; TOWRE: *β*(46) = −0.004, *p* = 0.481, SE = 0.005) groups showed almost no changes in reading scores. Interestingly, both control groups did show increases in WJ RF (dyslexic - *β*(42) = 0.008, *p* = 0.043, SE = 0.004; typical - *β*(50) = 0.020, *p* = 8.23E-05, SE = 0.005) despite displaying no significant gains in any other reading assessment. These results were further supported by a second model fit with the same parameters along with a group interaction (Table S7). This interaction model confirmed large increases in reading score across all measures for the intervention group (*p*<0.001) and significantly smaller rates of change in score for the control groups for WJ BRS (Dys Ctrl: *β*(265) = −0.025, *p* = 1.36E-04, SE = 0.006; Typ Ctrl: *β*(268) = −0.028, *p* = 2.53E-05, SE = 0.006) and TOWRE (Dys Ctrl: *β*(257) = −0.023, *p* = 2.15E-05, SE = 0.005; Typ Ctrl: *β*(256) = −0.029, *p* = 3.61E-08, SE = 0.005). This significant interaction suggests that the reading improvement was specifically driven by the intervention itself. In contrast, while reading fluency and math (WJ Math Facts Fluency - WJ MFF) improved over time for the intervention group (WJ RF: *β*(254) = 0.020, *p* = 2.03E-15, SE = 0.002; WJ MFF: *β*(258) = 0.007, *p* = 0,010, SE = 0.003), this effect was not specific to the intervention group (WJ RF: *p* = 0.894 - typical controls only; WJ MFF: *p* > 0.200 - both control groups). After establishing that the intervention was effective in improving reading skills in the intervention group alone, we next tested the hypothesis that this reading improvement drives plasticity in VWFA. As our results suggest that the intervention group was the only group to experience large change in reading ability, we explored the remaining analyses with group comparisons. The intention here is that the intervention group should be thought of as a group that underwent significant growth in reading ability whereas the control groups should be thought of as groups that did not undergo drastic improvement.

**Figure 3.**
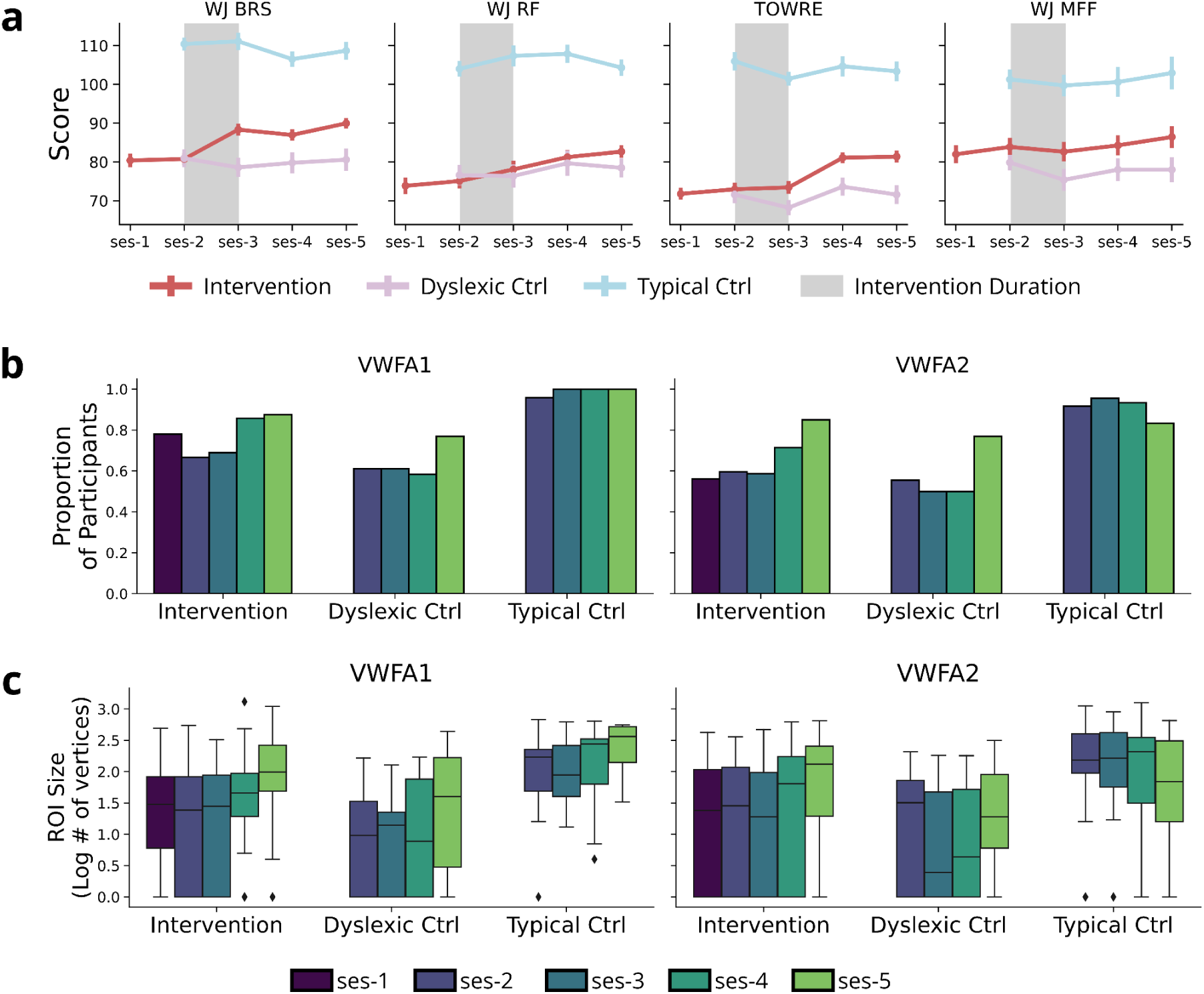
Change in Reading and Visual Word Form Area. **a,** Change in standard score for various reading assessments (Woodcock-Johnson Basic Reading Skills - WJ BRS; Woodcock-Johnson Reading Fluency - WJ RF; and Test of Word Reading Efficiency - TOWRE) and a control math assessment in all three groups of participants. Data are displayed as the mean value with error bars representing the standard error. **b**, Proportion of participants with usable data who had a Visual Word Form Area (VWFA) 1 and 2 of any size present at every time point in each group. **c**, Size (log transformed number of vertices) at every time point in each group for VWFA-1 and VWFA-2. Box plots display the median indicator, the box displays the interquartile range (IQR; 25%-75% range), and whiskers represent +/− 1.5 times the IQR. Outliers in the data are individually plotted above and below box whiskers. N = 44 intervention, 19 dyslexic & 24 typical controls. Source data are provided in a public data repository.

We first determined if the proportion of detectable VWFAs increased after the intervention in children with dyslexia. To do so, we fit generalized linear mixed effects (GLME) models (binomial family for a dichotomous outcome measure) within each group (intervention, dyslexic control, and typical control), predicting the presence of a VWFA ROI as a function of time (in days from baseline). We also included regressors of age (at baseline visit), movement (mean framewise displacement calculated separately for each session), and number of runs per session (after exclusion for data quality, see Methods) to control for effects of data quality or developmental factors (unless otherwise specified all models include these covariates). Using this approach, we found that the intervention group experiences significant increases in the probability of detecting VWFA-1 (*β* = 0.005, *p* = 0.023, SE = 0) and VWFA-2 (*β* = 0.009, *p* = 8.16E-04, SE = 0). The dyslexic control group showed no change in probability of detecting VWFA-1 (*β* = 0.005, *p* = 0.223, SE = 0) and VWFA-2 (*β* = 0.008, *p* = 0.114, SE = 0) and the typical control group actually showed a decrease in probability of detecting VWFA-2 (*β* = −0.101, *p* = 0.032, SE = 0). Since nearly every participant in the typical control group had a well-defined VWFA-1 from the beginning of the study, we could not fit a GLMER model to the control group here. Due to similar model convergence issues changes in some FFAs for groups were not able to be calculated (See Table S8 for a complete breakdown).

To examine changes in VWFA size, we ran a LME where we estimated the size of each separate session ROI (in log transformed number of vertices) as a function of the interaction between group and time with a random intercept for participant, treating the intervention group as the reference (Figure 3C; Table S9). We found that the ROI size increased with time in the intervention group for both VWFA-1 (*β*(248) = 0.320, *p* = 1.12E-07, SE = 0.059; ∼2.090 vertices per day) and VWFA-2 (*β*(238)= 0.245, *p* = 1.37E-08, SE = 0.059; ∼1.760 vertices) while there was a negative interaction effect with time for the typical control group in VWFA-2 (*β*(242) = −0.233, *p* = 0.014, SE = 0.094), indicating that the typical control group did not show growth in VWFA-2. Furthermore, compared to the intervention group, the typical control group had a larger VWFA-1 (*β*(124) = 103.273, *p* = 0.004, SE = 35.274) and a larger VWFA-2 (*β*(90) = 217.092, *p* = 4.55E-06, SE = 44.493). This once again confirms that children with dyslexia have smaller VWFAs than typical readers. As an alternative analysis approach to determine if this size difference persisted throughout the course of the study, we ran a series of two-sample t-tests comparing the size of the intervention participants’ VWFAs to the size of the typical controls’ VWFAs (see Table S10). In doing so, we found that by the final time point, size differences between the two groups disappeared in VWFA-2 (*t* = 0.106, *p* = 0.917), however the difference still remained in VWFA-1 (*t* = −3.037, *p* = 0.004). Furthermore, a correlation between VWFA size and all reading ability measures remained (*p* ≤ 0.041, see Figure S2) even when constrained to data from the last timepoint, further emphasizing the continued relationship between reading ability and VWFA size. Finally, no size changes over time were observed in control FFA regions.

To ensure that changes in VWFA size were truly the result of a change in tuning properties of the VOTC and not simply the effect of methodological decisions such as the threshold used to define the ROIs, we ran additional analyses on VWFA size at various thresholds. To accomplish this, we drew a large anatomical VOTC ROI in fsaverage template space, projected this label onto the native cortical surface for each intervention participant, and used this label to constrain text-selective contrast maps. We then selected all vertices within these VOTC-masked contrast maps that had a *t* value greater than or equal to different threshold values, in increments of 0.5 ranging from 0.5 to 4.5. Figure 4 shows that regardless of the chosen threshold, intervention participants did in fact show an increase in the number of text-selective vertices in VOTC over time. Furthermore, to ensure that the relationship between reading ability and VWFA size was also maintained across thresholds, we repeated the analysis visualized in Figure 2d and computed correlations between pre-intervention VWFA size and assessment score for each threshold (Supplementary Figure S3). We found that the relationship between reading assessment ability and VWFA was still highly correlated at every threshold and with every reading assessment (*p* < 0.000).

**Figure 4.**
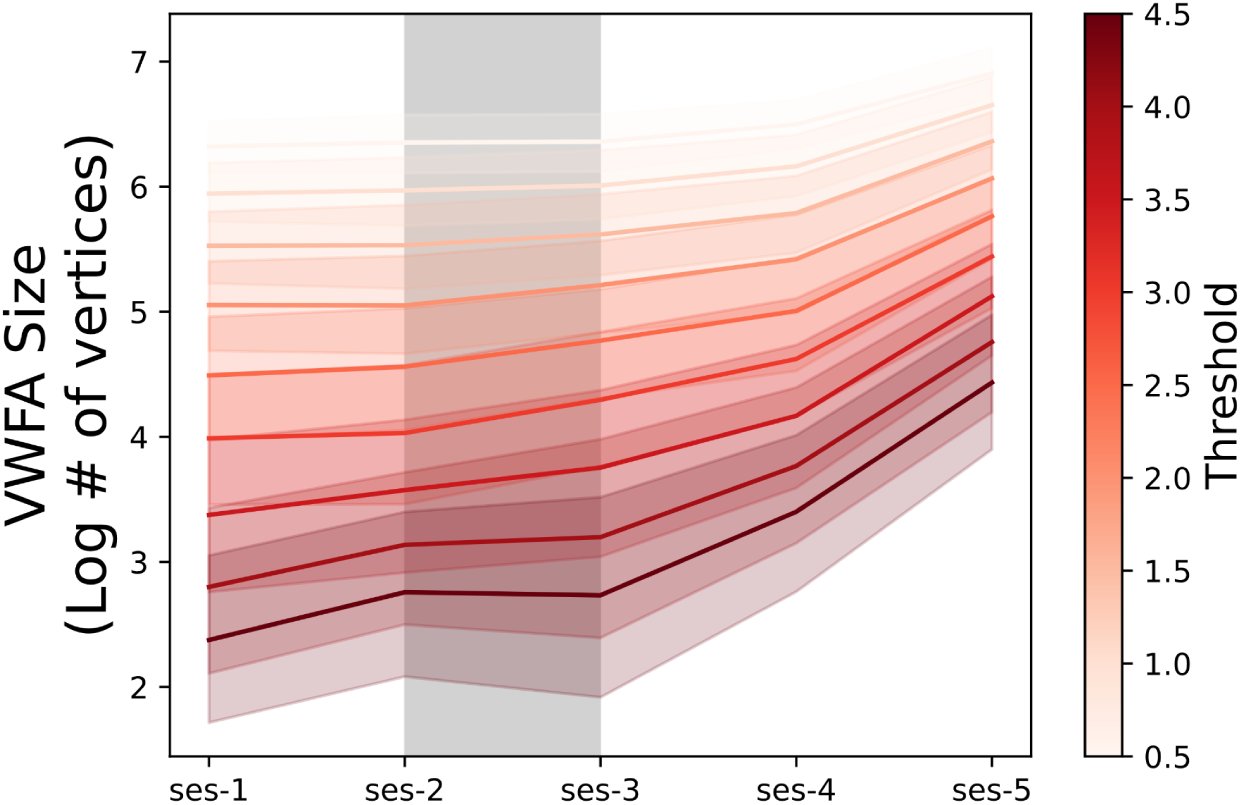
Visual Word Form Area Shows Growth Across Thresholds. Change in size of VWFA in the intervention participants (n=44). For the purposes of this analysis, Visual Word Form Area (VWFA) was defined as any vertex in the Ventral Occipitotemporal Cortex that had a *t* value >= the threshold value in a contrast map comparing activation to text > activation to all other categories. Thresholds were tested in increments of 0.5 ranging from 0.5 to 4.5. Intervention period is visualized with a grey box between sessions 2 and 3. Source data are provided in a public data repository.

### VWFA Differences Persist After Intervention

We sought to determine if the differences that existed between typical and dyslexic readers at baseline persisted after the conclusion of the intervention, despite confirmed increases in the presence and size of VWFA in the intervention group. We fit a linear mixed effects model looking at percent signal change as a function of the interaction between group, time, and visual category, with a random intercept of participant (Figure 5a, Table S11). This model revealed that in VWFA-1, typical readers had a greater magnitude of percent signal change to text than intervention participants (*β*(77) = 0.372, *p* = 0.024, SE = 0.162), and that the difference between activation to text and non-text stimuli was greater for typical readers than intervention participants (*β*(481) = −0.179, *p* = 0.007, 0.066), further confirming our cross-sectional results and findings from previous studies regarding hypoactivation of the region in dyslexic readers^17,18^. There were no changes in activations for the intervention group over time in VWFA-1 (*β*(482) = 0.000, *p* = 0.158, SE < 0.000), suggesting that the group effect observed at baseline persists despite intervention in VWFA-1. In VWFA-2, the intervention group showed increased signal to text over time (*β*(425) = 0.0003, *p* = 4.08E-04, SE < 0.000), mirroring findings from previous work^27^. To determine if the elevated object sensitivity in VWFA-2 for dyslexic readers seen at baseline persisted after intervention, we ran a post-hoc analysis comparing mean percent signal change from baseline (time point 2) to the final time point (time point 5) in the intervention participants. A paired samples t-test revealed that while response to text increased significantly in the intervention group (*t*(29) = 3.144, *p* = 0.004), responses to objects only marginally increased (*t*(29) = 2.206, *p* = 0.035).

**Figure 5.**
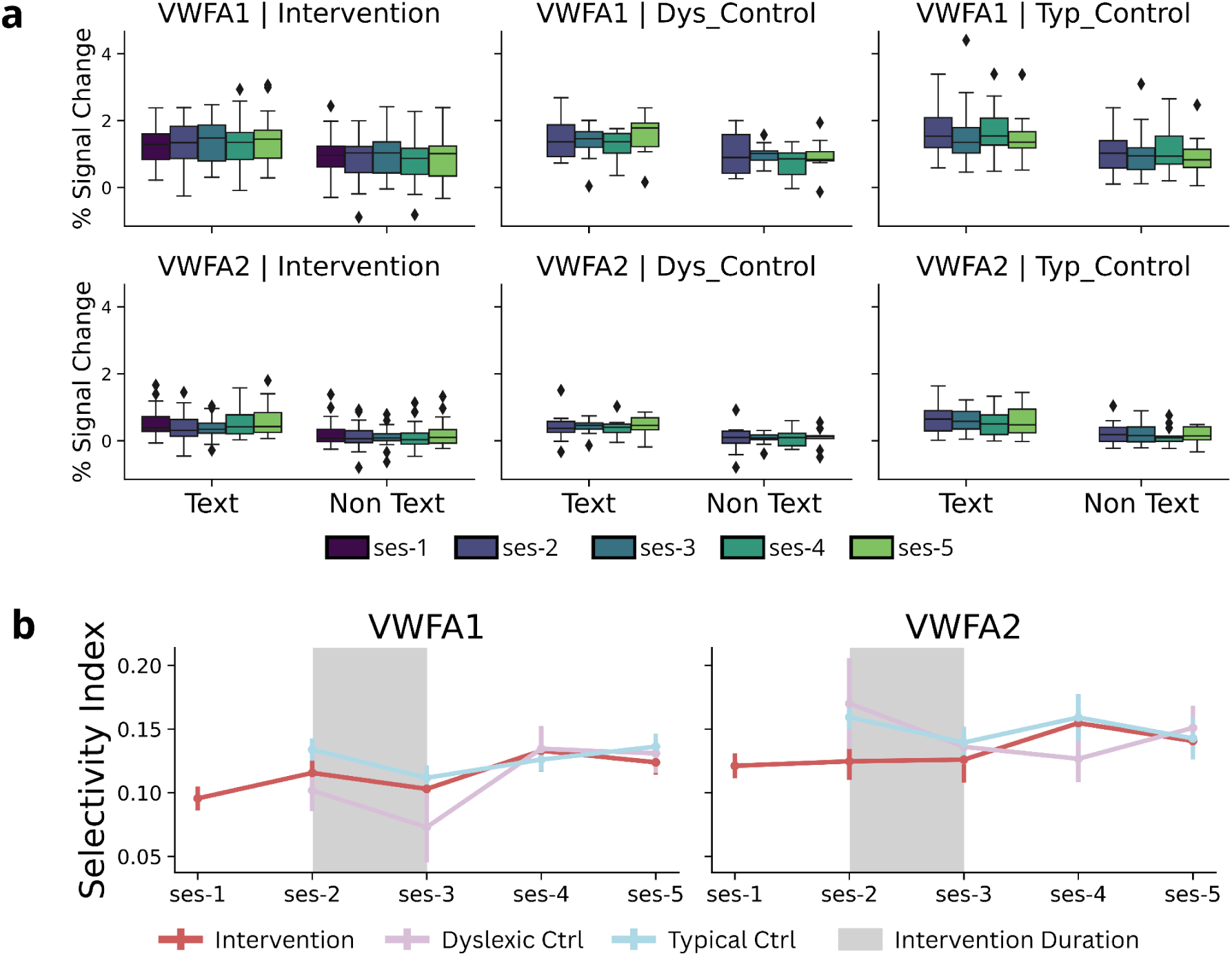
VWFA Does Not Show Learning-Induced Increases of Text Sensitivity. **a,** Activation (in units of percent signal change) to text and non-text categories, at each time point, within Visual Word Form area (VWFA) 1 and 2. Box plots display the median indicator, the box displays the interquartile range (IQR; 25%-75% range), and whiskers represent +/− 1.5 times the IQR. Outliers in the data are individually plotted above and below box whiskers. **b**, Change in selectivity index across each time point in VWFA-1 and VWFA-2. Data are displayed as the mean value with error bars representing the standard error. Intervention period is visualized with a grey box between sessions 2 and 3. N = 44 intervention, 19 dyslexic & 24 typical controls. Note: Data for WASI were collected at a pre-study screener when participants entered a participant recruitment dashboard. Time between WASI score and start of the study varies by participant. Source data are provided in a public data repository.

To further investigate any longitudinal change in tuning properties of the VWFA, we ran another LME looking at text selectivity index as a function of the interaction between time and group (Figure 5b, Table S12). We found that the intervention group had a small but significant increase in text-selectivity in VWFA-1 (*β*(215) = 0.0001, *p* = 0.013, SE < 0.000) and in VWFA-2 (*β*(189) = 0.0001, *p* = 0.044, SE < 0.000). We also found no effect of group nor an interaction effect of group and time indicating that the trajectory of selectivity index change over time was not significantly different in the intervention group in comparison to the control groups. This is likely due to the fact that, qualitatively, we see more variability in the control groups text-selectivity changes (See Figure 5b), indicating that a longer time scale may be needed to observe stable trends within control groups.

### Visual Word Form Area size tracks growth in reading ability

After determining that VWFA size changes with time, we next sought to investigate whether this growth is coupled with reading improvement. To this end, we calculated a reading “trait” score for each assessment for every participant by computing an average score across all available time points. We next calculated a reading state score by subtracting each participant’s trait score from their score in each time point (see Table S13 for descriptive summary table). We use trait and state scores to tease apart the effects of within participant change in score from the between-participant individual differences (captured by the trait scores)^33^. We then ran a linear mixed effects model looking at ROI size as a function of reading trait and the interaction between reading state and study group, with a random intercept of participant (Tables 1 & 2). This analysis revealed a main effect of reading trait across all reading assessments in both VWFA-1 (WJ BRS: *β*(88) = 0.020, *p* = 0.009, SE = 0.007; WJ RF: *β*(86) = 0.013, *p* = 0.028, SE = 0.006; TOWRE: *β*(87) = 0.022, *p* = 0.004, SE = 0.007) and VWFA-2 (WJ BRS: *β*(89) = 0.036, *p* = 1.38E-06, SE = 0.009; WJ RF: *β*(88) = 0.030, *p* = 0.5.47E-05, SE = 0.007; TOWRE: *β*(89) = 0.047, *p* = 3.02E-07, SE = 0.008). This was not the case for the control math assessment (VWFA-1: *β*(77) = 0.001, *p* = 0.868, SE = 0.005; VWFA-2: *β*(80) = 0.010, *p* = 0.122, SE = 0.006). This corroborates our findings regarding the pre-intervention time point and further supports our hypothesis that a small VWFA is related to weaker reading ability. The model also revealed a main effect of reading state across all reading assessments on both VWFA-1 size (WJ BRS: *β*(239) = 0.023, *p* = 1.00E-03, SE = 0.007; WJ RF: *β*(241) = 0.039, *p* = 8.06E-07, SE = 0.008; TOWRE: *β*(239) = 0.025, *p* = 0.002, SE = 0.008) and VWFA-2 size (WJ BRS: *β*(239) =0.031, *p* =4.08E-06, SE = 0.006; WJ RF: *β*(240) = 0.051, *p* = 1.42E-11, SE = 0.007; TOWRE: *β*(238) = 0.037, *p* =7.54E-07, SE = 0.007). This suggests that a greater increase in reading ability is related to a greater increase in VWFA size. Surprisingly, for WJ RF, dyslexic controls displayed a weaker relationship between reading trait and VWFA-1 size (*β*(82) = −0.361, *p* = 0.037, SE = 0.170). Additionally, for typical controls, there was a weaker relationship between change in reading state and change in ROI size in VWFA-2 (*β*(245) = −0.040, *p* = 0.036, SE = 0.019) for WJ BRS and for both VWFA-1 (*β*(234) = −0.039, *p* = 0.016, SE = 0.016) and VWFA-2 (*β*(236) = −0.063, *p* = 4.52E-05, SE = 0.015) for WJ RF. This suggests that change in word reading and reading fluency may have a greater relationship to change in VWFA size than our reading efficiency. These findings were maintained when re-running models on individual assessment raw scores as opposed to composite standard scores (Supplementary Table S14). Furthermore, this analysis approach was repeated on measures of VWFA presence, percent signal change, and selectivity and results can be found in Supplementary Tables S15, S16, and S17 respectively.

**Table 1.** Longitudinal Results for Change in VWFA Size with Trait and State Scores. Results of linear mixed effects models (LME) calculating the size (log transformed number of vertices) of a region of interest (ROI) as a function of assessment trait and the interaction between assessment state and participant group with added covariates of participant age, mean framewise displacement during the scan (mean FD), and the number of usable runs of the experiment along with a random intercept by participant. Intervention group is treated as the reference category. Trait: within-participant average over time; State: within-participant change relative to trait; VWFA: Visual Word Form Area; WJ BRS: Woodcock-Johnson Basic Reading Skills; WJ RF: Woodcock-Johnson Reading Fluency; TOWRE: Test of Word Reading Efficiency; WJ MFF: Woodcock-Johnson Math Facts Fluency; Dys Ctrl: Dyslexic Control; Typ Ctrl: Typical Control; FD: Framewise Displacement Significant results are displayed in bold and asterisks indicate the degree of significance (p < 0.001: ***, p < 0.01: **, p < 0.05: *, p < 0.1: .). Significant results of p<0.05 are indicated with bold font. Note: No corrections for multiple comparisons were made due to the use of small, manually-defined ROIs. Source data are provided in a public data repository.

| Log Size ~ Reading State*Subgroup Age + Movement + RunNums + Reading Trait + (1 \| Participant) |  |  |  |  |  |  |  |  |  |  |
| --- | --- | --- | --- | --- | --- | --- | --- | --- | --- | --- |
|  | VWFA1 |  |  |  |  | VWFA2 |  |  |  |  |
| | $\beta$ | Std. Err | DOF | t | p | $\beta$ | Std. Err | DOF | t | p |
| **WJ BRS** |  |  |  |  |  |  |  |  |  |  |
| Intercept (Intervention) | **-2.137** | **0.835** | **114** | **-2.560** | **0.012\*** | **-3.074** | **0.987** | **108** | **-3.113** | **0.002\*\*** |
| WJ BRS Trait | **0.020** | **0.007** | **88** | **2.684** | **0.009\*\*** | **0.036** | **0.009** | **89** | **3.984** | **1.38E-04\*\*\*** |
| WJ BRS State | **0.023** | **0.007** | **239** | **3.418** | **1.00E-03\*\*\*** | **0.031** | **0.006** | **239** | **4.717** | **4.08E-06\*\*\*** |
| Group: Dys Ctrl | -0.251 | 0.175 | 82 | -1.435 | 0.155 | -0.024 | 0.214 | 84 | -0.113 | 0.911 |
| Group: Typ Ctrl | 0.177 | 0.239 | 83 | 0.741 | 0.461 | -0.089 | 0.292 | 85 | -0.304 | 0.762 |
| Age | **0.176** | **0.050** | **82** | **3.551** | **1.00E-03\*\*\*** | 0.105 | 0.061 | 85 | 1.731 | 0.087. |
| Movement (Mean FD) | -0.753 | 0.391 | 314 | -1.925 | 0.055. | -0.552 | 0.390 | 300 | -1.415 | 0.158 |
| Num Good Runs | 0.069 | 0.069 | 300 | 0.997 | 0.319 | 0.098 | 0.068 | 282 | 1.435 | 0.152 |
| Dys Ctrl * WJ BRS State | -0.018 | 0.017 | 238 | -1.066 | 0.287 | -0.022 | 0.016 | 239 | -1.387 | 0.167 |
| Typ Ctrl * WJ BRS State | -0.039 | 0.020 | 247 | -1.935 | 0.054. | **-0.040** | **0.019** | **245** | **-2.106** | **0.036\*** |
| **WJ RF** |  |  |  |  |  |  |  |  |  |  |
| Intercept (Intervention) | -1.136 | 0.681 | 117 | -1.667 | 0.098. | **-1.585** | **0.792** | **108** | **-2.001** | **0.048\*** |
| WJ RF Trait | **0.013** | **0.006** | **86** | **2.242** | **0.028\*** | **0.030** | **0.007** | **88** | **4.242** | **5.47E-05\*\*\*** |
| WJ RF State | **0.039** | **0.008** | **241** | **5.067** | **8.06E-07\*\*\*** | **0.051** | **0.007** | **240** | **7.097** | **1.42E-11\*\*\*** |
| Group: Dys Ctrl | **-0.361** | **0.170** | **82** | **-2.120** | **0.037\*** | -0.199 | 0.205 | 84 | -0.968 | 0.336 |
| Group: Typ Ctrl | 0.327 | 0.233 | 79 | 1.403 | 0.165 | -0.017 | 0.282 | 83 | -0.061 | 0.951 |
| Age | **0.154** | **0.051** | **80** | **3.009** | **0.004\*\*** | 0.056 | 0.062 | 83 | 0.906 | 0.367 |
| Movement (Mean FD) | -0.705 | 0.377 | 308 | -1.870 | 0.062. | -0.580 | 0.368 | 294 | -1.578 | 0.116 |
| Num Good Runs | 0.029 | 0.066 | 284 | 0.446 | 0.656 | 0.011 | 0.063 | 269 | 0.177 | 0.860 |
| Dys Ctrl * WJ RF State | -0.002 | 0.022 | 235 | -0.102 | 0.919 | -0.033 | 0.021 | 236 | -1.600 | 0.111 |
| Typ Ctrl * WJ RF State | **-0.039** | **0.016** | **234** | **-2.424** | **0.016\*** | **-0.063** | **0.015** | **236** | **-4.157** | **4.52E-05\*\*\*** |
| **TOWRE** |  |  |  |  |  |  |  |  |  |  |
| Intercept (Intervention) | **-1.839** | **0.713** | **118** | **-2.581** | **0.011\*** | -2.632 | 0.789 | 111 | -3.334 | 0.001** |
| TOWRE Trait | **0.022** | **0.007** | **87** | **2.971** | **0.004\*\*** | **0.047** | **0.008** | **89** | **5.542** | **0.00E+00\*\*\*** |
| TOWRE State | **0.025** | **0.008** | **239** | **3.112** | **0.002\*\*** | **0.037** | **0.007** | **238** | **5.082** | **0.00E+00\*\*\*** |
| Group: Dys Ctrl | -0.192 | 0.177 | 82 | -1.087 | 0.280 | 0.127 | 0.203 | 84 | 0.624 | 0.534 |
| Group: Typ Ctrl | 0.080 | 0.258 | 86 | 0.311 | 0.756 | -0.436 | 0.295 | 88 | -1.479 | 0.143 |
| Age | **0.138** | **0.052** | **82** | **2.643** | **0.010\*\*** | 0.011 | 0.060 | 84 | 0.184 | 0.854 |
| Movement (Mean FD) | -0.600 | 0.389 | 308 | -1.541 | 0.124 | -0.454 | 0.372 | 296 | -1.221 | 0.223 |
| Num Good Runs | 0.081 | 0.067 | 289 | 1.202 | 0.230 | 0.078 | 0.063 | 273 | 1.242 | 0.215 |
| Dys Ctrl * TOWRE State | -0.035 | 0.022 | 237 | -1.578 | 0.116 | -0.040 | 0.020 | 236 | -1.965 | 0.051. |
| Typ Ctrl * TOWRE State | -0.018 | 0.018 | 234 | -1.034 | 0.302 | -0.026 | 0.016 | 234 | -1.593 | 0.113 |
| **WJ MFF** |  |  |  |  |  |  |  |  |  |  |
| Intercept (Intervention) | -0.670 | 0.722 | 112 | -0.928 | 0.356 | -0.892 | 0.882 | 104 | -1.012 | 0.314 |
| WJ MFF Trait | 0.001 | 0.005 | 77 | 0.166 | 0.868 | 0.010 | 0.006 | 80 | 1.564 | 0.122 |
| WJ MFF State | 0.006 | 0.009 | 244 | 0.700 | 0.484 | 0.004 | 0.009 | 242 | 0.493 | 0.623 |
| Group: Dys Ctrl | -0.344 | 0.180 | 83 | -1.911 | 0.060. | -0.136 | 0.227 | 84 | -0.599 | 0.551 |
| Group: Typ Ctrl | **0.651** | **0.186** | **82** | **3.508** | **1.00E-03\*\*\*** | **0.621** | **0.234** | **83** | **2.649** | **0.010\*\*** |
| Age | **0.187** | **0.052** | **84** | **3.633** | **0.00E+00\*\*\*** | **0.123** | **0.065** | **85** | **1.892** | **0.062.** |
| Movement (Mean FD) | **-0.864** | **0.393** | **313** | **-2.198** | **0.029\*** | **-0.836** | **0.401** | **299** | **-2.087** | **0.038\*** |
| Num Good Runs | 0.084 | 0.068 | 293 | 1.231 | 0.219 | **0.074** | **0.068** | **276** | **1.088** | **0.278** |
| Dys Ctrl * WJ MFF State | -0.013 | 0.018 | 246 | -0.700 | 0.484 | 0.004 | 0.017 | 244 | 0.205 | 0.838 |
| Typ Ctrl * WJ MFF State | 0.009 | 0.018 | 239 | 0.483 | 0.630 | -0.012 | 0.018 | 238 | -0.701 | 0.484 |

**Table 2.**
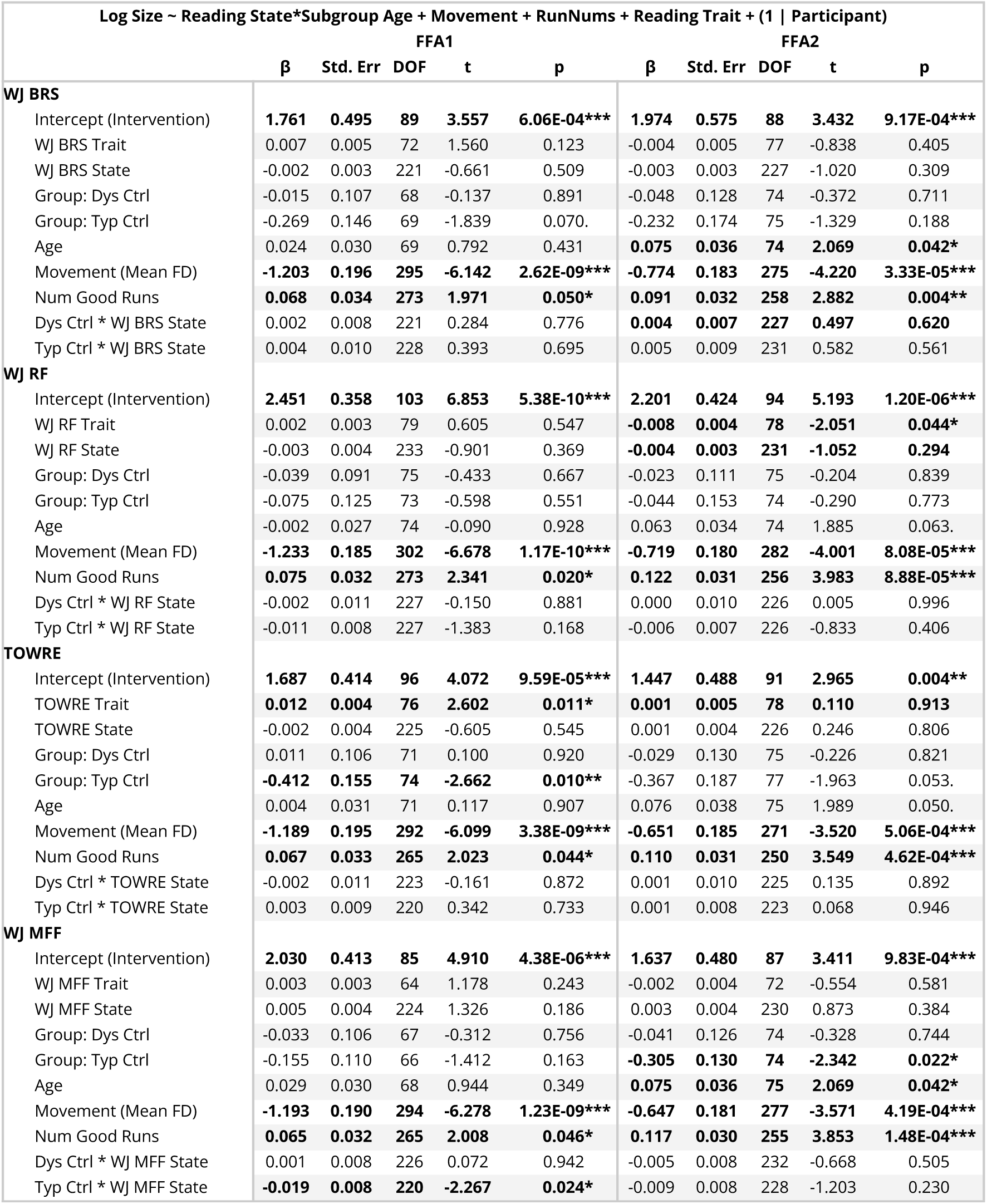
Longitudinal Results for Change in FFA Size with Trait and State Scores. Results of linear mixed effects models (LME) calculating the size (log transformed number of vertices) of a region of interest (ROI) as a function of assessment trait and the interaction between assessment state and participant group with added covariates of participant age, mean framewise displacement during the scan (mean FD), and the number of usable runs of the experiment along with a random intercept by participant. Intervention group is treated as the reference category. Trait: within-participant average over time; State: within-participant change relative to trait; FFA: Fusiform Face Area; WJ BRS: Woodcock-Johnson Basic Reading Skills; WJ RF: Woodcock-Johnson Reading Fluency; TOWRE: Test of Word Reading Efficiency; WJ MFF: Woodcock-Johnson Math Facts Fluency; Dys Ctrl: Dyslexic Control; Typ Ctrl: Typical Control; FD: Framewise Displacement Significant results are displayed in bold and asterisks indicate the degree of significance (p < 0.001: ***, p < 0.01: **, p < 0.05: *, p < 0.1: .). Significant results of p<0.05 are indicated with bold font. Note: No corrections for multiple comparisons were made due to the use of small, manually-defined ROIs. Source data are provided in a public data repository.

**Table 3.** Participant Information and Behavioral Results at First Visit. Demographic information, behavioral assessment scores, and functional MRI task accuracy for participants divided by study group. Gender and multilingual status are collected as self-report responses. WJ BRS: Woodcock-Johnson Basic Reading Skills; WJ RF: Woodcock-Johnson Reading Fluency; TOWRE: Test of Word Reading Efficiency; WJ MFF: Woodcock-Johnson Math Facts Fluency; WASI: Wechsler Abbreviated Scale of Intelligence.

| Participant Information and Scores at First Visit |  |  |  |
| --- | --- | --- | --- |
|  | Control Group |  | Intervention Group |
|  | *Dyslexic* | *Typical* |  |
| N | 19 | 24 | 44 |
| Gender (f/m) | 9 / 10 | 13 / 11 | 20 / 23 |
| Mean age ± SD | 10.2 ± 1.8 | 9.9 ± 1.4 | 9.8 ± 1.1 |
| Multilingual Count | 7 | 11 | 14 |
| **Behavioral Assessments** |  |  |  |
| mean score ± SD | | | |
| WJ BRS | 81.0 ± 10.5 | 110.1 ± 8.5 | 80.4 ± 11.4 |
| WJ RF | 76.5 ± 12.0 | 103.7 ± 9.9 | 73.8 ± 13.9 |
| TOWRE | 71.4 ± 9.8 | 106.0 ± 11.3 | 71.8 ± 10.1 |
| WJ MFF | 79.3 ± 9.0 | 101.4 ± 13.0 | 82.0 ± 15.1 |
| WASI* | 104.8 ± 10.4 | 116.6 ± 10.3 | 101.6 ± 10.8 |
| **fMRI Task Accuracy** |  |  |  |
| mean ± SD | | | |
| hit rate | 0.562 ± 0.203 | 0.649 ± 0.211 | 0.592 ± 0.205 |
| $d'$ | 2.451 ± 0.617 | 2.736 ± 0.718 | 2.555 ± 0.660 |

Finally, to extend our investigation of the relationship between VWFA size and dyslexia, we ran an additional analysis to determine what would be the expected VWFA size given an intervention dosage theoretically sufficient to bring a child’s scores to typical reading levels. This analysis included two steps: we first calculated what is the predicted amount of intervention needed to drive an individual child with dyslexia to achieve reading ability comparable to typical readers in our sample. Then, we examined what the projected VWFA size would be given this duration of intervention. For this analysis we assume a linear effect of intervention duration on reading improvement, based on previous findings^34^. To do this, we first calculate the average WJ BRS score and the average VWFA size for the typical control group at the pre-intervention time point. We then fit a LME to all intervention participants looking at WJ BRS score as a function of time with a random intercept of participant, using data from the pre- and post-intervention time points. Using the result coefficient for time (*β* = 0.101, *p* =1.32E-05, SE = 0.023, %CI = (0.056, 0.147)) and the average typical and average dyslexic BRS scores, we calculated the predicted time (in number of days of intervention) needed for a dyslexic participant to increase their scores to match the average typical reader’s BRS score. In doing so, we found that dyslexic participants would need 265 days of intervention to reach the average score of typical readers - much longer than the 40 days offered in this study. Finally, we fit a second LME to both the pre- and post-intervention time points and the pre-intervention and 1 year follow-up timepoints separately for the dyslexic readers looking at VWFA size as a function of days of intervention. Using these results, we can see that, even with the appropriate amount of intervention to close the gap between dyslexic and typical reader assessment scores, dyslexic readers are still expected to have a smaller VWFA compared with the typical readers from our sample (Figure 6).

**Figure 6.**
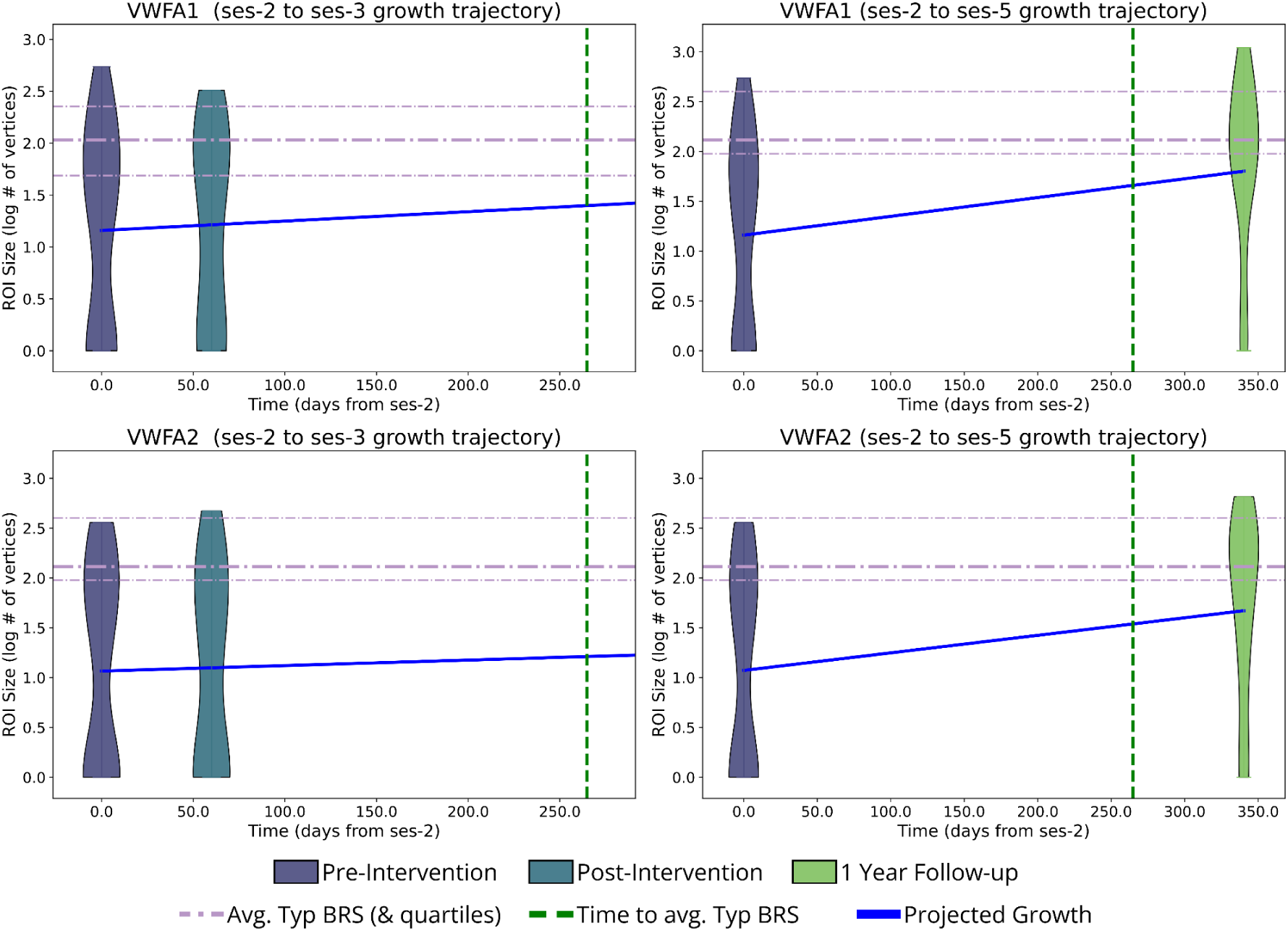
Predicted Visual Word Form Area Size after Sufficient Intervention. Predicted growth of Visual Word Form Area (VWFA) 1 and 2 over time with sufficient time of intervention (265 days; dashed green line) to increase Basic Reading Skills (BRS) scores to the average typical control BRS score assuming linear growth. Violins show the distribution of sizes (in log transformed number of vertices) of VWFAs in the intervention group, in the pre- and post-intervention time points (top) and in the pre-intervention and 1-year follow-up time points (bottom). Dashed purple lines represent the average size (and 25th and 75th percentiles) of a VWFA in the typical control group. Blue lines show the expected growth in VWFA size given a linear estimate of change from the pre-intervention time point to both the post-intervention and 1-year follow-up time points. Source data are provided in a public data repository.

We used a linear prediction of change in VWFA size from ses-2 to ses-3 and a linear prediction of change in VWFA size from ses-2 to ses-5 because, as evident from Figure 3c, there is variability in the rate of ROI growth. By looking at both rates of change, we ensure that the finding holds across multiple trajectories. We see that both the shallow (ses-2 to ses-3 change) and steep (ses-2 to ses-5 change) slopes for predicted growth still do not intersect with the expected typical reader VWFA size even after the hypothetical 265 days of instruction (while ses-5 here represents on average 337 (sd = 14.369) days from the intervention onset at ses-2, only 40 of these were intervention days (days from ses-2 until ses-3), and the rest were not). Specifically, we see that projected VWFAs remain smaller in the dyslexic children using the estimates of change from pre-intervention to post-intervention (VWFA1: 1.399 log # of vertices, VWFA2: 1.212 log # of vertices) and from pre-intervention to 1-year follow-up (VWFA1: 1.660 log # of vertices, VWFA2: 1.539 log # of vertices), compared to the typical readers (VWFA1: 2.0296 log # of vertices, VWFA2: 2.114 log # of vertices). Ultimately, this suggests that even with the appropriate amount of intervention to narrow the gap between dyslexic and typical reader assessment scores, dyslexic readers are still expected to have a smaller VWFA than typical readers.

## Discussion

The balance between plasticity and stability in the developing brain is a key question across spatial scales ranging from molecules^35^, to cells^36^, to circuits^37^, to human perception and cognition^20^. In humans, dyslexia research in particular has long debated which components of the brain’s reading circuitry are plastic and dynamically change with learning, and which (if any) are stable traits that differentiate individuals with dyslexia even after reading improvement. We use the term “state” to refer to aspects of neural responses that are tightly linked to changes in behavior. We use the term “trait” to refer to aspects of neural responses that remain different among participants of different reading levels (e.g. dyslexic versus control) even as their reading skills improve. By combining an intervention design with longitudinal measurements we are able to determine properties of the reading circuitry that are plastic and amenable to change versus those that remain relatively stable throughout the year even with learning. Here, we found both plasticity and stability in the Visual Word Form Area (VWFA). On the one hand, the size and neural tuning properties of the VWFA did change with learning. On the other hand, dramatic differences remained between individuals with dyslexia and typical readers after the intervention, and are projected to remain even if the intervention were extended in length. Based on these findings, we conclude that differences in the VWFA are likely one of the persistent traits that underlie the continual challenges many children with dyslexia face.

We combined several techniques to measure the nature of visual word form-evoked responses in left ventral occipitotemporal cortex (VOTC) in a longitudinal sample of control and intervention participants.The results mentioned are all necessarily related but importantly shed light on various functional properties of VWFA. The current findings highlight significant differences in the existence, size, and functional tuning properties of the VWFA between dyslexic and typical readers. We created a variety of different categories of text stimuli in order to ensure that our experiment was driving responses in both VWFA-1 and VWFA-2. We didn’t find dramatic differences in the stimulus tuning of these regions and this is an important follow-up point for future research. Consistent with previous cross-sectional research, our results confirm that dyslexic readers are less likely to have a detectable VWFA and that this region is smaller compared to their typical peers^9–11^. The reduced detection rates combined with our novel finding of a smaller VWFA size suggest that this region is less specialized for text processing in children with reading difficulties. This may reflect less specialization and efficiency in processing visual word forms, supporting the hypothesis that dyslexia is associated with underdeveloped neural substrates critical for fluent reading. Lastly, the positive correlation between VWFA size and reading ability, across all three reading assessments used here, underscores the functional importance of these regions in supporting literacy. This relationship suggests that a more developed VWFA, as reflected by larger ROI sizes, is associated with better reading performance, potentially due to greater neural resources being allocated to process written language. Importantly, we found that these differences persisted even after intervention, indicating that VWFA size and tuning are potential underlying traits of dyslexia that persist at least a full year following targeted intervention. Admittedly, this finding rests on an assumption that the growth trajectory of VWFA is linear, as is seen with behavioral changes in reading ability^34^. While future studies, which take place over a larger timescale, are needed to confirm the linearity of the VWFA growth, our findings do show that the rate of change in VWFA size is less than the rate of change in reading ability. However, it’s important to note that this effect might be influenced by remaining group differences in reading ability. Non-linear models relating the dynamics VWFA and reading growth could illuminate new findings that make us reconsider the nuanced relationship between the development of the brain’s reading circuitry and the establishment of reading skills. This novel finding suggests that abnormalities in VWFA are enduring characteristics, which likely contribute to the persistent challenges with automating word recognition observed in dyslexic individuals despite improvements in behavioral measures.

Tuning properties of the VWFA also appear to be related to changes in reading ability. As demonstrated in previous studies^11^, text selectivity, or the degree to which response to text is higher than responses to other visual categories, is highly correlated with reading ability. However, previous research has had mixed findings on whether this relationship is due to general hypoactivation of the VWFA^13,14^ as opposed to elevated responses to other categories for dyslexic readers relative to typical readers^11,38^. Our results suggest that the differences in tuning properties between typical and dyslexic readers may be due to a combination of both aspects. Specifically, our percent signal change and selectivity index analyses revealed weaker text-evoked activation in VWFA for dyslexic participants compared to their typical peers, in parallel with a stronger response to visual objects. Crucially, these functional differences were not widespread throughout the entire VOTC as previous research suggested^14^. Rather, this response was specific to the VWFA and was not observed in the adjacent Fusiform Face Area (FFA). This further emphasizes the specialized role of the VWFA in reading.

Our longitudinal analyses shed further light on the nature of change in VWFA driven by improved reading ability. As expected, the intervention successfully improved reading ability across several assessments, consistent with previous studies^34^. Several longitudinal studies have shown that VWFA emerges and develops as children learn to read through the school years^39–41^. Here, we found that this process can be accelerated by an intensive intervention over the course of several weeks. Our intervention design also allows us to disentangle the effects of learning to read from typical brain development that occurs with age. The increased detectability of VWFA-2 in the intervention group and not the dyslexic control group is particularly notable, as it suggests that learning to read can uniquely stimulate the development of critical brain regions that may be underdeveloped in dyslexic readers. This finding is encouraging, as it demonstrates the brain’s plasticity and its ability to adapt in response to a targeted educational environment.

In spite of significant changes in VWFA characteristics following the intervention, our findings suggest that the gap in VWFA size for dyslexic readers compared to typical readers remains a year later. Though differing analytical approaches showed contradictory results about persistent size differences within VWFA-2, differences within VWFA-1 remained regardless of analytical approach. Furthermore, the trait by state analysis revealed that, regardless of group assignment or intervention, size of the VWFA is still highly related to reading ability, suggesting potential constraints on the extent of neural plasticity in this region. Both these findings imply differences in the capacity for plasticity in the VWFA for people with dyslexia. On the other hand, over the timescale of a year differences continue to ameliorate. This leaves open the possibility that there are differences in the developmental time course that will play out in unanticipated ways over longer timescales. Additionally, typical readers continued to show greater text-evoked percent signal change in VWFA-1 compared to intervention participants. Additionally, the elevated object sensitivity observed in dyslexic readers persisted post-intervention, indicating that this patch of cortex continues to respond broadly to different visual categories with less specificity for words. While text selectivity increased for intervention participants in VWFA-2, it did not show a significant improvement in VWFA-1. These persistent differences raise important questions about the limitations of current interventions and the potential need for additional or alternative strategies.

Together, these findings offer a nuanced view of brain plasticity, where intervention can partially improve VWFA function but may not fully “normalize” it. One might ask how this relates to the argument of etiology underlying dyslexia and associated endophenotypes associated with reading difficulties. Though we utilize a variety of methods to measure characteristics of the VWFA that persist despite improved reading ability, we acknowledge that the VWFA is not the only neural system related to reading difficulties. For example, the additive risk factor model from our previous work suggest multiple independent mechanisms spanning visual and language regions^42^, and the new definition of dyslexia proposed by Catts and colleagues suggests a combination of multiple risk and protective mechanisms^43^. Furthermore, it is important to note that our findings are correlational in nature and not causal. We observe that differences in VWFA are related to reading difficulty, but we have not tested the casual relationship of this region. Future research utilizing a carefully monitored, randomized control trial over a larger time-scale is needed to determine causal relationships between the neural and behavioral markers of dyslexia. Absent a casual relationship, our findings do show that aspects of VWFA (like size) are closely tied to one’s reading ability. This supports the notion that while targeted interventions can drive functional changes in the dyslexic brain, some differences may be inherently stable traits that require prolonged or more intensive interventions than possible during a single summer in order to normalize. The dual findings of plasticity and stability imply that VWFA characteristics are influenced by both inherent, developmental factors and responsive, experience-driven changes.

This study is the first to closely compare intervention and control groups across multiple time points, delineating the trajectory of change within one year. Our findings of long lasting changes in the VWFA well after the completion of the intervention emphasize the potential for long-term enhancement through specific, targeted educational programs. The changes observed indicate that particular learning experiences can bring about measurable plasticity, while the stable traits reveal constraints on the plasticity of higher level visual cortex following short-term intervention. Future research should build on these findings by conducting longitudinal analyses across diverse populations and extended timeframes, further investigating the neural mechanisms underlying these functional changes. More specifically, future studies may want to narrow in on individual differences that contribute to a person’s response to intervention, both behaviorally and neurally, while carefully monitoring the kinds and amount of instruction available to each child in order to deepen understanding of causal relationships that are not captured in the current study. While past work has revealed that several factors are related to reading challenges ^44^, a deeper understanding of the heterogeneity of dyslexia is needed for clarity on casual relationships. These insights will be instrumental in understanding effective educational programs and designing interventions to support children with dyslexia. Furthermore, we created a variety of different categories of text stimuli in order to ensure that our experiment was driving responses in both VWFA-1 and VWFA-2. We didn’t find dramatic differences in the stimulus tuning of these regions and this is an important follow-up point for future research.

This study is not without it’s limitations. Firstly, we want to acknowledge that the lack of a randomized control trial limits the extent of the causal relationships that can be made from this data.We made special efforts to make the intervention group as diverse as possible to make our findings more generalizable to populations that are typically underrepresented in research. However, this was difficult to obtain with the controls where there was no benefit to participating in the study. As a result, participant groups are not even matched across all demographic backgrounds. Furthermore, the lack of long term follow ups in several of our participants due to the timing and budget constraints resulting from the COVID-19 pandemic weaken our longitudinal findings. This is especially important as one considers that there may be more nuanced explanations for the longitudinal trajectory of some of our findings (such as with text-selectivity) that may display different dynamics over a larger time scale. Additionally, imbalanced groups affect the statistical power of our analyses and participant attrition could potentially lead to bias in the results. However, we found that the amount of usable data was relatively consistent across groups. We calculated a maximum possible number of runs per group (if each person completed 4 runs of the localizer for each timepoint they were recruited for) and we calculated the total number of usable runs. We found that the intervention group had 80.5% of the maximum possible runs, the dyslexic controls had 73.9% of the maximum possible runs, and the typical controls had 74.7% of the maximum total runs after excluding for things like motion, sleep, and other artifact-related issues. Though we have taken care to exclude data with motion artifacts and control for motion in our statistical analyses, it is important to acknowledge that the effects of motion on fMRI data are complex and could, potentially, have unanticipated effects on the results. We also recognize that, given the imperfect spatial resolution and signal to noise ration, size and response measurements within an ROI might not be measured independently of one another but are instead necessarily related. Finally, though our findings suggest stability in some VWFA traits found in dyslexic readers, extended experimental designs are needed to determine if these differences persist throughout development or in spite of complete behavioral remediation.

In conclusion, we provide robust evidence that dyslexic readers exhibit significant reductions in VWFA size and specialization compared to typical readers, and that targeted interventions can partially mitigate these differences. However, the persistence of some disparities after one year demonstrates a neural characteristic of dyslexia that may remain despite behavioral evidence of remediated reading ability, suggesting the rate of plasticity in this region might explain the enduring challenges that some people with dyslexia experience. This is not to say that interventions are not effective - indeed the behavioral results clearly demonstrate meaningful growth during and after the intervention - but many people with dyslexia continue to have struggles with reading fluency. Our results demonstrate that this patch of cortex is plastic in the dyslexic brain and changes with learning, but that rates of change are slower in those with dyslexia and might be related to continued challenges. This highlights the need for ongoing research to optimize intervention strategies and to explore the full potential of neuroplasticity in supporting literacy development in children with dyslexia. Causality is always a challenge in human neuroscience and intervention studies are few and far between. Though not a randomized controlled trial, the longitudinal intervention design provides unique insights into the balance between stability and plasticity in the developing brain, furthering our understanding of the neurobiology of dyslexia, and offering broader insights into the interplay between brain development and learning.

## Methods

All study protocols were approved by the Stanford School of Medicine Institutional Review Board, and informed assent from the children and written consent from their guardians were obtained prior to participation.

### Participants

A total of 90 participants enrolled in the study. Of these 90 enrolled participants, 12 control participants were included in the study as part of an extension to the original grant to compensate for COIVD-19 delays. Three participants dropped out of the study before any usable fMRI data could be collected. Our final sample for analysis included 87 participants that had at least one usable run of the functional localizer at any given time point of the study. Participants were divided into two groups - an intervention group and a control group. The intervention group consisted of 44 participants who we expected to see significant changes in reading scores throughout the course of the study and the control group consisted of 43 participants (19 participants with dyslexia and 24 participants who were considered typical readers), in whom we did not expect to see significant changes. By including both typical and dyslexic children in the control group, we aimed to see how developmental trajectories may differ between not only the intervention and control participants, but also control participants with varying levels of reading abilities. See Table 2 for a breakdown of age, gender, and scores by participant group for the final study sample. See Table S18 for a breakdown of additional demographic information.

All participants had normal or corrected-to-normal vision, no neurological or hearing issues, and were either monolingual English speakers or used English for at least 60% of their day, having learned the language before age three.

Participants were screened in a 45 minute virtual meeting. Participants were screened using the Wechsler Abbreviated Scale of Intelligence, Second Edition (WASI)^45^, ensuring scores above the 14th percentile for their age group. Exclusions were made for two intervention participants who scored slightly lower (79 compared with 84), where the experimenter’s assessment was that their performance was not representative of their true ability. Participants who scored below a standard score of 85 on either the Woodcock-Johnson Basic Reading Skills (WJ BRS)^46^ or the Test of Word Reading Efficiency (TOWRE)^47^ were classified as “dyslexic” for the purposes of this study (either as an intervention or a control participant). Four participants were not available for a pre-study screening. All four participants had no parent reported reading difficulties and were classified as typical controls based on assessment data from their first study visit; all participants scored over 100 on TOWRE.

Participants were compensated with a flat rate of $120 for every time-point they completed for the study. The intervention was also provided free of charge for all participants who met study criteria for a dyslexia designation. Gender were not considered in the study design as there were no hypotheses regarding gender differences and no gender-specific analyses were performed in the present study. Additionally, gender information was collected from a parent report survey.

### Study Timeline

Participants completed several visits over a 13 month period. Participants completed a baseline visit before the start of the intervention period (or at the beginning of summer for control participants). Eight weeks later and after they finished the intervention, participants returned for their first follow-up visit. Two additional follow-up visits were completed approximately 6 months and 1 year after the start of the intervention. Due to funding and timing constraints associated with the 12 participants enrolled in the study through the extension year, only pre- and post-intervention time points were collected without longer-term follow-up. Intervention group participants also completed an additional pre-baseline visit that occurred one to two months before the baseline visit. See Supplementary Figure S4 for more detail. Behavioral assessments for each time point were collected no more than 2 weeks after the in-person scanning visit.

### Reading Intervention

Participants in the intervention group completed 160 hours of the Seeing Stars: Symbol Imagery for Fluency, Orthography, Sight Words, and Spelling program ^48^ - a curriculum from Lindamood-Bell that has been extensively studied and shown to improve reading ability in children with dyslexia ^34,49,50^. The intervention was delivered for 4 hours a day, 5 days a week, over 8 weeks of the participants’ summer vacation from school. Due to the COVID-19 pandemic, the sessions were conducted remotely via Zoom. All sessions were led by certified Lindamood-Bell instructors. The individualized, multisensory curriculum focused on phonological and orthographic skills, progressing from letters to connected text. Children were guided through activities like air-writing and visualizing letter-sound connections to build literacy foundations. The program emphasized decoding, spelling, fluency, and comprehension. All participants received the intervention at no cost. Dyslexic control group participants were offered free access to the same intervention the summer after their participation in the study.

### Behavioral Assessments

Each visit included a thorough assessment of reading and cognitive skills by trained researchers at Stanford University. The repeated tests at each visit included the Woodcock-Johnson IV (WJ) and Test of Word Reading Efficiency-2 (TOWRE). WJ subtests were used to create Basic Reading Skills (BRS; Letter-Word Identification - LWID, Word Attack - WA) and Reading Fluency (RF; Oral Reading - OR, Sentence Reading Fluency - SRF) composite scores ^46^. The TOWRE tests for Sight-Word Efficiency (SWE) and Phonemic Decoding Efficiency (PDE) were used to calculate the TOWRE ^47^. Additionally, the WJ Math Facts Fluency (MFF) subtest was administered. Alternative test forms were used across visits to ensure reliability and mitigate practice effects. All assessments, except for SRF and MFF, were administered remotely via video call to reduce face-to-face contact between participants and researchers as the study began during the COVID-19 pandemic. Assessment video call sessions were video and audio recorded and all assessments were independently scored by 2 or more trained researchers.

### fMRI Experimental Design

The functional localizer experiment used in the study was an adaptation of White and colleagues’ (2023)^31^ functional localizer experiment. For this experiment, participants viewed 5 main categories of visual stimuli: text, pseudo fonts, objects, faces, and limbs (Figure 1). Each category was composed of more specific subcategories. Text contained high and low frequency words, pseudo words, and consonant strings; pseudo fonts was composed of pseudo fonts designed to contain similar visual features to the Sloan and Courier fonts; objects contained images of random objects like fruits, construction machinery, instruments, and sunglasses; faces contained images of male and female faces facing forward and to either side; and limbs contained images of disembodied hands, arms, feet, and legs. Text stimuli were selected to span every level of semantic familiarity for participants with the intention of tapping elucidating the nuanced functional properties of VWFA-1 and −2. Stimuli were presented in groups of three (all from the same subcategory) on a single frame with a large image on either side of fixation and a smaller image at the center of the screen under the fixation dot (See Figure 1). Size of stimuli were set to balance the visual acuity and processing advantage of fovea and periphery such that they would use comparable amounts of cortical territory.

A single trial consisted of 4 frames of triplets of the same category. Each frame was presented for 800 ms, followed by a 200 ms blank fixation screen. Each subcategory was displayed 5 times during a run and category order was randomly assigned. Additionally, 5 blocks of blank screen trials were randomly dispersed during a run and every run began and ended with a 5 second blank screen.

During each run of the experiment, participants were asked to perform a task; a one-back image repetition task or a fixation color change task. During the one-back task, participants were instructed to press a button every time the image on the screen repeated. During the fixation task, participants were instructed to press a button every time the fixation dot changed colors. Participants used the index finger of their dominant hand to respond. Fixation and one-back targets occurred randomly during 33% of trials in every run, regardless of the instructions provided to the participant.

The experiment was coded in MATLAB (version 2021a) using the Psychloolbox package (version 3.0.17). Participants were monitored via a web camera to ensure they were awake and attending to the task. If a participant fell asleep during the run, the run was stopped and excluded from analyses. Participants completed 4 runs of the experiment (2 of each task) at each visit of the study.

### MRI Acquisition

Data for this study were collected as part of a large longitudinal project which included multiple MRI modalities. In each visit, participants completed the functional localizer reported here, in addition to resting-state functional scans, diffusion scans and quantitative structural scans reported elsewhere^51^. The total scan time was broken down to two sessions to allow participants to rest. Participants were scanned using a General Electric Sigma MR750 3T scanner at Stanford University’s Center for Cognitive and Neurobiological Imaging (CNI). Before the main scanning session, participants attended an introductory session where they practiced a brief version of the experiment in a mock scanner. This session helped them get accustomed to the scanner noises, tasks, and response box. They also practiced staying still with the aid of the MoTrak Head Motion Tracking System (RRID:SCR_009607), which provided feedback on their motion.

Functional runs were collected using a gradient echo EPI sequence with a multiband factor of 3, ensuring whole-brain coverage across 51 slices. The acquisition parameters included a TR of 1.19s, a TE of 30ms, and a flip angle of 62, resulting in a spatial resolution of 2.4 mm³ isotropic voxels. Each run consisted of 232 frames and lasted 4 minutes and 36 seconds. Additionally, a high-resolution T1-weighted anatomical scan was acquired with a spatial resolution of 0.9 mm³ isotropic voxels.

To maximize data quality and minimize potential confounds, research coordinators are trained to monitor 1) motion in real time based on FIRMM, 2) participant engagement and focus based on a webcam focused on the participant’s eyes while they are in the scanner, 3) responses during the task, 4) potential artifacts in the fMRI data. If any data quality issues are detected (e.g., participant closing eyes, moving, etc.) the scan is ended early and researchers communicate with the participant to ensure attentiveness before restarting the scan.

### MR Proprocessing

Functional data preprocessing was carried out using fMRIPrep version 23.1.3 ^52^, which is built upon Nipype version 1.8.6 ^53,54^.

#### Anatomical Data Preprocessing

The T1-weighted (T1w) image was first reoriented to align with the AC-PC axis using ANTSpy version 0.4.2 ^55^ and served as the anatomical reference throughout the preprocessing workflow. The T1w image was then skull-stripped with Synthstrip ^56^ and processed through FreeSurfer’s recon-all pipeline (version 7.3.2; ^57^) using the Synthseg robust algorithm ^58^ for segmentation and surface reconstruction. All surfaces underwent visual inspection, with manual edits made where necessary. The resulting FreeSurfer derivatives were subsequently utilized by the fMRIPrep pipeline. Segmentation of brain tissue into cerebrospinal fluid (CSF), white matter (WM), and gray matter (GM) was performed on the skull-stripped T1w image using FAST (FSL, RRID; ^59^). Spatial normalization to a standard space (MNI152NLin2009cAsym) was achieved through nonlinear registration with antsRegistration (ANTs; Tustison et al.), using skull-stripped versions of both the T1w reference and the T1w template.

#### Functional Data Preprocessing

For each BOLD run, a reference volume was created using a custom method implemented in fMRIPrep. Before any spatiotemporal filtering, head-motion parameters relative to the BOLD reference (including transformation matrices and six rotation and translation parameters) were estimated using mcflirt (FSL; ^60^). A **B0** nonuniformity map was estimated from two echo-planar imaging (EPI) references using topup ^61^, and the estimated fieldmap was rigidly aligned to the target EPI reference run. The field coefficients were then applied to the reference EPI using the appropriate transformation. Slice-time correction was performed on the BOLD runs to align all slices to the middle slice using 3dTshift from AFNI (^62^; RRID). The BOLD reference was co-registered to the T1w reference using bbregister (FreeSurfer), which implements boundary-based registration^63^. Framewise motion parameters and BOLD signal within the white matter (WM) and corticospinal fluid (CSF) were calculated and later used as confound regressors in the BOLD response estimation. The BOLD time-series were resampled into standard space, resulting in preprocessed BOLD runs in MNI152NLin2009cAsym space, and were also resampled onto the fsnative and fsaverage FreeSurfer surfaces. All resampling steps were executed with a single interpolation process, combining all necessary transformations (e.g., head-motion correction, susceptibility distortion correction, and co-registration to anatomical and output spaces). Volumetric resampling was conducted using antsApplyTransforms (ANTs) with Lanczos interpolation to reduce smoothing effects, while surface resampling was performed using mri_vol2surf (FreeSurfer).

#### BOLD Response Estimation

We used mriqc (version 22.0.1;^64^) to assess data quality and exclude noisy runs. Runs were excluded from analysis if the mean framewise displacement (FD) was 0.5 mm or larger, or if more than 30% of frames had an FD greater than 0.5 mm. Additionally, runs where participants failed to keep their eyes open were also excluded. BOLD response estimation was carried out by fitting a general linear model (GLM) to the BOLD time-series in each participant’s native surface using Nilearn (version 0.5.0). The design matrix for the GLM included signals from the white matter and CSF, along with their first derivatives, as well as the six motion parameters. First- and second-order polynomial drift terms were also included. The estimated BOLD responses (beta weights) are reported in units of percent signal change, reflecting the change in response relative to blank trials where participants viewed a blank fixation screen.

### Statistical Analysis

All analyses reported in this study involved 2-sided statistical tests.

#### Response Estimation

Activation maps for neural responses to the visual localizer were created by fitting general linear models convolving the SPM hemodynamic response function to stimulus onsets using the nilearn package (version 0.10.4) in python^65,66^ (version 3.11.8). This approach was used to create two different types of heat maps. First, statistical maps displaying response preference for one visual category compared to others were created to define functional ROIs. For defining VWFAs, this was done by contrasting the response to text (high frequency and low frequency words, consonant strings, and pseudo words) to response to all other visual categories (faces, pseudo fonts, objects, and limbs). A similar process was done for defining FFAs with contrasts of faces to other visual categories. Resulting maps were reported as t-statistics for every vertex on the cortical surface. For analyzing response profiles within ROIs, heat maps were created for each of the visual categories with values of percent signal change from baseline. These values were averaged across all vertices in each ROI.

#### Linear Mixed-Effects Modeling

We used the lme4 package (version 1.1.31) in R^67^ (version 2022.12.0) to run linear mixed effects models (LMEs) to investigate differences in baseline percent signal change, and all longitudinal changes in assessment scores, size, signal, and selectivity. All LMEs that utilized neuroimaging data also included covariates of participant age at baseline, along with the mean framewise displacement and the number of usable runs per session of data collection. Every time an LME included a group level parameter and interaction, the intervention group was set as the reference category. In baseline analyses, the dyslexic group was set as the reference category. Similarly, every time an LME included an analysis of percent signal change, response to text was set as the reference category.

### Region of Interest Definition

Regions of interest (ROIs) were manually drawn on the native surface of every participant using the Freeview viewer of FreeSurfer Suite^68^. To determine exact boundaries of these ROIs, we calculated statistical maps that compared the estimated BOLD response of a target category (i.e. text) to the other four remaining categories (i.e. pseudo fonts, faces, limbs, objects). Contrasts were weighted by the number of sub categories to ensure balance across the varying number of blocks per stimulus category. Maps were thresholded at t-value > 3 and defined with the following anatomical constraints: All ROIs were defined within the bounds of the ventral occipitotemporal cortex (VOTC); posterior of the anterior tip of the occipitotemporal sulcus (OTS), lateral to the collateral sulcus, anterior to the posterior transverse collateral sulcus, and medial to and inclusive of the OTS. VWFA clusters were defined using any vertices that met threshold value which fell on the left occipitotemporal sulcus and lateral portion of the fusiform gyrus (any patch of cortex lateral to the mid-fusiform sulcus). See Supplemental Video S1 for a walk-through of specific steps taken to draw a sample VWFA. FFA clusters were defined using any vertices that met threshold value which fell on the fusiform gyrus and mid fusiform sulcus. Continuous clusters of activation that extended further medially or laterally than the described anatomy were bounded by the borders of the described anatomy.

The more posterior VWFA-1 and FFA-1 were discerned from the more anterior VWFA-2^69^ and FFA-2 using the boundaries of the Fusiform Gyrus cytoarchitectonic FG2 & FG4 ROIs from Rosenke et al. ^70^. These template ROIs were projected from average space to the native space of every participant. VWFA-1 and FFA-1 were roughly extended to the anterior boundary FG2 while VWFA-2 and FFA-2 extended from the posterior boundary of FG4 until the anterior tip of the fusiform gyrus. Continuous patches of activation were split at natural saddles in activation that fell closest to these anterior and posterior boundaries. Medial and lateral boundaries of native anatomy were strictly adhered to.

Two sets of ROIs were drawn with these criteria. The first set of ROIs, on which the majority of analysis was conducted, used data from every available run for each participant across all time points to create statistical maps. These “combined session” ROIs were used for all analyses involving percent signal change and category selectivity. With this approach, we were able to define VWFA-1 in 73 participants and VWFA-2 in 66 participants out of the 87 total participants. A second set of ROIs was drawn for each time point of the study using statistical maps that were created from all available runs (Supplementary Figure S5) per study time point. These “separate session” ROIs were used for analyses of VWFA emergence and size. See Supplementary (Figure S6 and Figure S7) for images of all ROIs drawn for this study.

## Data Availability

The data used in this study have been deposited in the Stanford Libraries Digital Repository database under DOI 10.25740/bq006zp5312 [https://purl.stanford.edu/bq006zp5312].

## Code Availability

Code used to generate manuscript display items and statistical analyses can be found in the GitHub repository https://github.com/jamielmitchell/Mitchell_TraitStateVWFA_2025.

## Acknowledgements

This work was funded by NICHD R01-HD095861 to JDY. We would like to thank the children and their families who participated in this study, the research assistants who contributed to data collection, Dr. Alex White for the task and stimulus design, and Lindamood-Bell Learning Center for donating their time and resources to provide the intervention for our participants.

## Author Contributions

JLM: Conceptualization, Formal analysis, Investigation, Writing (original draft & review and editing), Visualization. MY: Investigation,Writing (review and editing), Supervision. HLS: Investigation, Data curation, Project administration. MJ: Investigation, Data curation, Project administration. MET: Investigation, Data curation, Project administration. KAT: Investigation, Data curation, Project administration. JET: Investigation, Data curation, Project administration. CC: Investigation, Data curation, Project administration. JDY: Conceptualization, Resources, Writing (review and editing), Supervision, Funding acquisition.

