## Supplemental Material for "The balance between stability and plasticity of the Visual Word Form Area in dyslexia"

| ROI Size Difference at Baseline | | | | | | |
|---|---|---|---|---|---|---|
| | t (dys > typ) | CI *low* | CI *high* | DOF | p | |
| VWFA1 | **-4.603** | **-1.348** | **-0.534** | **82** | **1.50E-05** | *** |
| VWFA2 | **-4.593** | **-1.491** | **-0.590** | **82** | **1.56E-05** | *** |
| FFA1 | 0.378 | -0.190 | 0.280 | 82 | 0.706 | |
| FFA2 | **2.262** | **0.037** | **0.577** | **82** | **0.026** | * |

**Table S1 | Cross-Sectional Results for ROI Size Difference at Baseline**
Results from a two-sided t-test comparing the size (in log number of vertices) difference between dyslexic participants (n = 59) and typical reader participants (n = 24) for both Visual Word Form Area (VWFA) 1 & 2, and Fusiform Face Area (FFA) 1 & 2. Significant results are displayed in bold and asterisks indicate the degree of significance ($p < 0.001$: *****, $p < 0.01$: ****, $p < 0.05$: ***, $p < 0.1$: .) Significant results of $p<0.05$ are indicated with bold font. Source data are provided in a public data repository.

| ROI Size (>0) Difference at Baseline | | | | | |
|---|---|---|---|---|---|
| | **t (dys > typ)** | **CI** *low* | *high* | **DOF** | **p** |
| VWFA1 | **-3.182** | **-0.650** | **-0.148** | **59** | **0.002 **** |
| VWFA2 | **-3.182** | **-0.642** | **-0.182** | **54** | **7.20E-04 ***** |
| FFA1 | -0.644 | -0.260 | 0.133 | 81 | 0.521 |
| FFA2 | 1.743 | -0.022 | 0.332 | 79 | 0.085 . |

**Table S2 | Cross-Sectional Results for ROI Size (>0) Difference at Baseline**
Results from a two-sided t-test comparing the size (in log number of vertices) difference between dyslexic participants and typical reader participants for both Visual Word Form Area (VWFA) 1 & 2, and Fusiform Face Area (FFA) 1 & 2. Participants were excluded from each analysis if they were missing the corresponding ROI. Significant results are displayed in bold and asterisks indicate the degree of significance (p < 0.001: ***, p < 0.01: **, p < 0.05: *, p < 0.1: .) Significant results of p<0.05 are indicated with bold font. Source data are provided in a public data repository.

| Log Size & Score Correlations | | | | | |
|---|---|---|---|---|---|
| | **r** | **CI** | | **DOF** | **p** |
| | | *low* | *high* | | |
| **VWFA1** | | | | | |
| WJ BRS | **0.579** | **0.416** | **0.706** | **82** | **7.86E-09***** |
| WJ RF | **0.511** | **0.329** | **0.656** | **79** | **1.09E-06***** |
| TOWRE | **0.581** | **0.415** | **0.709** | **79** | **1.32E-08***** |
| WJ MFF | **0.282** | **0.072** | **0.468** | **82** | **0.009**** |
| **VWFA2** | | | | | |
| WJ BRS | **0.573** | **0.408** | **0.701** | **82** | **1.27E-08***** |
| WJ RF | **0.601** | **0.441** | **0.724** | **79** | **2.96E-09***** |
| TOWRE | **0.631** | **0.478** | **0.746** | **79** | **2.76E-10***** |
| WJ MFF | **0.393** | **0.195** | **0.560** | **82** | **2.20E-04***** |
| **FFA1** | | | | | |
| WJ BRS | 0.022 | -0.193 | 0.236 | 82 | 0.840 |
| WJ RF | 0.133 | -0.088 | 0.341 | 79 | 0.238 |
| TOWRE | 0.175 | -0.045 | 0.379 | 79 | 0.118 |
| WJ MFF | 0.055 | -0.162 | 0.266 | 82 | 0.621 |
| **FFA2** | | | | | |
| WJ BRS | -0.169 | -0.370 | 0.047 | 82 | 0.125 |
| WJ RF | -0.135 | -0.343 | 0.086 | 79 | 0.230 |
| TOWRE | -0.051 | -0.266 | 0.170 | 79 | 0.653 |
| WJ MFF | -0.163 | -0.365 | 0.053 | 82 | 0.138 |

**Table S3 | Cross-Sectional Correlation Results for ROI Size and Assessments at Baseline**
Results from two-sided Pearson R tests correlating size of Visual Word Form Area (VWFA) 1 & 2 and Fusiform Face Area (FFA) 1 & 2 to reading and math ability. Size is calculated as the log transformed number of vertices. Woodcock-Johnson Basic Reading Skill score (WJ BRS), Reading Fluency score (WJ RF), and Test of Word Reading Efficiency Index (TOWRE). Significant results are displayed in bold and asterisks indicate the degree of significance ($p < 0.001$: ***, $p < 0.01$: **, $p < 0.05$: *, $p < 0.1$: .) Significant results of $p<0.05$ are indicated with bold font. Source data are provided in a public data repository.

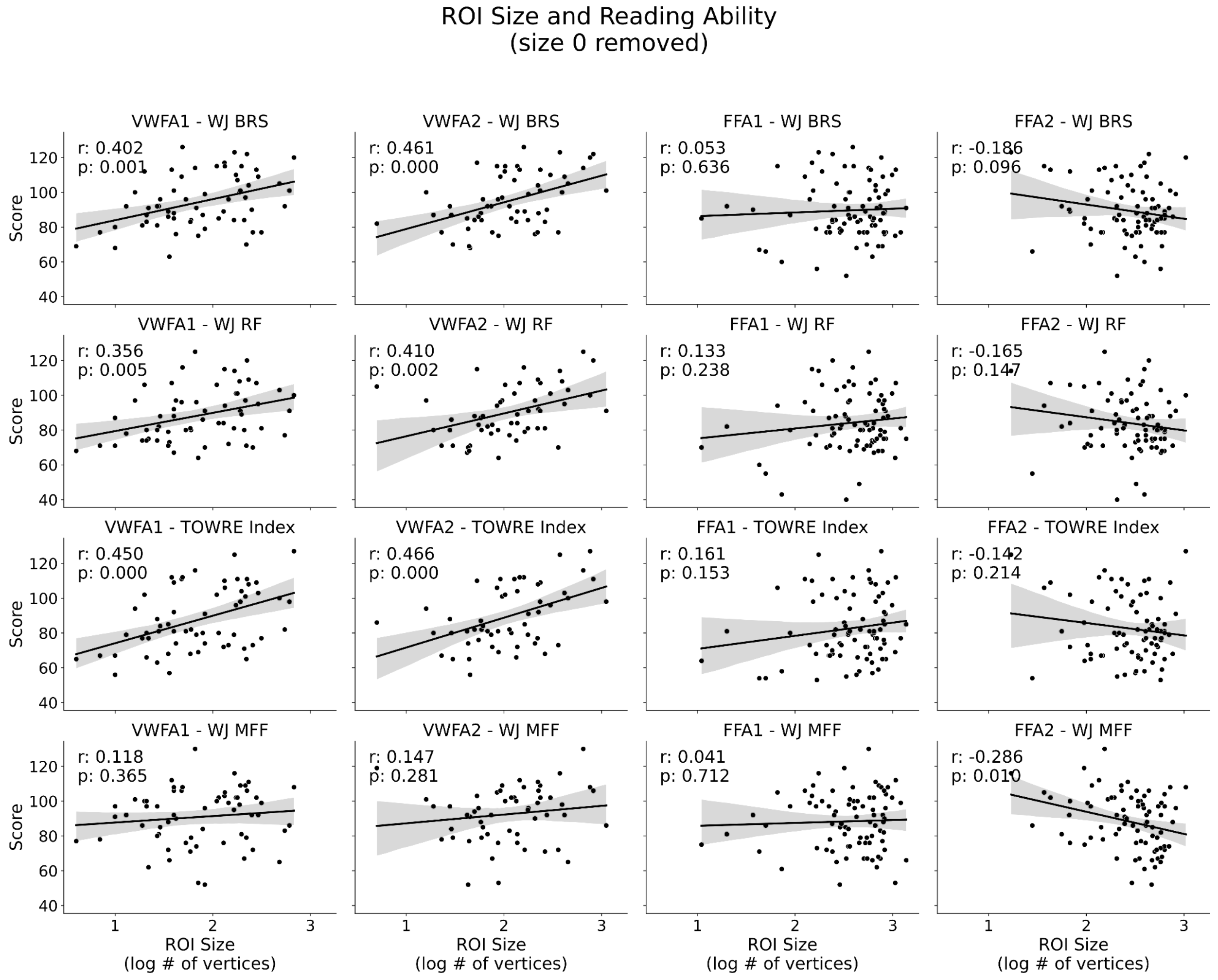


**Figure S1 |Correlation Between Visual Word Form Area Sizes (>0) and Reading Ability**
Visual Word Form Area (VWFA) size is positively correlated with reading ability even when excluding participants with no VWFA (VWFAs with a size of 0 vertices). Size of Fusiform Face Area (FFA) is not related to reading ability. VWFA and FFA size are not related to math ability. Woodcock-Johnson Basic Reading Skill score (WJ BRS), Reading Fluency score (WJ RF), and Test of Word Reading Efficiency Index (TOWRE). Source data are provided in a public data repository.

| Mean % Signal Change ~ Category * Subgroup + Age + Movement + RunNums + (1 \| Participant) | | | | | | | | | | | | | | | | | | | | |
|---|---|---|---|---|---|---|---|---|---|---|---|---|---|---|---|---|---|---|---|---|
| | **VWFA1** | | | | | **VWFA2** | | | | | **FFA1** | | | | | **FFA2** | | | | |
| | **β** | **Std. Err** | **DOF** | **t** | **p** | **β** | **Std. Err** | **DOF** | **t** | **p** | **β** | **Std. Err** | **DOF** | **t** | **p** | **β** | **Std. Err** | **DOF** | **t** | **p** |
| Intercept (Dyslexic Readers \| Text) | 1.126 | 0.927 | 65 | 1.215 | 0.229 | 0.073 | 0.568 | 59 | 0.128 | 0.898 | 0.314 | 0.695 | 79 | 0.452 | 0.652 | **1.243** | **0.402** | **77** | **3.093** | **0.003**** |
| Category: Pseudo Fonts | **-0.198** | **0.059** | **272** | **-3.345** | **9.40E-04***** | **-0.223** | **0.035** | **248** | **-6.329** | **1.15E-09***** | -0.034 | 0.047 | 328 | -0.734 | 0.464 | -0.056 | 0.033 | 320 | -1.716 | 0.087. |
| Category: Objects | **-0.252** | **0.059** | **272** | **-4.258** | **2.84E-05***** | **-0.291** | **0.035** | **248** | **-8.237** | **1.02E-14***** | **1.002** | **0.047** | **328** | **21.356** | **4.88E-64***** | **0.415** | **0.033** | **320** | **12.773** | **1.79E-30***** |
| Category: Faces | **-0.852** | **0.059** | **272** | **-14.369** | **3.22E-35***** | **-0.468** | **0.035** | **248** | **-13.253** | **1.14E-30***** | **1.161** | **0.047** | **328** | **24.738** | **5.59E-77***** | **0.680** | **0.033** | **320** | **20.909** | **8.28E-62***** |
| Category: Limbs | **-0.323** | **0.059** | **272** | **-5.447** | **1.15E-07***** | **-0.275** | **0.035** | **248** | **-7.789** | **1.85E-13***** | **0.895** | **0.047** | **328** | **19.081** | **3.97E-55***** | **0.530** | **0.033** | **320** | **16.315** | **5.74E-44***** |
| Group: Typical Readers | **0.353** | **0.174** | **87** | **2.024** | **0.046*** | **0.213** | **0.105** | **79** | **2.015** | **0.047*** | 0.176 | 0.145 | 108 | 1.215 | 0.227 | 0.009 | 0.087 | 119 | 0.102 | 0.919 |
| Age | **0.119** | **0.058** | **65** | **2.039** | **0.046*** | 0.036 | 0.039 | 59 | 0.933 | 0.355 | 0.071 | 0.047 | 79 | 1.510 | 0.135 | -0.019 | 0.027 | 77 | -0.707 | 0.481 |
| Movement (Mean FD) | -0.791 | 0.699 | 65 | -1.131 | 0.262 | -0.309 | 0.410 | 59 | -0.754 | 0.454 | -0.427 | 0.555 | 79 | -0.769 | 0.444 | **-1.074** | **0.318** | **77** | **-3.382** | **0.001**** |
| # of Good Runs | -0.227 | 0.149 | 65 | -1.524 | 0.132 | 0.016 | 0.086 | 59 | 0.183 | 0.856 | -0.025 | 0.103 | 79 | -0.242 | 0.809 | **-0.128** | **0.062** | **77** | **-2.071** | **0.042*** |
| Category: Pseudo Fonts * Group: Typical Readers | -0.185 | 0.103 | 272 | -1.786 | 0.075. | **-0.135** | **0.062** | **248** | **-2.189** | **0.030*** | -0.088 | 0.088 | 328 | -1.007 | 0.315 | -0.108 | 0.061 | 320 | -1.753 | 0.081. |
| Category: Objects * Group: Typical Readers | **-0.380** | **0.103** | **272** | **-3.673** | **2.88E-04***** | **-0.206** | **0.062** | **248** | **-3.343** | **9.57E-04***** | **-0.275** | **0.088** | **328** | **-3.131** | **0.002**** | **-0.194** | **0.061** | **320** | **-3.154** | **0.002**** |
| Category: Faces * Group: Typical Readers | -0.060 | 0.103 | 272 | -0.577 | 0.565 | **-0.145** | **0.062** | **248** | **-2.360** | **0.019*** | -0.095 | 0.088 | 328 | -1.078 | 0.282 | -0.107 | 0.061 | 320 | -1.742 | 0.082. |
| Category: Limbs * Group: Typical Readers | -0.198 | 0.103 | 272 | -1.915 | 0.057. | -0.060 | 0.062 | 248 | -0.976 | 0.330 | -0.093 | 0.088 | 328 | -1.062 | 0.289 | -0.081 | 0.061 | 320 | -1.312 | 0.190 |

**Table S4 | Cross-Sectional LME Results for the Percent Signal Change at Baseline**

Results from a linear mixed effects model (LME) looking at the interaction between group and response to stimulus category (in units of mean percent signal change) withing Visual Word Form Area (VWFA) 1 & 2 and Fusiform Face Area (FFA) 1 & 2.. Woodcock-Johnson Basic Reading Skill score (WJ BRS), Reading Fluency score (WJ RF), and Test of Word Reading Efficiency Index (TOWRE). Significant results are displayed in bold and asterisks indicate the degree of significance ($p < 0.001$: ***, $p < 0.01$: **, $p < 0.05$: *, $p < 0.1$: .). Significant results of $p<0.05$ are indicated with bold font. Note: No corrections for multiple comparisons were made due to the use of small, manually-defined ROIs. Source data are provided in a public data repository.

| Text Selectivity & Score Correlations | | | | | | |
|---|---|---|---|---|---|---|
| | r | CI | | DOF | p | |
| | | *low* | *high* | | | |
| **VWFA1** | | | | | | |
| WJ BRS | **0.439** | **0.226** | **0.612** | **67** | **1.61E-04** | ******* |
| WJ RF | **0.403** | **0.182** | **0.585** | **66** | **6.53E-04** | ******* |
| TOWRE | **0.356** | **0.125** | **0.551** | **64** | **0.003** | ****** |
| WJ MFF | 0.180 | -0.059 | 0.400 | 67 | 0.139 | |
| **VWFA2** | | | | | | |
| WJ BRS | **0.383** | **0.147** | **0.578** | **60** | **0.002** | ****** |
| WJ RF | **0.400** | **0.165** | **0.592** | **59** | **0.001** | ****** |
| TOWRE | **0.397** | **0.157** | **0.593** | **57** | **0.002** | ****** |
| WJ MFF | 0.145 | -0.109 | 0.381 | 60 | 0.262 | |

**Table S5 | Cross-Sectional Correlations for Selectivity and Assessments at Baseline**
Results from two-sided Pearson R tests correlating the text selectivity of Visual Word Form Area (VWFA) 1 & 2 a to reading and math ability. Selectivity index is calculated as the difference between activation to text versus non text, divided by the sum of activation to all stimuli. Woodcock-Johnson Basic Reading Skill score (WJ BRS), Reading Fluency score (WJ RF), and Test of Word Reading Efficiency Index (TOWRE). Significant results are displayed in bold and asterisks indicate the degree of significance ($p < 0.001$: ***, $p < 0.01$: **, $p < 0.05$: *, $p < 0.1$: .) Significant results of $p<0.05$ are indicated with bold font. Source data are provided in a public data repository.

| | Score ~ Time + (1 \| Participant) | | | | | | | | | | | | | | |
|---|---|---|---|---|---|---|---|---|---|---|---|---|---|---|---|
| | Intervention Group | | | | | Dyslexic Control Group | | | | | Typical Control Group | | | | |
| | β | Std. Err | DOF | t | p | β | Std. Err | DOF | t | p | β | Std. Err | DOF | t | p |
| **WJ BRS** | | | | | | | | | | | | | | | |
| Intercept | **82.678** | **1.412** | **47** | **58.554** | **1.29E-45*****| **79.820** | **2.440** | **20** | **32.718** | **6.02E-19***** | **110.237** | **1.803** | **25** | **61.148** | **4.03E-29***** |
| Time (Days from ses2) | **0.021** | **0.003** | **168** | **7.233** | **1.61E-11***** | -0.004 | 0.006 | 42 | -0.702 | 0.487 | -0.006 | 0.005 | 50 | -1.383 | 0.173 |
| **WJ RF** | | | | | | | | | | | | | | | |
| Intercept | **75.447** | **1.903** | **44** | **39.653** | **2.76E-36***** | **76.248** | **2.771** | **19** | **27.512** | **1.19E-16***** | **105.118** | **2.324** | **23** | **45.229** | **2.40E-24***** |
| Time (Days from ses2) | **0.020** | **0.002** | **163** | **8.037** | **1.77E-13***** | **0.008** | **0.004** | **42** | **2.084** | **0.043*** | **0.020** | **0.005** | **50** | **4.288** | **8.23E-05***** |
| **TOWRE** | | | | | | | | | | | | | | | |
| Intercept | **73.134** | **1.494** | **45** | **48.955** | **1.37E-40***** | **69.856** | **2.060** | **20** | **33.918** | **1.94E-19***** | **103.481** | **2.121** | **24** | **48.799** | **2.66E-25***** |
| Time (Days from ses2) | **0.026** | **0.002** | **165** | **12.251** | **5.74E-25***** | 0.003 | 0.005 | 42 | 0.660 | 0.513 | -0.004 | 0.005 | 46 | -0.711 | 0.481 |
| **WJ MFF** | | | | | | | | | | | | | | | |
| Intercept | **82.707** | **2.364** | **44** | **34.989** | **6.43E-34***** | **77.360** | **2.351** | **22** | **32.908** | **5.84E-20***** | **100.144** | **2.781** | **25** | **36.014** | **8.59E-23***** |
| Time (Days from ses2) | **0.007** | **0.003** | **167** | **2.567** | **0.011*** | 0.000 | 0.006 | 44 | 0.018 | 0.985 | 0.009 | 0.005 | 49 | 1.798 | 0.078. |

**Table S6 | Longitudinal LME Results for Change in Assessment Scores by Group**

Results from a linear mixed effects model (LME) calculating assessment score as a function of time (in days from baseline) with a random intercept by participant for each study group separately. Woodcock-Johnson Basic Reading Skill score (WJ BRS), Reading Fluency score (WJ RF), and Test of Word Reading Efficiency Index (TOWRE). Significant results are displayed in bold and asterisks indicate the degree of significance ($p < 0.001$: ***, $p < 0.01$: **, $p < 0.05$: *, $p < 0.1$: .). Significant results of $p<0.05$ are indicated with bold font. Note: No corrections for multiple comparisons were made due to the use of small, manually-defined ROIs. Source data are provided in a public data repository.

| Score ~ Time * Subgroup + (1 \| Participant) | | | | | |
|---|---|---|---|---|---|
| | **β** | **Std. Err** | **DOF** | **t** | **p** |
| **WJ BRS** | | | | | |
| Intercept \| Intervention Group | **82.678** | **1.400** | **85** | **59.037** | **7.35E-71*** ** |
| Time (Days from ses2) | **0.021** | **0.003** | **256** | **7.605** | **5.44E-13*** ** |
| Group: Dys Ctrl | -2.874 | 2.646 | 98 | -1.086 | 0.280 |
| Group: Typ Ctrl | **27.589** | **2.439** | **97** | **11.311** | **1.95E-19*** ** |
| Group: Dys Ctrl * Time | **-0.025** | **0.006** | **265** | **-3.872** | **1.36E-04*** ** |
| Group: Typ Ctrl * Time | **-0.028** | **0.006** | **268** | **-4.287** | **2.53E-05*** ** |
| **WJ RF** | | | | | |
| Intercept \| Intervention Group | **75.447** | **1.815** | **84** | **41.563** | **9.16E-58*** ** |
| Time (Days from ses2) | **0.020** | **0.002** | **254** | **8.468** | **2.03E-15*** ** |
| Group: Dys Ctrl | 0.799 | 3.352 | 89 | 0.238 | 0.812 |
| Group: Typ Ctrl | **29.667** | **3.137** | **88** | **9.457** | **4.60E-15*** ** |
| Group: Dys Ctrl * Time | **-0.011** | **0.005** | **258** | **-2.109** | **0.036*** |
| Group: Typ Ctrl * Time | 0.001 | 0.005 | 258 | 0.133 | 0.894 |
| **TOWRE** | | | | | |
| Intercept \| Intervention Group | **73.133** | **1.465** | **83** | **49.918** | **6.45E-64*** ** |
| Time (Days from ses2) | **0.026** | **0.002** | **250** | **11.764** | **1.03E-25*** ** |
| Group: Dys Ctrl | -3.271 | 2.732 | 92 | -1.197 | 0.234 |
| Group: Typ Ctrl | **30.311** | **2.516** | **90** | **12.047** | **1.71E-20*** ** |
| Group: Dys Ctrl * Time | **-0.023** | **0.005** | **257** | **-4.328** | **2.15E-05*** ** |
| Group: Typ Ctrl * Time | **-0.029** | **0.005** | **256** | **-5.682** | **3.61E-08*** ** |
| **WJ MFF** | | | | | |
| Intercept \| Intervention Group | **82.709** | **2.112** | **86** | **39.168** | **1.49E-56*** ** |
| Time (Days from ses2) | **0.007** | **0.003** | **258** | **2.579** | **0.010*** |
| Group: Dys Ctrl | -5.341 | 3.900 | 91 | -1.370 | 0.174 |
| Group: Typ Ctrl | **17.437** | **3.601** | **90** | **4.842** | **5.29E-06*** ** |
| Group: Dys Ctrl * Time | -0.007 | 0.006 | 262 | -1.109 | 0.268 |
| Group: Typ Ctrl * Time | 0.003 | 0.006 | 263 | 0.410 | 0.682 |

**Table S7 | Longitudinal LME Results for Change in Assessment Scores with Group Interaction**
Results from a linear mixed effects model (LME) calculating assessment score as a function of the interaction between time (in days from baseline) and participant group with a random intercept by participant. Woodcock-Johnson Basic Reading Skill score (WJ BRS), Reading Fluency score (WJ RF), and Test of Word Reading Efficiency Index (TOWRE). Significant results are displayed in bold and asterisks indicate the degree of significance (p < 0.001: ***, p < 0.01: **, p < 0.05: *, p < 0.1: .). Significant results of p<0.05 are indicated with bold font. Note: No corrections for multiple comparisons were made due to the use of small, manually-defined ROIs. Source data are provided in a public data repository.

| Presence ~ Time + Age + Motion + RunNums + (1 \| Participant) | coef | Std. Err | z | P>\|z\| |
|---|---|---|---|---|
| **VWFA1** | | | | |
| ***Intervention*** | | | | |
| Intercept | -12.347 | 7 | -1.884 | 0.060. |
| Time (Days from ses2) | **0.005** | **0** | **2.277** | **0.023*** |
| Age | **1.295** | **1** | **2.087** | **0.037*** |
| Movement (Mean FD) | -2.245 | 3 | -0.726 | 0.468 |
| Num Good Runs | 0.568 | 1 | 0.984 | 0.325 |
| ***Dyslexic Controls*** | | | | |
| Intercept | -11.322 | 6 | -1.763 | 0.078. |
| Time (Days from ses2) | 0.005 | 0 | 1.218 | 0.223 |
| Age | **1.223** | **1** | **2.196** | **0.028*** |
| Movement (Mean FD) | 2.652 | 4 | 0.619 | 0.536 |
| Num Good Runs | -0.260 | 1 | -0.458 | 0.647 |
| **VWFA2** | | | | |
| ***Intervention*** | | | | |
| Intercept | -15.285 | 9 | -1.619 | 0.106 |
| Time (Days from ses2) | **0.009** | **0** | **3.347** | **8.16E-04***** |
| Age | **1.820** | **1** | **2.011** | **0.044*** |
| Movement (Mean FD) | -8.490 | 5 | -1.616 | 0.106 |
| Num Good Runs | 0.171 | 1 | 0.233 | 0.816 |
| ***Dyslexic Controls*** | | | | |
| Intercept | -17.353 | 11 | -1.531 | 0.126 |
| Time (Days from ses2) | 0.008 | 0 | 1.581 | 0.114 |
| Age | 1.251 | 1 | 1.296 | 0.195 |
| Movement (Mean FD) | 1.406 | 5 | 0.287 | 0.774 |
| Num Good Runs | 1.285 | 1 | 1.484 | 0.138 |
| ***Typical Controls*** | | | | |
| Intercept | **-103.229** | **17** | **-6.179** | **6.44E-10***** |
| Time (Days from ses2) | **-0.101** | **0** | **-2.140** | **0.032*** |
| Age | **9.816** | **2** | **3.972** | **7.12E-05***** |
| Movement (Mean FD) | **118.469** | **20** | **5.935** | **2.93E-09***** |
| Num Good Runs | **12.017** | **5** | **2.225** | **0.026*** |
| **FFA2** | | | | |
| ***Intervention*** | | | | |
| Intercept | -4.912 | 12 | -0.399 | 0.690 |
| Time (Days from ses2) | -0.001 | 0 | -0.087 | 0.931 |
| Age | 0.680 | 1 | 0.541 | 0.589 |
| Movement (Mean FD) | -2.617 | 4 | -0.746 | 0.456 |
| Num Good Runs | 1.536 | 1 | 1.253 | 0.210 |
| ***Typical Controls*** | | | | |
| Intercept | **-393.827** | **14** | **-27.369** | **6.33E-165***** |
| Time (Days from ses2) | 0.007 | 0 | 0.084 | 0.933 |
| Age | **50.138** | **5** | **10.576** | **3.84E-26***** |
| Movement (Mean FD) | 10.558 | 16 | 0.674 | 0.501 |
| Num Good Runs | 1.365 | 11 | 0.122 | 0.903 |

**Table S8 | Longitudinal GLMER Results for Change in ROI Detection**

Results from a generalized linear mixed model (GLMER) calculating presence of Visual Word Form Area (VWFA) 1 & 2 and Fusiform Face Area (FFA) 2 as a function of time (in days from baseline scan) with added covariates of participant age, mean framewise displacement during the scan (mean FD), and the number of usable runs of the experiment along with a random intercept by participant. The model was fit separately for each participant group. Note that due to convergence issues changes in some regions of interest (ROIs) for were not able to be calculated. All instances where there were no model convergence issues are reported. Significant results are displayed in bold and asterisks indicate the degree of significance ($p < 0.001$: ***, $p < 0.01$: **, $p < 0.05$: *, $p < 0.1$: .). Significant results of $p<0.05$ are indicated with bold font. Note: No corrections for multiple comparisons were made due to the use of small, manually-defined ROIs. Source data are provided in a public data repository.

| Size ~ Time * Subgroup + Age + Movement + RunNums + (1 \| Participant) | | | | | | | | | | | | | | | | | | | | |
|---|---|---|---|---|---|---|---|---|---|---|---|---|---|---|---|---|---|---|---|---|
| | VWFA1 | | | | | VWFA2 | | | | | FFA1 | | | | | FFA2 | | | | |
| | β | Std. Err | DOF | t | p | β | Std. Err | DOF | t | p | β | Std. Err | DOF | t | p | β | Std. Err | DOF | t | p |
| Intercept | -160.639 | 121.414 | 144 | -1.323 | 0.188 | -51.921 | 146.375 | 102 | -0.355 | 0.724 | 187.130 | 224.958 | 113 | 0.832 | 0.407 | 19.159 | 180.138 | 122 | 0.106 | 0.915 |
| Time (Days from ses2) | **0.320** | **0.059** | **248** | **5.467** | **1.12E-07 ***** | **0.245** | **0.042** | **238** | **5.881** | **1.37E-08 ***** | 0.116 | 0.084 | 237 | 1.388 | 0.166 | 0.034 | 0.068 | 244 | 0.503 | 0.615 |
| Group: Dys Ctrl | -45.181 | 38.595 | 127 | -1.171 | 0.244 | -20.868 | 48.432 | 91 | -0.431 | 0.668 | -6.904 | 73.072 | 98 | -0.094 | 0.925 | 9.146 | 58.409 | 106 | 0.157 | 0.876 |
| Group: Typ Ctrl | **103.273** | **35.274** | **124** | **2.928** | **0.004 **** | **217.092** | **44.493** | **90** | **4.879** | **4.55E-06 ***** | -10.380 | 66.989 | 96 | -0.155 | 0.877 | **-115.299** | **53.534** | **104** | **-2.154** | **0.034 *** |
| Age | 17.074 | 10.190 | 89 | 1.676 | 0.097 . | 7.708 | 13.633 | 83 | 0.565 | 0.573 | 15.074 | 20.133 | 81 | 0.749 | 0.456 | 17.718 | 16.052 | 87 | 1.104 | 0.273 |
| Movement (Mean FD) | -69.897 | 81.790 | 318 | -0.855 | 0.393 | -51.221 | 63.867 | 277 | -0.802 | 0.423 | **-363.557** | **123.694** | **298** | **-2.939** | **0.004 **** | **-223.259** | **100.886** | **302** | **-2.213** | **0.028 *** |
| Num Good Runs | 20.078 | 14.047 | 300 | 1.429 | 0.154 | 15.131 | 10.533 | 258 | 1.436 | 0.152 | **64.451** | **20.730** | **273** | **3.109** | **0.002 **** | **69.988** | **16.936** | **278** | **4.132** | **4.75E-05 ***** |
| Time * Group: Dys Ctrl | -0.083 | 0.133 | 257 | -0.619 | 0.536 | -0.165 | 0.096 | 241 | -1.723 | 0.086 . | 0.049 | 0.191 | 243 | 0.257 | 0.797 | 0.257 | 0.157 | 249 | 1.638 | 0.103 |
| Time * Group: Typ Ctrl | 0.043 | 0.130 | 263 | 0.333 | 0.740 | **-0.233** | **0.094** | **242** | **-2.475** | **0.014 *** | 0.016 | 0.188 | 246 | 0.085 | 0.932 | -0.158 | 0.154 | 252 | -1.029 | 0.305 |

**Table S9 | Longitudinal LME Results for ROI Size**

Results from a linear mixed effects model (LME) calculating size of Visual Word Form Area (VWFA) 1 & 2 and Fusiform Face Area (FFA) 1 & 2 as a function of the interaction between time (in days from baseline scan) and participant group with added covariates of participant age, mean framewise displacement during the scan (mean FD), and the number of usable runs of the experiment along with a random intercept by participant. The intervention group is treated as the reference category. Significant results are displayed in bold and asterisks indicate the degree of significance ($p < 0.001$: ***, $p < 0.01$: **, $p < 0.05$: *, $p < 0.1$: .). Significant results of $p<0.05$ are indicated with bold font. Note: No corrections for multiple comparisons were made due to the use of small, manually-defined ROIs. Source data are provided in a public data repository.

| ROI Log Size Differences (Intervention > Typical Control) | | | | | | | | |
|---|---|---|---|---|---|---|---|---|
| | **ses-2 (n = 42 int \| 15 ctrl)** | | **ses-3 (n = 29 int \| 22 ctrl)** | | **ses-4 (n = 42 int \| 24 ctrl)** | | **ses-5 (n = 40 int \| 12 ctrl)** | |
| **ROI** | **t** | **p** | **t** | **p** | **t** | **p** | **t** | **p** |
| **VWFA1** | **-4.387** | **4.40E-05 ***** | **-3.856** | **3.68E-04 ***** | **-2.551** | **0.016 *** | **-3.037** | **0.004 **** |
| **VWFA2** | **-4.600** | **2.30E-05 ***** | **-4.188** | **1.17E-04 ***** | **-2.416** | **0.022 *** | 0.106 | 0.917 |
| **FFA1** | 0.541 | 0.592 | -1.492 | 0.142 | 0.046 | 0.964 | 0.979 | 0.347 |
| **FFA2** | **2.061** | **0.049 *** | -0.362 | 0.719 | 1.992 | 0.064 | 1.621 | 0.129 |

**Table S10 | Session Specific Differences in ROI Size**

Results from a series of two-sided t-tests comparing the difference in ROI size between intervention participants and typical control participants. Significant results are displayed in bold and asterisks indicate the degree of significance ($p < 0.001$: ***, $p < 0.01$: **, $p < 0.05$: *, $p < 0.1$: .) Significant results of $p<0.05$ are indicated with bold font. Source data are provided in a public data repository.

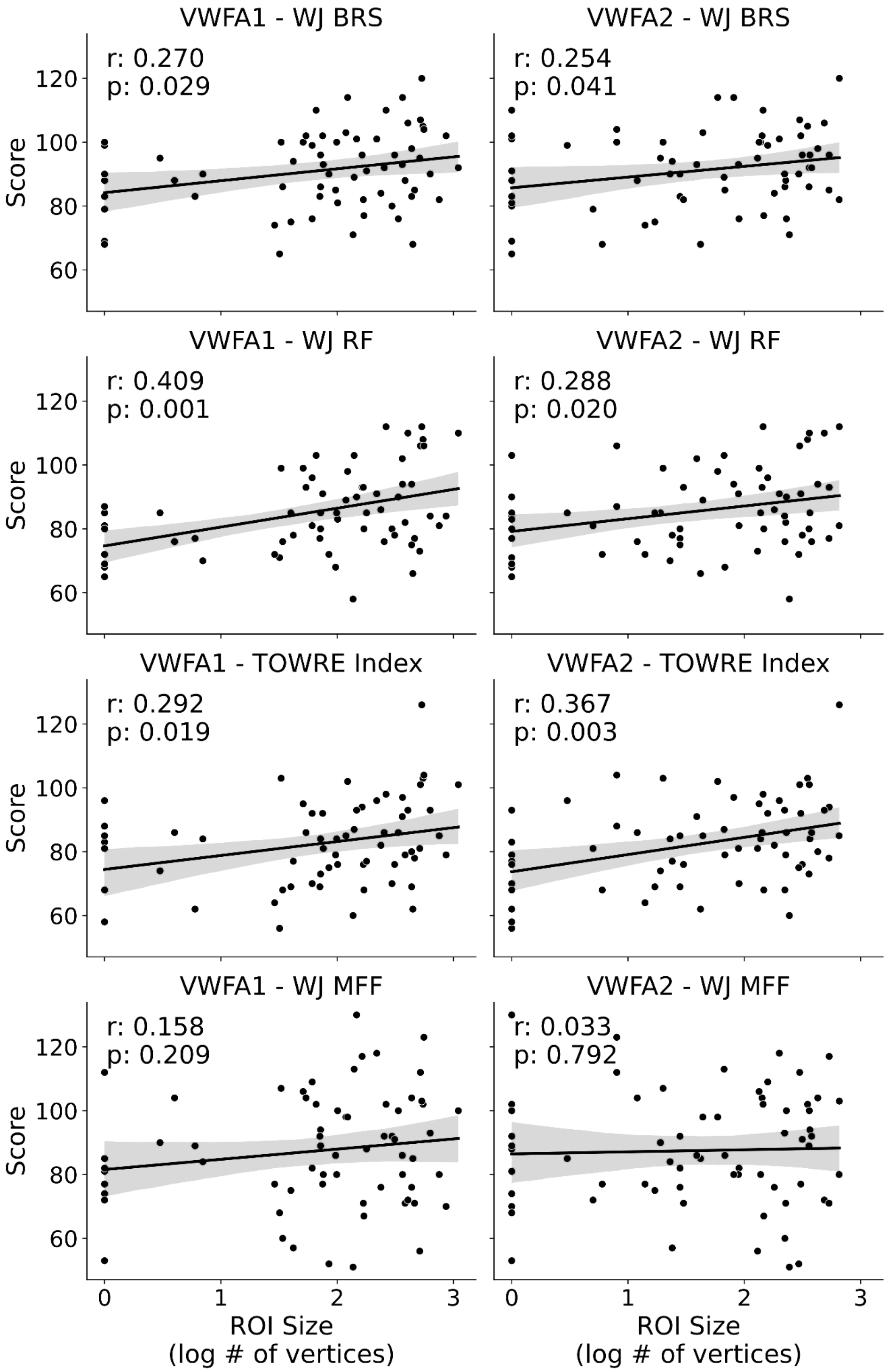


**Figure S2 | Correlations Between VWFA Size and Assessment Score at the One-Year Follow-Up**
The size of Visual Word Form Area (VWFA) defined as log10 number of vertices correlated with reading and math abilities. Woodcock-Johnson Basic Reading Skill score (WJ BRS), Reading Fluency score (WJ RF), and Test of Word Reading Efficiency Index (TOWRE). Source data are provided in a public data repository.

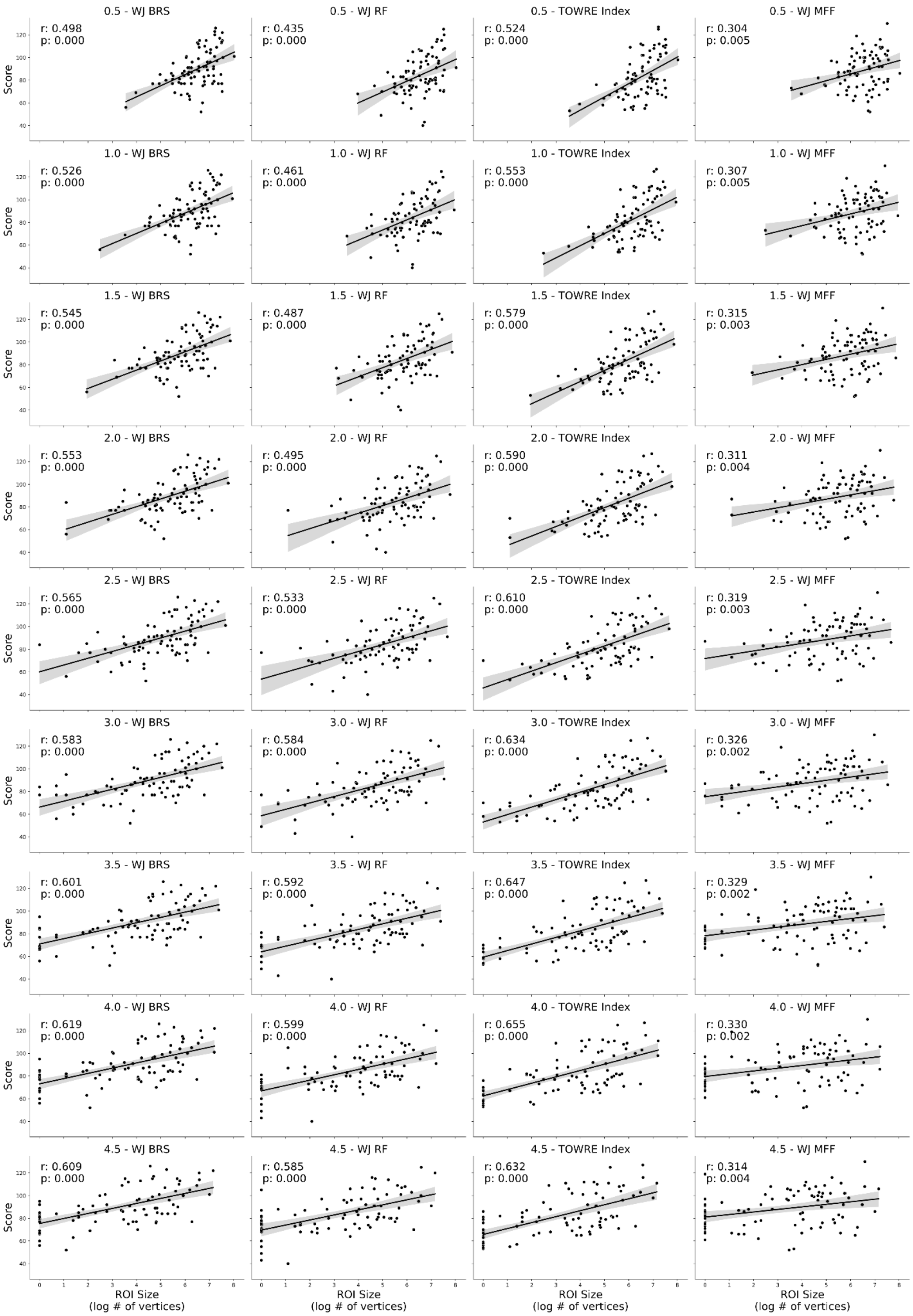


**Figure S3 | Correlations Between VWFA Size and Assessment Score Across Thresholds**
The size of Visual Word Form Area (VWFA) defined using various t statistic thresholds (ranging from 0.5 to 4.5 in increments of 0.5) from a text > other visual stimuli contrast correlated with reading and math abilities. Woodcock-Johnson Basic Reading Skill score (WJ BRS), Reading Fluency score (WJ RF), and Test of Word Reading Efficiency Index (TOWRE). Source data are provided in a public data repository.

| | PSC ~ Time * Category * Subgroup + Age + Movement + RunNums + (1 \| Participant) | | | | | | | | | | | | | | | | | | | |
|---|---|---|---|---|---|---|---|---|---|---|---|---|---|---|---|---|---|---|---|---|
| | VWFA1 | | | | | VWFA2 | | | | | FFA1 | | | | | FFA2 | | | | |
| | β | Std. Err | DOF | t | p | β | Std. Err | DOF | t | p | β | Std. Err | DOF | t | p | β | Std. Err | DOF | t | p |
| Intercept (Intervention \| Text) | 0.202 | 0.546 | 75 | 0.370 | 0.713 | 0.041 | 0.345 | 70 | 0.119 | 0.906 | 0.285 | 0.432 | 96 | 0.659 | 0.511 | 0.413 | 0.226 | 102 | 1.827 | 0.071. |
| Time (days from ses2) | 0.000 | 0.000 | 482 | 1.414 | 0.158 | **0.0003** | **0.000** | **425** | **3.563** | **4.08E-04*** ** | 0.000 | 0.000 | 558 | -0.224 | 0.823 | 0.000 | 0.000 | 557 | -0.042 | 0.967 |
| Category: Not Text | **-0.390** | **0.034** | **481** | **-11.435** | **5.98E-27*** ** | **-0.312** | **0.026** | **423** | **-12.159** | **2.21E-29*** ** | **0.823** | **0.035** | **557** | **23.578** | **9.51E-86*** ** | **0.409** | **0.022** | **553** | **18.616** | **2.00E-60*** ** |
| Group: Dys Ctrl | 0.092 | 0.207 | 79 | 0.442 | 0.660 | -0.019 | 0.119 | 74 | -0.163 | 0.871 | -0.111 | 0.151 | 100 | -0.740 | 0.461 | -0.023 | 0.080 | 111 | -0.288 | 0.774 |
| Group: Typ Ctrl | **0.372** | **0.162** | **77** | **2.302** | **0.024*** | 0.177 | 0.097 | 73 | 1.821 | 0.073. | 0.203 | 0.136 | 100 | 1.491 | 0.139 | 0.056 | 0.073 | 109 | 0.772 | 0.441 |
| Age | 0.101 | 0.052 | 68 | 1.951 | 0.055. | 0.059 | 0.033 | 61 | 1.814 | 0.075. | 0.062 | 0.041 | 83 | 1.508 | 0.135 | 0.007 | 0.021 | 83 | 0.311 | 0.756 |
| Movement (Mean FD) | -0.144 | 0.147 | 512 | -0.974 | 0.330 | -0.131 | 0.113 | 452 | -1.157 | 0.248 | -0.276 | 0.151 | 598 | -1.825 | 0.069. | **-0.307** | **0.093** | **620** | **-3.296** | **0.001**** |
| # of Good Runs | 0.027 | 0.025 | 490 | 1.104 | 0.270 | **-0.047** | **0.017** | **435** | **-2.668** | **0.008**** | -0.011 | 0.024 | 587 | -0.438 | 0.661 | -0.026 | 0.015 | 588 | -1.694 | 0.091. |
| Time * Category: Not Text | **-0.0004** | **0.0002** | **481** | **-1.972** | **0.049*** | 0.000 | 0.000 | 423 | -1.874 | 0.062. | 0.000 | 0.000 | 557 | -0.115 | 0.908 | 0.000 | 0.000 | 553 | -0.168 | 0.867 |
| Time * Group: Dys Ctrl | 0.000 | 0.000 | 483 | 1.183 | 0.237 | 0.000 | 0.000 | 426 | -0.149 | 0.881 | 0.000 | 0.000 | 561 | -0.018 | 0.986 | 0.000 | 0.000 | 558 | 0.737 | 0.462 |
| Time * Group: Typ Ctrl | **-0.001** | **0.0003** | **484** | **-2.677** | **0.008**** | 0.000 | 0.000 | 427 | -1.081 | 0.280 | 0.000 | 0.000 | 563 | -1.039 | 0.299 | 0.000 | 0.000 | 562 | 0.861 | 0.389 |
| Category: Not Text * Group: Dys Ctrl | -0.003 | 0.088 | 481 | -0.034 | 0.973 | -0.031 | 0.060 | 423 | -0.510 | 0.611 | -0.110 | 0.078 | 557 | -1.406 | 0.160 | 0.000 | 0.050 | 553 | 0.004 | 0.997 |
| Category: Not Text * Group: Typ Ctrl | **-0.179** | **0.066** | **481** | **-2.709** | **0.007**** | **-0.114** | **0.048** | **423** | **-2.359** | **0.019*** | **-0.201** | **0.070** | **557** | **-2.859** | **0.004**** | **-0.157** | **0.045** | **553** | **-3.526** | **4.57E-04*** ** |
| Time * Category: Not Text * Group: Dys Ctrl | 0.000 | 0.000 | 481 | -0.300 | 0.764 | 0.000 | 0.000 | 423 | 0.819 | 0.413 | 0.000 | 0.000 | 557 | 0.870 | 0.385 | 0.000 | 0.000 | 553 | 0.501 | 0.617 |
| Time * Category: Not Text * Group: Typ Ctrl | 0.000 | 0.000 | 481 | 0.936 | 0.350 | 0.000 | 0.000 | 423 | 1.020 | 0.309 | 0.000 | 0.000 | 557 | -0.494 | 0.621 | 0.000 | 0.000 | 553 | -0.829 | 0.408 |

**Table S11 | Longitudinal LME Results for Percent Signal Change Over Time**

Results from a linear mixed effects model (LME) calculating mean response (in units of percent signal change, PSC) of Visual Word Form Area (VWFA) 1 & 2 and Fusiform Face Area (FFA) 1 & 2 as a function of the interaction between time (in days from baseline scan) and participant group with added covariates of participant age, mean framewise displacement during the scan (mean FD), and the number of usable runs of the experiment along with a random intercept by participant. The intervention group and response to the text are treated as the reference categories. Significant results are displayed in bold and asterisks indicate the degree of significance ($p < 0.001$: ***, $p < 0.01$: **, $p < 0.05$: *, $p < 0.1$: .). Significant results of $p<0.05$ are indicated with bold font. Note: No corrections for multiple comparisons were made due to the use of small, manually-defined ROIs. Source data are provided in a public data repository.

| | **Text Selectivity ~ Time * Subgroup + Age + Movement + RunNums + (1 \| Participant)** | | | | | | | | | | | | | | | | | | | |
|---|---|---|---|---|---|---|---|---|---|---|---|---|---|---|---|---|---|---|---|---|
| | **VWFA1** | | | | | **VWFA2** | | | | | **FFA1** | | | | | **FFA2** | | | | |
| | **β** | **Std. Err** | **DOF** | **t** | **p** | **β** | **Std. Err** | **DOF** | **t** | **p** | **β** | **Std. Err** | **DOF** | **t** | **p** | **β** | **Std. Err** | **DOF** | **t** | **p** |
| Intercept (Intervention) | 0.034 | 0.060 | 129 | 0.562 | 0.575 | 0.086 | 0.060 | 119 | 1.417 | 0.159 | **-0.142** | **0.058** | **122** | **-2.442** | **0.016*** | -0.054 | 0.062 | 86 | -0.872 | 0.386 |
| Time (days from ses2) | **0.000** | **0.000** | **215** | **2.511** | **0.013*** | **0.000** | **0.000** | **189** | **2.027** | **0.044*** | 0.000 | 0.000 | 241 | 0.210 | 0.834 | 0.000 | 0.000 | 226 | -0.503 | 0.616 |
| Group: Dys Ctrl | -0.014 | 0.021 | 115 | -0.682 | 0.496 | 0.029 | 0.019 | 109 | 1.546 | 0.125 | 0.013 | 0.019 | 102 | 0.670 | 0.505 | -0.014 | 0.021 | 78 | -0.691 | 0.492 |
| Group: Typ Ctrl | 0.019 | 0.016 | 107 | 1.199 | 0.233 | 0.026 | 0.015 | 104 | 1.719 | 0.089. | **0.053** | **0.017** | **102** | **3.120** | **0.002**** | **0.042** | **0.019** | **77** | **2.210** | **0.030*** |
| Age | -0.003 | 0.005 | 74 | -0.522 | 0.603 | 0.000 | 0.005 | 66 | 0.051 | 0.959 | 0.000 | 0.005 | 84 | 0.047 | 0.963 | -0.004 | 0.006 | 72 | -0.698 | 0.487 |
| Movement (Mean FD) | 0.076 | 0.040 | 272 | 1.933 | 0.054. | 0.008 | 0.044 | 241 | 0.178 | 0.859 | **-0.092** | **0.034** | **293** | **-2.737** | **0.007**** | **-0.076** | **0.024** | **259** | **-3.140** | **0.002**** |
| # of Good Runs | **0.022** | **0.007** | **247** | **3.095** | **0.002**** | 0.010 | 0.007 | 224 | 1.381 | 0.169 | -0.009 | 0.006 | 278 | -1.570 | 0.118 | -0.005 | 0.004 | 239 | -1.222 | 0.223 |
| Time * Group: Dys Ctrl | 0.000 | 0.000 | 220 | 1.095 | 0.275 | 0.000 | 0.000 | 195 | -1.637 | 0.103 | **0.000** | **0.000** | **247** | **-1.978** | **0.049*** | 0.000 | 0.000 | 227 | -0.630 | 0.529 |
| Time * Group: Typ Ctrl | 0.000 | 0.000 | 228 | -0.810 | 0.419 | 0.000 | 0.000 | 204 | -1.195 | 0.233 | 0.000 | 0.000 | 251 | -0.132 | 0.895 | 0.000 | 0.000 | 230 | 0.023 | 0.981 |

**Table S12 | Longitudinal LME Results for Change in Text Selectivity**

Results from a linear mixed effects model (LME) calculating text selectivity of Visual Word Form Area (VWFA) 1 & 2 and Fusiform Face Area (FFA) 1 & 2 as a function of the interaction between time (in days from baseline scan) and participant group with added covariates of participant age, mean framewise displacement during the scan (mean FD), and the number of usable runs of the experiment along with a random intercept by participant. The intervention group is treated as the reference category. Selectivity index is calculated as the difference between activation to text versus non text, divided by the sum of activation to all stimuli. Significant results are displayed in bold and asterisks indicate the degree of significance ($p < 0.001$: ***, $p < 0.01$: **, $p < 0.05$: *, $p < 0.1$: .). Significant results of $p<0.05$ are indicated with bold font. Note: No corrections for multiple comparisons were made due to the use of small, manually-defined ROIs. Source data are provided in a public data repository.

| Breakdown of Trait and State Scores by Group | | | | | | |
|---|---|---|---|---|---|---|
| | | | Count of Participants | | | |
| | Average Trait Score ± SD | Variability of State Score (SD) | *High Trait High State Variability* | *High Trait Low State Variability* | *Low Trait High State Variability* | *Low Trait Low State Variability* |
| **WJ BRS** | | | | | | |
| Intervention | 84.911 ± 9.1833 | 6.0469 | 8 | 14 | 14 | 8 |
| Dyslexic Control | 79.2851 ± 10.0264 | 4.6563 | 5 | 3 | 4 | 7 |
| Typical Control | 109.4965 ± 8.5242 | 3.8043 | 6 | 6 | 6 | 6 |
| **WJ RF** | | | | | | |
| Intervention | 77.567 ± 12.5996 | 5.150 | 9 | 15 | 13 | 7 |
| Dyslexic Control | 77.1711 ± 11.6583 | 3.2707 | 3 | 7 | 6 | 3 |
| Typical Control | 107.3514 ± 10.0258 | 4.4085 | 5 | 7 | 6 | 5 |
| **TOWRE** | | | | | | |
| Intervention | 75.8716 ± 9.8512 | 5.2355 | 10 | 13 | 12 | 9 |
| Dyslexic Control | 70.2105 ± 8.4551 | 3.5628 | 4 | 6 | 5 | 4 |
| Typical Control | 103.0625 ± 10.4646 | 4.2008 | 6 | 5 | 6 | 7 |
| **WJ MFF** | | | | | | |
| Intervention | 83.3886 ± 15.4096 | 4.861 | 14 | 8 | 8 | 14 |
| Dyslexic Control | 77.3509 ± 9.4687 | 4.7147 | 4 | 5 | 5 | 5 |
| Typical Control | 101.0174 ± 13.0201 | 4.1373 | 5 | 8 | 7 | 4 |

**Table S13 | Assessment State & Trait Scores Summary Table**

Mean trait score and standard deviation for each group and assessment along with the standard deviation for state scores (mean is 0 for all state scores). Participant counts were determined by calculating the median trait score for each combination of assessment and group. If a participant's trait fell above the median, they were classified as “High Trait” and everyone else was classified as “Low Trait”. The median of the absolute value of all state score standard deviations for each participant and assessment was used as the cutoff for state variability and individuals with an absolute value of standard deviation higher than the cutoff point were classified as “High State” while everyone else was “Low State”. Counts represent the number of participants in each group that had each combination of high and low trait score and state variability. Woodcock-Johnson Basic Reading Skill score (WJ BRS), Reading Fluency score (WJ RF), and Test of Word Reading Efficiency Index (TOWRE). Source data are provided in a public data repository.

| Log Size ~ Reading Trait Raw + Reading State Raw*Subgroup + Age + Movement + RunNums + (1 \| Participant) | | | | | | | | | | | | | | | | | | | | |
|---|---|---|---|---|---|---|---|---|---|---|---|---|---|---|---|---|---|---|---|---|
| | VWFA1 | | | | | VWFA2 | | | | | FFA1 | | | | | FFA2 | | | | |
| | β | Std. Err | DOF | t | p | β | Std. Err | DOF | t | p | β | Std. Err | DOF | t | p | β | Std. Err | DOF | t | p |
| WJ LWID | | | | | | | | | | | | | | | | | | | | |
| Intercept (Intervention) | -0.696 | 0.573 | 137 | -1.214 | 0.227 | -0.416 | 0.666 | 125 | -0.625 | 0.533 | 2.288 | 0.341 | 106 | 6.711 | 9.80E-10*** | 1.616 | 0.395 | 97 | 4.086 | 9.01E-05*** |
| WJ LWID Trait | 0.039 | 0.011 | 89 | 3.656 | 4.34E-04*** | 0.062 | 0.013 | 90 | 4.837 | 5.37E-06*** | 0.014 | 0.007 | 75 | 2.210 | 0.030* | -0.002 | 0.008 | 78 | -0.271 | 0.787 |
| WJ LWID State | 0.038 | 0.008 | 239 | 4.720 | 4.02E-06*** | 0.038 | 0.008 | 240 | 4.798 | 2.82E-06*** | -0.001 | 0.004 | 224 | -0.297 | 0.767 | -0.001 | 0.004 | 228 | -0.191 | 0.849 |
| Group: Dys Ctrl | -0.281 | 0.167 | 82 | -1.680 | 0.097. | -0.085 | 0.204 | 85 | -0.418 | 0.677 | -0.021 | 0.105 | 70 | -0.204 | 0.839 | -0.029 | 0.127 | 74 | -0.227 | 0.821 |
| Group: Typ Ctrl | 0.123 | 0.213 | 85 | 0.577 | 0.566 | -0.078 | 0.259 | 87 | -0.301 | 0.764 | -0.302 | 0.133 | 72 | -2.277 | 0.026* | -0.313 | 0.160 | 76 | -1.950 | 0.055. |
| Age | 0.021 | 0.067 | 83 | 0.315 | 0.753 | -0.143 | 0.081 | 85 | -1.764 | 0.081. | -0.035 | 0.042 | 71 | -0.848 | 0.399 | 0.082 | 0.050 | 75 | 1.617 | 0.110 |
| Movement (Mean FD) | -0.578 | 0.378 | 314 | -1.529 | 0.127 | -0.439 | 0.383 | 302 | -1.146 | 0.253 | -1.213 | 0.194 | 298 | -6.241 | 1.50E-09*** | -0.752 | 0.183 | 276 | -4.102 | 5.40E-05*** |
| Num Good Runs | 0.054 | 0.067 | 298 | 0.805 | 0.421 | 0.082 | 0.067 | 282 | 1.232 | 0.219 | 0.065 | 0.034 | 273 | 1.908 | 0.057. | 0.093 | 0.031 | 256 | 2.960 | 0.003** |
| Dys Ctrl * WJ LWID State | 0.012 | 0.022 | 236 | 0.540 | 0.589 | -0.007 | 0.022 | 237 | -0.336 | 0.737 | -0.007 | 0.011 | 221 | -0.614 | 0.540 | 0.004 | 0.010 | 226 | 0.385 | 0.701 |
| Typ Ctrl * WJ LWID State | -0.056 | 0.035 | 235 | -1.635 | 0.103 | -0.098 | 0.033 | 237 | -2.940 | 0.004** | -0.009 | 0.017 | 221 | -0.526 | 0.599 | -0.009 | 0.015 | 226 | -0.573 | 0.567 |
| WJ WA | | | | | | | | | | | | | | | | | | | | |
| Intercept (Intervention) | -0.881 | 0.603 | 135 | -1.461 | 0.146 | -0.574 | 0.710 | 119 | -0.809 | 0.420 | 2.267 | 0.346 | 99 | 6.544 | 2.64E-09*** | 1.566 | 0.395 | 96 | 3.968 | 1.40E-04*** |
| WJ WA Trait | 0.048 | 0.024 | 86 | 2.016 | 0.047* | 0.089 | 0.030 | 86 | 3.006 | 0.003** | 0.019 | 0.014 | 69 | 1.337 | 0.186 | -0.011 | 0.017 | 75 | -0.674 | 0.503 |
| WJ WA State | 0.043 | 0.015 | 243 | 2.914 | 0.004** | 0.068 | 0.014 | 242 | 4.881 | 1.91E-06*** | -0.006 | 0.007 | 223 | -0.898 | 0.370 | -0.008 | 0.006 | 230 | -1.184 | 0.237 |
| Group: Dys Ctrl | -0.190 | 0.192 | 84 | -0.989 | 0.326 | 0.089 | 0.238 | 85 | 0.374 | 0.709 | 0.008 | 0.116 | 68 | 0.067 | 0.947 | -0.069 | 0.137 | 74 | -0.502 | 0.617 |
| Group: Typ Ctrl | 0.358 | 0.221 | 83 | 1.616 | 0.110 | 0.228 | 0.275 | 84 | 0.829 | 0.410 | -0.221 | 0.134 | 67 | -1.650 | 0.104 | -0.271 | 0.158 | 74 | -1.715 | 0.091. |
| Age | 0.124 | 0.060 | 83 | 2.083 | 0.040* | 0.006 | 0.074 | 84 | 0.081 | 0.936 | 0.003 | 0.036 | 67 | 0.072 | 0.943 | 0.090 | 0.043 | 74 | 2.107 | 0.039* |
| Movement (Mean FD) | -0.683 | 0.389 | 313 | -1.756 | 0.080. | -0.550 | 0.385 | 297 | -1.428 | 0.154 | -1.244 | 0.192 | 293 | -6.484 | 3.78E-10*** | -0.718 | 0.182 | 277 | -3.945 | 1.01E-04*** |
| Num Good Runs | 0.092 | 0.067 | 294 | 1.379 | 0.169 | 0.090 | 0.065 | 276 | 1.384 | 0.168 | 0.062 | 0.032 | 266 | 1.917 | 0.056. | 0.110 | 0.030 | 256 | 3.615 | 3.62E-04*** |
| Dys Ctrl * WJ WA State | -0.047 | 0.035 | 240 | -1.341 | 0.181 | -0.032 | 0.033 | 239 | -0.946 | 0.345 | 0.027 | 0.017 | 221 | 1.598 | 0.112 | 0.021 | 0.015 | 228 | 1.335 | 0.183 |
| Typ Ctrl * WJ WA State | -0.039 | 0.041 | 245 | -0.949 | 0.344 | -0.075 | 0.039 | 243 | -1.898 | 0.059. | -0.005 | 0.020 | 225 | -0.246 | 0.806 | 0.011 | 0.018 | 231 | 0.587 | 0.558 |
| WJ SRF | | | | | | | | | | | | | | | | | | | | |
| Intercept (Intervention) | -0.091 | 0.676 | 116 | -0.135 | 0.893 | 1.493 | 0.778 | 108 | 1.920 | 0.058. | 2.558 | 0.392 | 91 | 6.517 | 3.96E-09*** | 1.293 | 0.453 | 89 | 2.858 | 0.005** |
| WJ SRF Trait | 0.010 | 0.006 | 87 | 1.695 | 0.094. | 0.030 | 0.007 | 88 | 4.195 | 6.48E-05*** | 0.004 | 0.004 | 72 | 1.229 | 0.223 | -0.003 | 0.004 | 77 | -0.828 | 0.410 |
| WJ SRF State | 0.033 | 0.006 | 245 | 5.548 | 7.49E-08*** | 0.040 | 0.006 | 244 | 7.222 | 6.48E-12*** | -0.002 | 0.003 | 228 | -0.641 | 0.522 | -0.001 | 0.003 | 232 | -0.542 | 0.589 |
| Group: Dys Ctrl | -0.362 | 0.175 | 83 | -2.066 | 0.042* | -0.218 | 0.209 | 84 | -1.046 | 0.298 | -0.055 | 0.105 | 68 | -0.529 | 0.599 | -0.028 | 0.124 | 74 | -0.227 | 0.821 |
| Group: Typ Ctrl | 0.421 | 0.223 | 83 | 1.885 | 0.063. | 0.047 | 0.266 | 85 | 0.178 | 0.860 | -0.214 | 0.133 | 69 | -1.608 | 0.112 | -0.257 | 0.158 | 74 | -1.630 | 0.107 |
| Age | 0.112 | 0.071 | 81 | 1.577 | 0.119 | -0.115 | 0.085 | 84 | -1.361 | 0.177 | -0.009 | 0.042 | 68 | -0.213 | 0.832 | 0.103 | 0.050 | 73 | 2.049 | 0.044* |
| Movement (Mean FD) | -0.569 | 0.373 | 312 | -1.525 | 0.128 | -0.436 | 0.366 | 298 | -1.191 | 0.235 | -1.232 | 0.190 | 296 | -6.466 | 4.13E-10*** | -0.691 | 0.182 | 279 | -3.794 | 1.81E-04*** |
| Num Good Runs | 0.049 | 0.065 | 289 | 0.765 | 0.445 | 0.034 | 0.062 | 274 | 0.540 | 0.589 | 0.071 | 0.032 | 266 | 2.194 | 0.029* | 0.120 | 0.031 | 255 | 3.913 | 1.17E-04*** |
| Dys Ctrl * WJ SRF State | -0.002 | 0.016 | 238 | -0.096 | 0.923 | -0.025 | 0.015 | 239 | -1.608 | 0.109 | 0.004 | 0.008 | 221 | 0.455 | 0.649 | 0.006 | 0.007 | 227 | 0.811 | 0.418 |
| Typ Ctrl * WJ SRF State | -0.026 | 0.011 | 238 | -2.280 | 0.023* | -0.050 | 0.011 | 239 | -4.683 | 4.74E-06*** | -0.010 | 0.006 | 222 | -1.878 | 0.062. | -0.004 | 0.005 | 228 | -0.849 | 0.397 |
| TOWRE SWE | | | | | | | | | | | | | | | | | | | | |
| Intercept (Intervention) | -0.145 | 0.596 | 127 | -0.244 | 0.808 | 0.834 | 0.671 | 115 | 1.244 | 0.216 | 2.530 | 0.351 | 101 | 7.208 | 1.07E-10*** | 1.453 | 0.407 | 94 | 3.572 | 5.63E-04*** |
| TOWRE SWE Trait | 0.017 | 0.006 | 86 | 3.009 | 0.003** | 0.035 | 0.007 | 88 | 5.193 | 1.32E-06*** | 0.008 | 0.003 | 74 | 2.269 | 0.026* | -0.001 | 0.004 | 78 | -0.209 | 0.835 |
| TOWRE SWE State | 0.028 | 0.005 | 243 | 5.539 | 7.90E-08*** | 0.033 | 0.005 | 242 | 6.921 | 4.01E-11*** | 0.000 | 0.003 | 228 | -0.025 | 0.980 | 0.002 | 0.002 | 230 | 1.031 | 0.304 |
| Group: Dys Ctrl | -0.284 | 0.172 | 82 | -1.656 | 0.102 | -0.064 | 0.202 | 83 | -0.318 | 0.751 | -0.026 | 0.105 | 70 | -0.252 | 0.802 | -0.035 | 0.126 | 74 | -0.277 | 0.783 |
| Group: Typ Ctrl | 0.289 | 0.207 | 84 | 1.400 | 0.165 | 0.039 | 0.242 | 86 | 0.163 | 0.871 | -0.281 | 0.126 | 72 | -2.226 | 0.029* | -0.328 | 0.151 | 76 | -2.171 | 0.033* |
| Age | 0.066 | 0.065 | 82 | 1.018 | 0.312 | -0.134 | 0.077 | 84 | -1.752 | 0.083. | -0.030 | 0.040 | 71 | -0.758 | 0.451 | 0.084 | 0.048 | 75 | 1.759 | 0.083. |
| Movement (Mean FD) | -0.544 | 0.373 | 310 | -1.461 | 0.145 | -0.387 | 0.358 | 295 | -1.080 | 0.281 | -1.193 | 0.193 | 294 | -6.193 | 1.98E-09*** | -0.647 | 0.182 | 274 | -3.559 | 4.39E-04*** |
| Num Good Runs | 0.035 | 0.064 | 287 | 0.540 | 0.590 | 0.030 | 0.061 | 271 | 0.486 | 0.627 | 0.063 | 0.033 | 265 | 1.911 | 0.057. | 0.109 | 0.030 | 251 | 3.569 | 4.30E-04*** |
| Dys Ctrl * TOWRE SWE State | -0.004 | 0.013 | 237 | -0.297 | 0.767 | -0.008 | 0.012 | 237 | -0.669 | 0.504 | -0.003 | 0.006 | 223 | -0.445 | 0.657 | -0.002 | 0.006 | 227 | -0.275 | 0.784 |

| | | | | | | | | | | | | | | | | | | | | |
|---|---|---|---|---|---|---|---|---|---|---|---|---|---|---|---|---|---|---|---|---|
| Typ Ctrl * TOWRE SWE State | -0.013 | 0.016 | 236 | -0.837 | 0.403 | -0.026 | 0.015 | 236 | -1.804 | 0.072. | -0.004 | 0.008 | 222 | -0.503 | 0.615 | 0.001 | 0.007 | 226 | 0.182 | 0.855 |
| TOWRE PDE | | | | | | | | | | | | | | | | | | | | |
| Intercept (Intervention) | -0.251 | 0.610 | 123 | -0.412 | 0.681 | 0.655 | 0.699 | 112 | 0.936 | 0.351 | 2.616 | 0.344 | 101 | 7.597 | 1.58E-11*** | 1.516 | 0.408 | 93 | 3.718 | 3.44E-04*** |
| TOWRE PDE Trait | 0.027 | 0.011 | 94 | 2.500 | 0.014* | 0.054 | 0.013 | 92 | 4.256 | 5.00E-05*** | 0.019 | 0.006 | 81 | 3.056 | 0.003** | 0.001 | 0.008 | 81 | 0.104 | 0.918 |
| TOWRE PDE State | 0.035 | 0.009 | 241 | 4.075 | 6.24E-05*** | 0.046 | 0.008 | 239 | 5.739 | 2.88E-08*** | -0.004 | 0.004 | 228 | -0.851 | 0.396 | -0.004 | 0.004 | 228 | -1.121 | 0.264 |
| Group: Dys Ctrl | -0.145 | 0.188 | 83 | -0.768 | 0.445 | 0.205 | 0.225 | 84 | 0.912 | 0.365 | 0.070 | 0.110 | 72 | 0.637 | 0.526 | -0.025 | 0.135 | 75 | -0.186 | 0.853 |
| Group: Typ Ctrl | 0.156 | 0.266 | 91 | 0.585 | 0.560 | -0.238 | 0.315 | 90 | -0.756 | 0.452 | -0.470 | 0.154 | 79 | -3.054 | 0.003** | -0.365 | 0.187 | 79 | -1.949 | 0.055. |
| Age | 0.099 | 0.062 | 84 | 1.609 | 0.111 | -0.058 | 0.073 | 84 | -0.787 | 0.433 | -0.035 | 0.036 | 73 | -0.968 | 0.336 | 0.071 | 0.044 | 75 | 1.624 | 0.109 |
| Movement (Mean FD) | -0.566 | 0.384 | 308 | -1.472 | 0.142 | -0.424 | 0.376 | 294 | -1.128 | 0.260 | -1.207 | 0.193 | 295 | -6.251 | 1.42E-09*** | -0.694 | 0.185 | 274 | -3.760 | 2.08E-04*** |
| Num Good Runs | 0.081 | 0.066 | 287 | 1.235 | 0.218 | 0.086 | 0.063 | 271 | 1.357 | 0.176 | 0.070 | 0.033 | 267 | 2.142 | 0.033* | 0.115 | 0.031 | 251 | 3.752 | 2.18E-04*** |
| Dys Ctrl * TOWRE PDE State | -0.037 | 0.022 | 234 | -1.624 | 0.106 | -0.049 | 0.021 | 234 | -2.340 | 0.020* | 0.007 | 0.011 | 223 | 0.632 | 0.528 | 0.012 | 0.010 | 224 | 1.185 | 0.237 |
| Typ Ctrl * TOWRE PDE State | -0.028 | 0.018 | 239 | -1.557 | 0.121 | -0.068 | 0.017 | 237 | -3.984 | 9.01E-05*** | -0.004 | 0.009 | 226 | -0.458 | 0.647 | 0.000 | 0.008 | 226 | -0.050 | 0.960 |
| WJ MFF | | | | | | | | | | | | | | | | | | | | |
| Intercept (Intervention) | -0.708 | 0.638 | 124 | -1.110 | 0.269 | 0.129 | 0.774 | 111 | 0.167 | 0.868 | 2.456 | 0.362 | 93 | 6.793 | 1.04E-09*** | 1.406 | 0.418 | 92 | 3.365 | 0.001** |
| WJ MFF Trait | -0.001 | 0.004 | 79 | -0.223 | 0.824 | 0.006 | 0.005 | 81 | 1.172 | 0.245 | 0.003 | 0.002 | 65 | 1.278 | 0.206 | -0.001 | 0.003 | 73 | -0.438 | 0.663 |
| WJ MFF State | 0.015 | 0.006 | 245 | 2.728 | 0.007** | 0.015 | 0.005 | 242 | 2.703 | 0.007** | 0.004 | 0.003 | 225 | 1.536 | 0.126 | 0.004 | 0.002 | 231 | 1.630 | 0.104 |
| Group: Dys Ctrl | -0.351 | 0.181 | 83 | -1.941 | 0.056. | -0.147 | 0.229 | 83 | -0.639 | 0.524 | -0.032 | 0.107 | 67 | -0.298 | 0.767 | -0.039 | 0.127 | 74 | -0.304 | 0.762 |
| Group: Typ Ctrl | 0.697 | 0.187 | 83 | 3.721 | 3.61E-04*** | 0.669 | 0.237 | 83 | 2.815 | 0.006** | -0.163 | 0.110 | 67 | -1.478 | 0.144 | -0.311 | 0.132 | 74 | -2.365 | 0.021* |
| Age | 0.200 | 0.063 | 80 | 3.158 | 0.002** | 0.074 | 0.081 | 82 | 0.915 | 0.363 | 0.002 | 0.037 | 65 | 0.044 | 0.965 | 0.086 | 0.045 | 73 | 1.928 | 0.058. |
| Movement (Mean FD) | -0.734 | 0.390 | 313 | -1.881 | 0.061. | -0.688 | 0.397 | 297 | -1.732 | 0.084. | -1.176 | 0.190 | 294 | -6.182 | 2.12E-09*** | -0.622 | 0.181 | 277 | -3.431 | 6.93E-04*** |
| Num Good Runs | 0.079 | 0.067 | 291 | 1.181 | 0.239 | 0.077 | 0.067 | 274 | 1.145 | 0.253 | 0.066 | 0.032 | 264 | 2.067 | 0.040* | 0.116 | 0.030 | 255 | 3.848 | 1.51E-04*** |
| Dys Ctrl * WJ MFF State | -0.005 | 0.015 | 238 | -0.320 | 0.749 | -0.002 | 0.014 | 238 | -0.111 | 0.911 | 0.001 | 0.007 | 220 | 0.198 | 0.843 | -0.001 | 0.006 | 228 | -0.100 | 0.920 |
| Typ Ctrl * WJ MFF State | -0.004 | 0.010 | 239 | -0.452 | 0.652 | -0.023 | 0.009 | 238 | -2.480 | 0.014* | -0.012 | 0.005 | 220 | -2.727 | 0.007** | -0.007 | 0.004 | 228 | -1.790 | 0.075. |

**Table S14 | LME Results for ROI Size with Raw State and Trait Scores**

Results of linear mixed effects models calculating the size (log transformed number of vertices) of a region of interest (ROI) as a function of raw assessment trait and the interaction between raw assessment state and participant group with added covariates of participant age, mean framewise displacement during the scan (mean FD), and the number of usable runs of the experiment along with a random intercept by participant. Intervention group is treated as the reference category. Trait: within-participant average over time; State: within-participant change relative to trait; VWFA: Visual Word Form Area; FFA: Fusiform Face Area; WJ LWID: Woodcock-Johnson Letter Word Identification; WJ WA: Woodcock-Johnson Word Attack; WJ SRF: Woodcock-Johnson Sentence Reading Fluency; TOWRE SWE: Test of Word Reading Efficiency - Sight Word Efficiency; TOWRE PDe: Test of Word Reading Efficiency - Phonemic Decoding Efficiency; WJ MFF: Woodcock-Johnson Math Facts Fluency; Dys Ctrl: Dyslexic Control; Typ Ctrl: Typical Control; FD: Framewise Displacement Significant results are displayed in bold and asterisks indicate the degree of significance ($p < 0.001$: ***, $p < 0.01$: **, $p < 0.05$: *, $p < 0.1$: .). Significant results of $p<0.05$ are indicated with bold font. Note: No corrections for multiple comparisons were made due to the use of small, manually-defined ROIs. Source data are provided in a public data repository.

| Presence ~ Reading Trait + Reading State * Subgroup + Age + Movement + RunNums + (1 \| Participant) | VWFA1 | | | | VWFA2 | | | | FFA2 | | | |
|---|---|---|---|---|---|---|---|---|---|---|---|---|
| | **coef** | **Std. Err** | **z** | **P>\|z\|** | **coef** | **Std. Err** | **z** | **P>\|z\|** | **coef** | **Std. Err** | **z** | **P>\|z\|** |
| **WJ BRS** | | | | | | | | | | | | |
| Intercept (Intervention) | **-19.773** | **5** | **-3.815** | **1.36E-04*** ** | **-29.562** | **9** | **-3.402** | **6.70E-04*** ** | -13.222 | 23 | -0.585 | 0.558 |
| WJ BRS Trait | **0.142** | **0** | **3.441** | **5.79E-04*** ** | **0.245** | **0** | **3.279** | **0.001** ** | -0.052 | 0 | -0.462 | 0.644 |
| WJ BRS State | 0.009 | 0 | 0.101 | 0.920 | -0.015 | 0 | -0.151 | 0.880 | -0.157 | 0 | -0.452 | 0.652 |
| Group: Dys Ctrl | 0.053 | 1 | 0.067 | 0.947 | -0.311 | 1 | -0.224 | 0.823 | 0.215 | 4 | 0.052 | 0.959 |
| Group: Typ Ctrl | 1.247 | 2 | 0.682 | 0.495 | -1.518 | 2 | -0.635 | 0.525 | 0.664 | 6 | 0.116 | 0.908 |
| Age | **1.010** | **0** | **3.056** | **0.002** ** | 0.752 | 0 | 1.634 | 0.102 | 2.090 | 2 | 0.855 | 0.393 |
| Movement (Mean FD) | -1.663 | 2 | -0.794 | 0.427 | -1.150 | 3 | -0.395 | 0.693 | -1.503 | 6 | -0.249 | 0.803 |
| Num Good Runs | 0.059 | 0 | 0.149 | 0.881 | **1.066** | **1** | **2.012** | **0.044*** | 3.041 | 2 | 1.662 | 0.096. |
| Dys Ctrl * WJ BRS State | 0.047 | 0 | 0.480 | 0.631 | 0.169 | 0 | 1.490 | 0.136 | 0.091 | 0 | 0.202 | 0.840 |
| Typ Ctrl * WJ BRS State | -0.010 | 0 | -0.021 | 0.983 | -0.175 | 0 | -0.633 | 0.526 | 1.069 | 1 | 1.180 | 0.238 |
| **WJ RF** | | | | | | | | | | | | |
| Intercept (Intervention) | **-13.464** | **4** | **-3.233** | **0.001** ** | **-22.698** | **8** | **-2.717** | **0.007** ** | 16.724 | 18 | 0.921 | 0.357 |
| WJ RF Trait | **0.088** | **0** | **2.819** | **0.005** ** | **0.237** | **0** | **3.091** | **0.002** ** | -0.246 | 0 | -0.887 | 0.375 |
| WJ RF State | 0.229 | 0 | 1.656 | 0.098. | 0.121 | 0 | 0.850 | 0.396 | -0.197 | 1 | -0.197 | 0.844 |
| Group: Dys Ctrl | 0.919 | 1 | 1.200 | 0.230 | 1.191 | 2 | 0.729 | 0.466 | 0.678 | 4 | 0.152 | 0.880 |
| Group: Typ Ctrl | 22.604 | 1135 | 0.020 | 0.984 | -0.059 | 3 | -0.023 | 0.982 | 7.073 | 12 | 0.591 | 0.555 |
| Age | **0.848** | **0** | **2.598** | **0.009** ** | 0.490 | 1 | 0.943 | 0.346 | 0.328 | 1 | 0.224 | 0.823 |
| Movement (Mean FD) | -1.578 | 2 | -0.777 | 0.437 | -2.735 | 3 | -0.877 | 0.380 | -1.257 | 8 | -0.151 | 0.880 |
| Num Good Runs | -0.055 | 0 | -0.138 | 0.890 | 0.373 | 1 | 0.743 | 0.457 | 3.681 | 3 | 1.404 | 0.160 |
| Dys Ctrl * WJ RF State | -0.116 | 0 | -0.800 | 0.424 | 0.176 | 0 | 1.093 | 0.274 | 0.043 | 1 | 0.044 | 0.965 |
| Typ Ctrl * WJ RF State | -1.592 | 111 | -0.014 | 0.989 | -0.292 | 0 | -1.103 | 0.270 | 0.118 | 1 | 0.086 | 0.932 |
| **TOWRE** | | | | | | | | | | | | |
| Intercept (Intervention) | **-16.383** | **5** | **-3.604** | **3.13E-04*** ** | **-22.109** | **7** | **-3.293** | **9.92E-04*** ** | -15.115 | 27 | -0.561 | 0.575 |
| TOWRE Trait | **0.130** | **0** | **3.023** | **0.003** ** | **0.335** | **0** | **3.926** | **8.63E-05*** ** | 0.020 | 0 | 0.135 | 0.893 |
| TOWRE State | -0.150 | 0 | -1.257 | 0.209 | -0.145 | 0 | -1.023 | 0.306 | -0.238 | 0 | -0.614 | 0.539 |
| Group: Dys Ctrl | 0.080 | 1 | 0.096 | 0.924 | -1.577 | 1 | -1.126 | 0.260 | -1.007 | 6 | -0.182 | 0.855 |
| Group: Typ Ctrl | 1.498 | 2 | 0.726 | 0.468 | -5.047 | 3 | -1.897 | 0.058. | -3.208 | 7 | -0.434 | 0.664 |
| Age | **0.808** | **0** | **2.334** | **0.020*** | -0.144 | 0 | -0.324 | 0.746 | 1.975 | 3 | 0.645 | 0.519 |

| | | | | | | | | | | | | |
|---|---|---|---|---|---|---|---|---|---|---|---|---|
| Movement (Mean FD) | -0.979 | 2 | -0.478 | 0.633 | -2.527 | 3 | -0.880 | 0.379 | -2.721 | 7 | -0.376 | 0.707 |
| Num Good Runs | 0.240 | 0 | 0.620 | 0.535 | 0.592 | 0 | 1.221 | 0.222 | 3.015 | 2 | 1.397 | 0.163 |
| Dys Ctrl * TOWRE State | 0.214 | 0 | 1.657 | 0.098. | **0.332** | **0** | **2.112** | **0.035*** | -0.019 | 1 | -0.034 | 0.973 |
| Typ Ctrl * TOWRE State | 0.143 | 0 | 0.342 | 0.732 | 0.160 | 0 | 0.647 | 0.517 | 0.141 | 1 | 0.263 | 0.793 |
| **WJ MFF** | | | | | | | | | | | | |
| Intercept (Intervention) | **-14.750** | **5** | **-3.079** | **0.002**** | -11.241 | 6 | -1.922 | 0.055. | -0.467 | 16 | -0.030 | 0.976 |
| WJ MFF Trait | 0.046 | 0 | 1.655 | 0.098. | 0.050 | 0 | 1.294 | 0.196 | -0.020 | 0 | -0.186 | 0.852 |
| WJ MFF State | -0.040 | 0 | -0.534 | 0.593 | 0.124 | 0 | 1.128 | 0.259 | -0.102 | 0 | -0.288 | 0.773 |
| Group: Dys Ctrl | 0.435 | 1 | 0.506 | 0.613 | 0.704 | 1 | 0.523 | 0.601 | 0.533 | 4 | 0.139 | 0.890 |
| Group: Typ Ctrl | **4.703** | **2** | **2.684** | **0.007**** | **4.047** | **2** | **2.096** | **0.036*** | -1.922 | 5 | -0.423 | 0.672 |
| Age | **1.220** | **0** | **3.281** | **0.001**** | 0.729 | 0 | 1.726 | 0.084. | 0.519 | 1 | 0.381 | 0.703 |
| Movement (Mean FD) | -2.089 | 2 | -1.012 | 0.311 | -4.288 | 3 | -1.635 | 0.102 | 0.270 | 6 | 0.047 | 0.962 |
| Num Good Runs | 0.269 | 0 | 0.720 | 0.471 | 0.552 | 0 | 1.230 | 0.219 | 2.359 | 2 | 1.482 | 0.138 |
| Dys Ctrl * WJ MFF State | 0.018 | 0 | 0.198 | 0.843 | -0.103 | 0 | -0.831 | 0.406 | 0.287 | 0 | 0.588 | 0.556 |
| Typ Ctrl * WJ MFF State | 0.043 | 0 | 0.158 | 0.874 | -0.235 | 0 | -1.359 | 0.174 | 0.062 | 0 | 0.132 | 0.895 |

** Due to model convergence issues, we were not able to fit a generalized linear model to the FFA1 data.*

**Table S15 | Longitudinal GLMER Results for ROI Presence with State and Trait Scores**

Results of generalized linear mixed models (GLMER) calculating presence of a region of interest (ROI) as a function of assessment trait and the interaction between assessment state and participant group with added covariates of participant age, mean framewise displacement during the scan (mean FD), and the number of usable runs of the experiment along with a random intercept by participant. Intervention group is treated as the reference category. Trait: within-participant average over time; State: within-participant change relative to trait; VWFA: Visual Word Form Area; FFA: Fusiform Face Area; WJ BRS: Woodcock-Johnson Basic Reading Skills; WJ RF: Woodcock-Johnson Reading Fluency; TOWRE: Test of Word Reading Efficiency; WJ MFF: Woodcock-Johnson Math Facts Fluency; Dys Ctrl: Dyslexic Control; Typ Ctrl: Typical Control; FD: Framewise Displacement Significant results are displayed in bold and asterisks indicate the degree of significance ($p < 0.001$: ***, $p < 0.01$: **, $p < 0.05$: *, $p < 0.1$: .). Significant results of $p<0.05$ are indicated with bold font. Note: No corrections for multiple comparisons were made due to the use of small, manually-defined ROIs. Source data are provided in a public data repository.

| % Signal Change ~ Reading Trait + Reading State * Subgroup + Age + Movement + RunNums + (1 \| Participant) | | | | | | | | | | | | | | | | | | | | |
|---|---|---|---|---|---|---|---|---|---|---|---|---|---|---|---|---|---|---|---|---|
| | **VWFA1** | | | | | **VWFA2** | | | | | **FFA1** | | | | | **FFA2** | | | | |
| | **β** | **Std. Err** | **DOF** | **t** | **p** | **β** | **Std. Err** | **DOF** | **t** | **p** | **β** | **Std. Err** | **DOF** | **t** | **p** | **β** | **Std. Err** | **DOF** | **t** | **p** |
| **WJ BRS** | | | | | | | | | | | | | | | | | | | | |
| Intercept (Intervention) | 0.079 | 0.889 | 78 | 0.089 | 0.929 | -0.744 | 0.555 | 72 | -1.340 | 0.184 | 0.271 | 0.669 | 103 | 0.405 | 0.686 | **0.971** | **0.360** | **108** | **2.699** | **0.008**** |
| WJ BRS Trait | 0.000 | 0.008 | 70 | -0.053 | 0.958 | 0.007 | 0.005 | 60 | 1.491 | 0.141 | 0.005 | 0.006 | 83 | 0.793 | 0.430 | -0.004 | 0.003 | 87 | -1.176 | 0.243 |
| WJ BRS State | 0.001 | 0.003 | 484 | 0.309 | 0.757 | **0.006** | **0.002** | **428** | **2.523** | **0.012*** | -0.003 | 0.004 | 566 | -0.736 | 0.462 | -0.002 | 0.002 | 564 | -0.861 | 0.390 |
| Group: Dys Ctrl | 0.119 | 0.206 | 68 | 0.579 | 0.564 | -0.007 | 0.113 | 60 | -0.064 | 0.949 | -0.119 | 0.146 | 81 | -0.815 | 0.417 | -0.015 | 0.077 | 83 | -0.192 | 0.848 |
| Group: Typ Ctrl | 0.235 | 0.249 | 68 | 0.944 | 0.348 | -0.076 | 0.150 | 59 | -0.509 | 0.613 | -0.062 | 0.201 | 81 | -0.309 | 0.758 | 0.072 | 0.105 | 84 | 0.689 | 0.493 |
| Age | 0.100 | 0.052 | 67 | 1.929 | 0.058. | 0.056 | 0.032 | 61 | 1.723 | 0.090. | 0.059 | 0.042 | 82 | 1.421 | 0.159 | 0.008 | 0.022 | 84 | 0.359 | 0.721 |
| Movement (Mean FD) | -0.160 | 0.211 | 530 | -0.758 | 0.449 | -0.157 | 0.167 | 477 | -0.938 | 0.349 | -0.272 | 0.264 | 640 | -1.029 | 0.304 | **-0.377** | **0.139** | **630** | **-2.716** | **0.007**** |
| Num Good Runs | 0.021 | 0.037 | 504 | 0.559 | 0.577 | -0.028 | 0.028 | 455 | -1.019 | 0.309 | -0.001 | 0.045 | 636 | -0.026 | 0.979 | -0.031 | 0.024 | 627 | -1.247 | 0.213 |
| Dys Ctrl * WJ BRS State | 0.000 | 0.009 | 484 | -0.037 | 0.971 | -0.006 | 0.007 | 427 | -0.895 | 0.372 | -0.001 | 0.011 | 565 | -0.108 | 0.914 | 0.001 | 0.006 | 562 | 0.215 | 0.830 |
| Typ Ctrl * WJ BRS State | 0.004 | 0.009 | 490 | 0.395 | 0.693 | -0.006 | 0.007 | 437 | -0.898 | 0.370 | 0.011 | 0.013 | 578 | 0.872 | 0.384 | 0.011 | 0.007 | 567 | 1.630 | 0.104 |
| **WJ RF** | | | | | | | | | | | | | | | | | | | | |
| Intercept (Intervention) | 0.254 | 0.698 | 78 | 0.364 | 0.717 | -0.361 | 0.434 | 72 | -0.831 | 0.409 | 0.433 | 0.543 | 106 | 0.798 | 0.426 | **0.905** | **0.289** | **108** | **3.128** | **0.002**** |
| WJ RF Trait | -0.003 | 0.006 | 69 | -0.430 | 0.669 | 0.006 | 0.004 | 59 | 1.403 | 0.166 | 0.004 | 0.005 | 81 | 0.827 | 0.411 | -0.004 | 0.003 | 85 | -1.661 | 0.100 |
| WJ RF State | 0.005 | 0.004 | 485 | 1.372 | 0.171 | **0.007** | **0.003** | **432** | **2.425** | **0.016*** | -0.002 | 0.005 | 568 | -0.390 | 0.697 | -0.002 | 0.003 | 565 | -0.569 | 0.570 |
| Group: Dys Ctrl | 0.123 | 0.201 | 68 | 0.614 | 0.542 | -0.031 | 0.110 | 59 | -0.282 | 0.779 | -0.149 | 0.142 | 80 | -1.044 | 0.300 | 0.005 | 0.075 | 81 | 0.072 | 0.943 |
| Group: Typ Ctrl | 0.303 | 0.235 | 67 | 1.291 | 0.201 | -0.061 | 0.148 | 58 | -0.413 | 0.681 | -0.087 | 0.196 | 78 | -0.445 | 0.657 | 0.104 | 0.102 | 80 | 1.021 | 0.310 |
| Age | 0.105 | 0.053 | 67 | 1.991 | 0.051. | 0.045 | 0.033 | 60 | 1.352 | 0.181 | 0.059 | 0.043 | 80 | 1.366 | 0.176 | 0.014 | 0.022 | 80 | 0.616 | 0.539 |
| Movement (Mean FD) | -0.200 | 0.206 | 530 | -0.969 | 0.333 | -0.202 | 0.166 | 475 | -1.222 | 0.222 | -0.355 | 0.260 | 634 | -1.365 | 0.173 | **-0.387** | **0.136** | **625** | **-2.841** | **0.005**** |
| Num Good Runs | 0.011 | 0.036 | 501 | 0.303 | 0.762 | **-0.053** | **0.027** | **452** | **-1.990** | **0.047*** | -0.012 | 0.044 | 621 | -0.283 | 0.778 | -0.027 | 0.024 | 610 | -1.151 | 0.250 |
| Dys Ctrl * WJ RF State | -0.005 | 0.012 | 482 | -0.418 | 0.676 | 0.006 | 0.008 | 424 | 0.672 | 0.502 | 0.006 | 0.014 | 555 | 0.432 | 0.666 | 0.005 | 0.008 | 552 | 0.605 | 0.545 |
| Typ Ctrl * WJ RF State | -0.013 | 0.008 | 482 | -1.731 | 0.084. | -0.008 | 0.006 | 425 | -1.349 | 0.178 | -0.012 | 0.010 | 557 | -1.104 | 0.270 | 0.000 | 0.006 | 553 | -0.009 | 0.993 |
| **TOWRE** | | | | | | | | | | | | | | | | | | | | |
| Intercept (Intervention) | 0.167 | 0.772 | 77 | 0.217 | 0.829 | -0.409 | 0.496 | 70 | -0.824 | 0.413 | 0.804 | 0.566 | 108 | 1.421 | 0.158 | **1.023** | **0.297** | **113** | **3.444** | **8.06E-04***** |
| TOWRE Trait | -0.002 | 0.008 | 68 | -0.251 | 0.803 | 0.006 | 0.005 | 58 | 1.097 | 0.277 | -0.002 | 0.006 | 85 | -0.276 | 0.783 | **-0.007** | **0.003** | **88** | **-2.065** | **0.042*** |
| TOWRE State | 0.003 | 0.004 | 478 | 0.671 | 0.503 | **0.007** | **0.003** | **423** | **2.526** | **0.012*** | -0.002 | 0.005 | 564 | -0.313 | 0.754 | -0.001 | 0.003 | 564 | -0.249 | 0.803 |
| Group: Dys Ctrl | 0.116 | 0.207 | 68 | 0.561 | 0.577 | 0.016 | 0.115 | 59 | 0.136 | 0.893 | -0.148 | 0.147 | 81 | -1.002 | 0.319 | -0.028 | 0.076 | 84 | -0.369 | 0.713 |
| Group: Typ Ctrl | 0.285 | 0.274 | 68 | 1.037 | 0.304 | -0.047 | 0.169 | 58 | -0.280 | 0.780 | 0.102 | 0.214 | 84 | 0.478 | 0.634 | 0.162 | 0.110 | 87 | 1.472 | 0.145 |
| Age | 0.103 | 0.053 | 68 | 1.932 | 0.058. | 0.046 | 0.033 | 60 | 1.383 | 0.172 | 0.064 | 0.044 | 82 | 1.460 | 0.148 | 0.019 | 0.022 | 84 | 0.844 | 0.401 |
| Movement (Mean FD) | -0.162 | 0.209 | 521 | -0.774 | 0.439 | -0.157 | 0.167 | 465 | -0.939 | 0.348 | -0.338 | 0.265 | 630 | -1.276 | 0.202 | **-0.393** | **0.138** | **619** | **-2.844** | **0.005**** |
| Num Good Runs | 0.022 | 0.036 | 492 | 0.615 | 0.539 | -0.043 | 0.027 | 441 | -1.616 | 0.107 | -0.006 | 0.044 | 621 | -0.135 | 0.892 | -0.028 | 0.024 | 613 | -1.185 | 0.236 |
| Dys Ctrl * TOWRE State | 0.000 | 0.013 | 478 | 0.002 | 0.998 | **-0.019** | **0.008** | **419** | **-2.248** | **0.025*** | 0.005 | 0.014 | 557 | 0.346 | 0.729 | -0.001 | 0.007 | 557 | -0.118 | 0.906 |

| | | | | | | | | | | | | | | | | | | | | |
|---|---|---|---|---|---|---|---|---|---|---|---|---|---|---|---|---|---|---|---|---|
| Typ Ctrl * TOWRE State | 0.002 | 0.008 | 475 | 0.266 | 0.790 | -0.011 | 0.006 | 418 | -1.793 | 0.074. | 0.009 | 0.011 | 555 | 0.768 | 0.443 | 0.005 | 0.006 | 554 | 0.747 | 0.456 |
| **WJ MFF** | | | | | | | | | | | | | | | | | | | | |
| Intercept (Intervention) | 0.554 | 0.695 | 75 | 0.797 | 0.428 | -0.142 | 0.435 | 71 | -0.327 | 0.745 | **1.148** | **0.551** | **103** | **2.085** | **0.040*** | **1.188** | **0.279** | **108** | **4.260** | **4.37E-05***** |
| WJ MFF Trait | -0.006 | 0.005 | 66 | -1.228 | 0.224 | 0.002 | 0.003 | 58 | 0.507 | 0.614 | -0.006 | 0.004 | 77 | -1.477 | 0.144 | **-0.006** | **0.002** | **80** | **-3.305** | **0.001**** |
| WJ MFF State | -0.001 | 0.004 | 488 | -0.268 | 0.789 | 0.001 | 0.003 | 434 | 0.393 | 0.694 | 0.000 | 0.005 | 573 | -0.066 | 0.948 | -0.002 | 0.003 | 573 | -0.554 | 0.580 |
| Group: Dys Ctrl | 0.101 | 0.201 | 68 | 0.503 | 0.617 | -0.021 | 0.112 | 60 | -0.186 | 0.853 | -0.175 | 0.143 | 80 | -1.225 | 0.224 | -0.036 | 0.072 | 83 | -0.495 | 0.622 |
| Group: Typ Ctrl | 0.330 | 0.178 | 67 | 1.851 | 0.069. | 0.077 | 0.106 | 59 | 0.723 | 0.473 | 0.159 | 0.146 | 80 | 1.090 | 0.279 | 0.091 | 0.073 | 83 | 1.242 | 0.218 |
| Age | 0.099 | 0.052 | 67 | 1.925 | 0.059. | 0.053 | 0.033 | 61 | 1.623 | 0.110 | 0.066 | 0.041 | 82 | 1.612 | 0.111 | 0.007 | 0.020 | 85 | 0.324 | 0.747 |
| Movement (Mean FD) | -0.164 | 0.206 | 535 | -0.796 | 0.426 | -0.237 | 0.166 | 479 | -1.428 | 0.154 | -0.296 | 0.259 | 642 | -1.142 | 0.254 | **-0.369** | **0.134** | **627** | **-2.748** | **0.006**** |
| Num Good Runs | 0.022 | 0.036 | 505 | 0.607 | 0.544 | -0.042 | 0.027 | 455 | -1.576 | 0.116 | -0.009 | 0.043 | 633 | -0.216 | 0.829 | -0.029 | 0.023 | 627 | -1.260 | 0.208 |
| Dys Ctrl * WJ MFF State | 0.007 | 0.012 | 497 | 0.555 | 0.579 | 0.007 | 0.008 | 429 | 0.904 | 0.367 | 0.006 | 0.011 | 564 | 0.576 | 0.565 | -0.002 | 0.006 | 573 | -0.366 | 0.715 |
| Typ Ctrl * WJ MFF State | -0.009 | 0.008 | 486 | -1.082 | 0.280 | 0.002 | 0.007 | 429 | 0.333 | 0.739 | -0.017 | 0.011 | 564 | -1.510 | 0.132 | 0.000 | 0.006 | 564 | -0.072 | 0.943 |

**Table S16 | Longitudinal LME Results for Percent Signal Change with State and Trait Scores**

Results of linear mixed effects models (LME) calculating the response magnitude (in units of percent signal change) of a region of interest (ROI) as a function of assessment trait and the interaction between assessment state and participant group with added covariates of participant age, mean framewise displacement during the scan (mean FD), and the number of usable runs of the experiment along with a random intercept by participant. Intervention group and text are treated as the reference categories. Trait: within-participant average over time; State: within-participant change relative to trait; VWFA: Visual Word Form Area; FFA: Fusiform Face Area; WJ BRS: Woodcock-Johnson Basic Reading Skills; WJ RF: Woodcock-Johnson Reading Fluency; TOWRE: Test of Word Reading Efficiency; WJ MFF: Woodcock-Johnson Math Facts Fluency; Dys Ctrl: Dyslexic Control; Typ Ctrl: Typical Control; FD: Framewise Displacement Significant results are displayed in bold and asterisks indicate the degree of significance ($p < 0.001$: ***, $p < 0.01$: **, $p < 0.05$: *, $p < 0.1$: .). Significant results of $p<0.05$ are indicated with bold font. Note: No corrections for multiple comparisons were made due to the use of small, manually-defined ROIs. Source data are provided in a public data repository.

Text Selectivity ~ Reading Trait + Reading State * Subgroup + Age + Movement + RunNums + (1 | Participant)

| | VWFA1 | | | | | VWFA2 | | | | | FFA1 | | | | | FFA2 | | | | |
|---|---|---|---|---|---|---|---|---|---|---|---|---|---|---|---|---|---|---|---|---|
| | β | Std. Err | DOF | t | p | β | Std. Err | DOF | t | p | β | Std. Err | DOF | t | p | β | Std. Err | DOF | t | p |
| **WJ BRS** | | | | | | | | | | | | | | | | | | | | |
| Intercept (Intervention) | 0.151 | 0.091 | 109 | 1.669 | 0.098. | 0.129 | 0.091 | 96 | 1.422 | 0.158 | -0.331 | 0.081 | 105 | -4.076 | 8.90E-05*** | -0.302 | 0.090 | 82 | -3.374 | 0.001** |
| WJ BRS Trait | 0.000 | 0.001 | 78 | -0.601 | 0.549 | 0.000 | 0.001 | 63 | -0.685 | 0.496 | 0.002 | 0.001 | 84 | 2.877 | 0.005** | 0.003 | 0.001 | 74 | 3.296 | 0.002** |
| WJ BRS State | 0.002 | 0.001 | 211 | 2.438 | 0.016* | 0.002 | 0.001 | 185 | 2.452 | 0.015* | 0.000 | 0.001 | 238 | -0.127 | 0.899 | 0.000 | 0.000 | 224 | 0.522 | 0.602 |
| Group: Dys Ctrl | 0.002 | 0.019 | 75 | 0.113 | 0.910 | 0.012 | 0.017 | 64 | 0.718 | 0.475 | 0.009 | 0.018 | 82 | 0.532 | 0.596 | -0.006 | 0.020 | 72 | -0.285 | 0.777 |
| Group: Typ Ctrl | 0.022 | 0.023 | 73 | 0.964 | 0.338 | 0.029 | 0.022 | 61 | 1.298 | 0.199 | 0.000 | 0.024 | 82 | -0.004 | 0.997 | -0.024 | 0.027 | 73 | -0.900 | 0.371 |
| Age | -0.005 | 0.005 | 73 | -1.047 | 0.298 | 0.001 | 0.005 | 65 | 0.134 | 0.894 | -0.001 | 0.005 | 83 | -0.149 | 0.882 | -0.005 | 0.006 | 72 | -0.981 | 0.330 |
| Movement (Mean FD) | 0.022 | 0.039 | 267 | 0.576 | 0.565 | 0.010 | 0.044 | 238 | 0.237 | 0.813 | -0.072 | 0.033 | 293 | -2.154 | 0.032* | -0.050 | 0.023 | 257 | -2.157 | 0.032* |
| Num Good Runs | 0.013 | 0.007 | 246 | 1.833 | 0.068. | 0.011 | 0.008 | 224 | 1.442 | 0.151 | -0.006 | 0.006 | 283 | -1.019 | 0.309 | -0.001 | 0.004 | 242 | -0.171 | 0.864 |
| Dys Ctrl * WJ BRS State | 0.000 | 0.002 | 210 | -0.147 | 0.883 | -0.004 | 0.002 | 183 | -1.850 | 0.066. | 0.001 | 0.001 | 237 | 0.919 | 0.359 | 0.000 | 0.001 | 224 | -0.052 | 0.958 |
| Typ Ctrl * WJ BRS State | -0.002 | 0.002 | 219 | -1.117 | 0.265 | 0.000 | 0.002 | 194 | 0.169 | 0.866 | -0.001 | 0.002 | 244 | -0.399 | 0.690 | 0.000 | 0.001 | 225 | -0.417 | 0.677 |
| **WJ RF** | | | | | | | | | | | | | | | | | | | | |
| Intercept (Intervention) | 0.059 | 0.072 | 110 | 0.818 | 0.415 | 0.085 | 0.072 | 99 | 1.188 | 0.238 | -0.232 | 0.064 | 111 | -3.591 | 4.93E-04*** | -0.175 | 0.068 | 83 | -2.582 | 0.012* |
| WJ RF Trait | 0.000 | 0.001 | 76 | 0.122 | 0.903 | 0.000 | 0.001 | 60 | 0.094 | 0.926 | 0.002 | 0.001 | 82 | 2.621 | 0.010* | 0.002 | 0.001 | 74 | 3.907 | 2.06E-04*** |
| WJ RF State | 0.002 | 0.001 | 213 | 3.004 | 0.003** | 0.003 | 0.001 | 190 | 3.444 | 7.06E-04*** | 0.000 | 0.001 | 240 | 0.538 | 0.591 | 0.000 | 0.000 | 225 | 0.478 | 0.633 |
| Group: Dys Ctrl | -0.004 | 0.019 | 75 | -0.213 | 0.832 | 0.014 | 0.016 | 62 | 0.829 | 0.410 | -0.003 | 0.017 | 81 | -0.161 | 0.873 | -0.016 | 0.018 | 71 | -0.858 | 0.394 |
| Group: Typ Ctrl | 0.010 | 0.022 | 71 | 0.459 | 0.648 | 0.015 | 0.022 | 58 | 0.711 | 0.480 | 0.004 | 0.023 | 79 | 0.187 | 0.852 | -0.021 | 0.025 | 71 | -0.828 | 0.410 |
| Age | -0.003 | 0.005 | 72 | -0.656 | 0.514 | 0.001 | 0.005 | 63 | 0.188 | 0.852 | -0.002 | 0.005 | 80 | -0.457 | 0.649 | -0.011 | 0.005 | 71 | -2.094 | 0.040* |
| Movement (Mean FD) | 0.059 | 0.039 | 267 | 1.493 | 0.137 | 0.014 | 0.043 | 237 | 0.335 | 0.738 | -0.083 | 0.033 | 292 | -2.499 | 0.013* | -0.070 | 0.023 | 260 | -3.009 | 0.003** |
| Num Good Runs | 0.019 | 0.007 | 244 | 2.634 | 0.009** | 0.009 | 0.007 | 220 | 1.214 | 0.226 | -0.010 | 0.006 | 274 | -1.810 | 0.071. | -0.005 | 0.004 | 238 | -1.269 | 0.206 |
| Dys Ctrl * WJ RF State | 0.002 | 0.002 | 209 | 0.788 | 0.431 | -0.004 | 0.002 | 181 | -1.512 | 0.132 | -0.002 | 0.002 | 233 | -0.855 | 0.393 | -0.003 | 0.001 | 221 | -2.127 | 0.035* |
| Typ Ctrl * WJ RF State | -0.003 | 0.002 | 209 | -1.563 | 0.119 | -0.003 | 0.002 | 182 | -2.070 | 0.040* | 0.000 | 0.001 | 235 | -0.262 | 0.794 | -0.001 | 0.001 | 221 | -0.954 | 0.341 |
| **TOWRE** | | | | | | | | | | | | | | | | | | | | |
| Intercept (Intervention) | 0.099 | 0.079 | 106 | 1.241 | 0.217 | 0.115 | 0.079 | 94 | 1.454 | 0.149 | -0.244 | 0.069 | 111 | -3.532 | 6.03E-04*** | -0.208 | 0.074 | 84 | -2.812 | 0.006** |
| TOWRE Trait | 0.000 | 0.001 | 74 | -0.596 | 0.553 | 0.000 | 0.001 | 60 | -0.467 | 0.643 | 0.002 | 0.001 | 85 | 2.236 | 0.028* | 0.003 | 0.001 | 74 | 3.345 | 0.001** |
| TOWRE State | 0.002 | 0.001 | 209 | 2.712 | 0.007** | 0.003 | 0.001 | 186 | 3.535 | 5.14E-04*** | 0.000 | 0.001 | 237 | 0.432 | 0.666 | -0.001 | 0.000 | 222 | -1.351 | 0.178 |
| Group: Dys Ctrl | 0.000 | 0.020 | 75 | -0.023 | 0.982 | 0.015 | 0.017 | 64 | 0.922 | 0.360 | 0.009 | 0.018 | 81 | 0.485 | 0.629 | -0.003 | 0.020 | 72 | -0.138 | 0.890 |
| Group: Typ Ctrl | 0.025 | 0.026 | 72 | 0.991 | 0.325 | 0.030 | 0.024 | 60 | 1.228 | 0.224 | 0.008 | 0.026 | 84 | 0.295 | 0.769 | -0.031 | 0.028 | 74 | -1.103 | 0.274 |
| Age | -0.003 | 0.005 | 74 | -0.686 | 0.495 | 0.001 | 0.005 | 65 | 0.157 | 0.876 | -0.003 | 0.005 | 82 | -0.572 | 0.569 | -0.010 | 0.006 | 72 | -1.752 | 0.084. |
| Movement (Mean FD) | 0.060 | 0.040 | 262 | 1.506 | 0.133 | 0.015 | 0.044 | 233 | 0.348 | 0.728 | -0.076 | 0.034 | 288 | -2.222 | 0.027* | -0.072 | 0.024 | 255 | -3.010 | 0.003** |
| Num Good Runs | 0.020 | 0.007 | 238 | 2.755 | 0.006** | 0.009 | 0.007 | 215 | 1.269 | 0.206 | -0.007 | 0.006 | 274 | -1.308 | 0.192 | -0.004 | 0.004 | 235 | -0.981 | 0.328 |
| Dys Ctrl * TOWRE State | 0.006 | 0.003 | 209 | 2.224 | 0.027* | -0.001 | 0.002 | 181 | -0.577 | 0.564 | -0.001 | 0.002 | 233 | -0.648 | 0.517 | 0.000 | 0.001 | 220 | -0.073 | 0.942 |
| Typ Ctrl * TOWRE State | 0.000 | 0.002 | 207 | -0.170 | 0.865 | 0.000 | 0.002 | 180 | -0.088 | 0.930 | 0.000 | 0.001 | 232 | -0.305 | 0.761 | -0.001 | 0.001 | 220 | -0.883 | 0.378 |
| **WJ MFF** | | | | | | | | | | | | | | | | | | | | |
| Intercept (Intervention) | 0.088 | 0.071 | 105 | 1.241 | 0.218 | 0.106 | 0.070 | 97 | 1.507 | 0.135 | -0.181 | 0.070 | 104 | -2.595 | 0.011* | -0.136 | 0.076 | 79 | -1.800 | 0.076. |
| WJ MFF Trait | 0.000 | 0.000 | 68 | -0.779 | 0.439 | 0.000 | 0.000 | 57 | -0.435 | 0.665 | 0.000 | 0.000 | 78 | 0.740 | 0.461 | 0.001 | 0.001 | 69 | 1.750 | 0.084. |
| WJ MFF State | 0.001 | 0.001 | 214 | 1.093 | 0.276 | 0.000 | 0.001 | 191 | -0.328 | 0.744 | 0.000 | 0.001 | 241 | -0.024 | 0.981 | 0.000 | 0.000 | 225 | -0.733 | 0.464 |

| | | | | | | | | | | | | | | | | | | | | |
|---|---|---|---|---|---|---|---|---|---|---|---|---|---|---|---|---|---|---|---|---|
| Group: Dys Ctrl | -0.003 | 0.019 | 75 | -0.160 | 0.874 | 0.014 | 0.016 | 63 | 0.838 | 0.405 | 0.002 | 0.018 | 81 | 0.134 | 0.894 | -0.012 | 0.020 | 70 | -0.570 | 0.570 |
| Group: Typ Ctrl | 0.019 | 0.017 | 72 | 1.148 | 0.255 | 0.021 | 0.015 | 62 | 1.328 | 0.189 | **0.046** | **0.018** | **81** | **2.508** | **0.014*** | 0.025 | 0.021 | 70 | 1.213 | 0.229 |
| Age | -0.004 | 0.005 | 73 | -0.758 | 0.451 | 0.000 | 0.005 | 65 | 0.004 | 0.997 | 0.000 | 0.005 | 83 | 0.090 | 0.929 | -0.004 | 0.006 | 71 | -0.750 | 0.456 |
| Movement (Mean FD) | 0.053 | 0.040 | 270 | 1.323 | 0.187 | 0.005 | 0.044 | 239 | 0.122 | 0.903 | **-0.082** | **0.033** | **290** | **-2.460** | **0.014*** | **-0.072** | **0.024** | **258** | **-3.040** | **0.003**** |
| Num Good Runs | **0.022** | **0.007** | **247** | **3.013** | **0.003**** | 0.012 | 0.007 | 223 | 1.562 | 0.120 | -0.008 | 0.006 | 277 | -1.384 | 0.168 | -0.005 | 0.004 | 239 | -1.179 | 0.239 |
| Dys Ctrl * WJ MFF State | -0.002 | 0.003 | 221 | -0.784 | 0.434 | 0.000 | 0.002 | 184 | 0.018 | 0.986 | 0.001 | 0.001 | 237 | 1.007 | 0.315 | 0.001 | 0.001 | 226 | 0.727 | 0.468 |
| Typ Ctrl * WJ MFF State | 0.002 | 0.002 | 212 | 0.869 | 0.386 | 0.000 | 0.002 | 185 | -0.115 | 0.908 | -0.001 | 0.001 | 237 | -0.830 | 0.407 | 0.000 | 0.001 | 223 | 0.062 | 0.951 |

**Table S17 | Longitudinal LME Results for Text Selectivity with State and Trait Scores**

Results of linear mixed effects models (LME) calculating the size (log transformed number of vertices) of a region of interest (ROI) as a function of assessment trait and the interaction between assessment state and participant group with added covariates of participant age, mean framewise displacement during the scan (mean FD), and the number of usable runs of the experiment along with a random intercept by participant. Intervention group is treated as the reference category. Selectivity index is calculated as the difference between activation to text versus non text, divided by the sum of activation to all stimuli. Trait: within-participant average over time; State: within-participant change relative to trait; VWFA: Visual Word Form Area; FFA: Fusiform Face Area; WJ BRS: Woodcock-Johnson Basic Reading Skills; WJ RF: Woodcock-Johnson Reading Fluency; TOWRE: Test of Word Reading Efficiency; WJ MFF: Woodcock-Johnson Math Facts Fluency; Dys Ctrl: Dyslexic Control; Typ Ctrl: Typical Control; FD: Framewise Displacement Significant results are displayed in bold and asterisks indicate the degree of significance (p < 0.001: ***, p < 0.01: **, p < 0.05: *, p < 0.1: .). Significant results of p<0.05 are indicated with bold font. Note: No corrections for multiple comparisons were made due to the use of small, manually-defined ROIs. Source data are provided in a public data repository.

| Participant Demographic Information | | | |
|---|---|---|---|
| | Control Group | | Intervention Group |
| | ***Dyslexic*** | ***Typical*** | |
| N | 19 | 24 | 44 |
| **Reported Annual Household Income** | | | |
| Min | $3,500 | $60,000 | $30,000 |
| Median | $212,500 | $300,000 | $150,000 |
| Max | $1,250,000 | $1,000,000 | $575,000 |
| SD | $267,076 | $210,345 | $115,411 |
| N Not Reported | 2 | 3 | 6 |
| **Race (count)** | | | |
| American Indian or Alaska Native | 1 | 0 | 1 |
| Asian | 3 | 6 | 3 |
| Black or African American | 4 | 0 | 6 |
| Native Hawaiian or Other Pacific Islander | 0 | 0 | 1 |
| White | 11 | 18 | 30 |
| Other | 3 | 1 | 3 |
| **Ethnicity** | | | |
| Hispanic (count) | 6 | 3 | 12 |

**Table S18 | Participant Demographic Information**

Additional demographic information broken down by participant group. All data presented in the table were collected via parents8.csv-report from the participant's first visit to the lab.

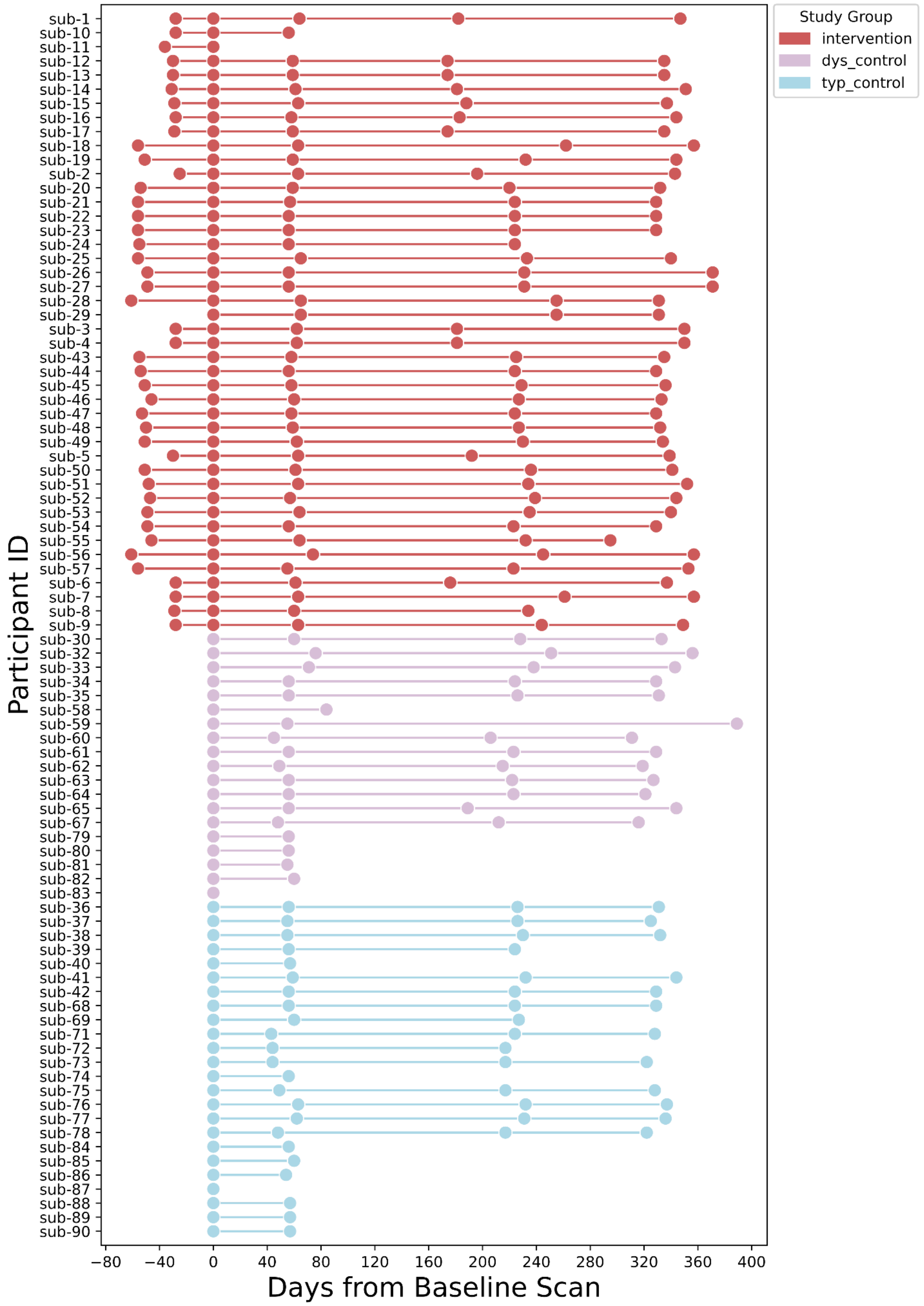


**Figure S4 | Data collection timeline for each participant relative to baseline scan.**
Of the 87 children who participated in the study, 10 participants were lost from long term followup in the study - 4 typical controls, 2 dyslexic controls, and 4 intervention participants. Additionally, 12 control participants were included in the study as part of an extension year for the study grant to compensate for COVID-19 delays and were only brought in for pre- and post-intervention time points. Source data are provided in a public data repository.

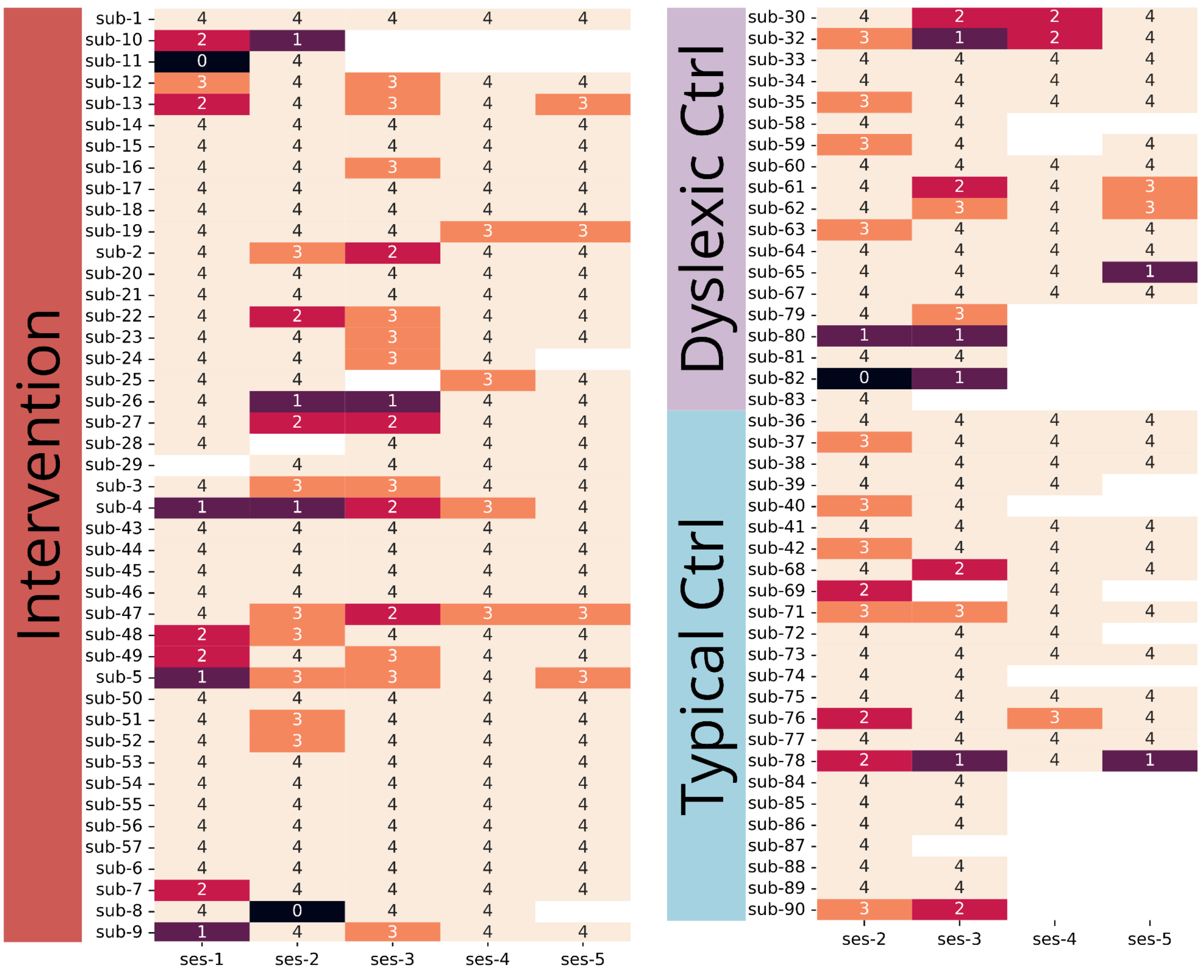

**Figure S5 | Number of low-motion runs of the functional localizer per participant in each time point**

Numbers reflected here are the number of runs collected from each participant after excluding individual runs for high movement. Runs were excluded from analysis if the mean framewise displacement (FD) was 0.5 mm or larger, or if more than 30% of frames had an FD greater than 0.5 mm. Source data are provided in a public data repository.

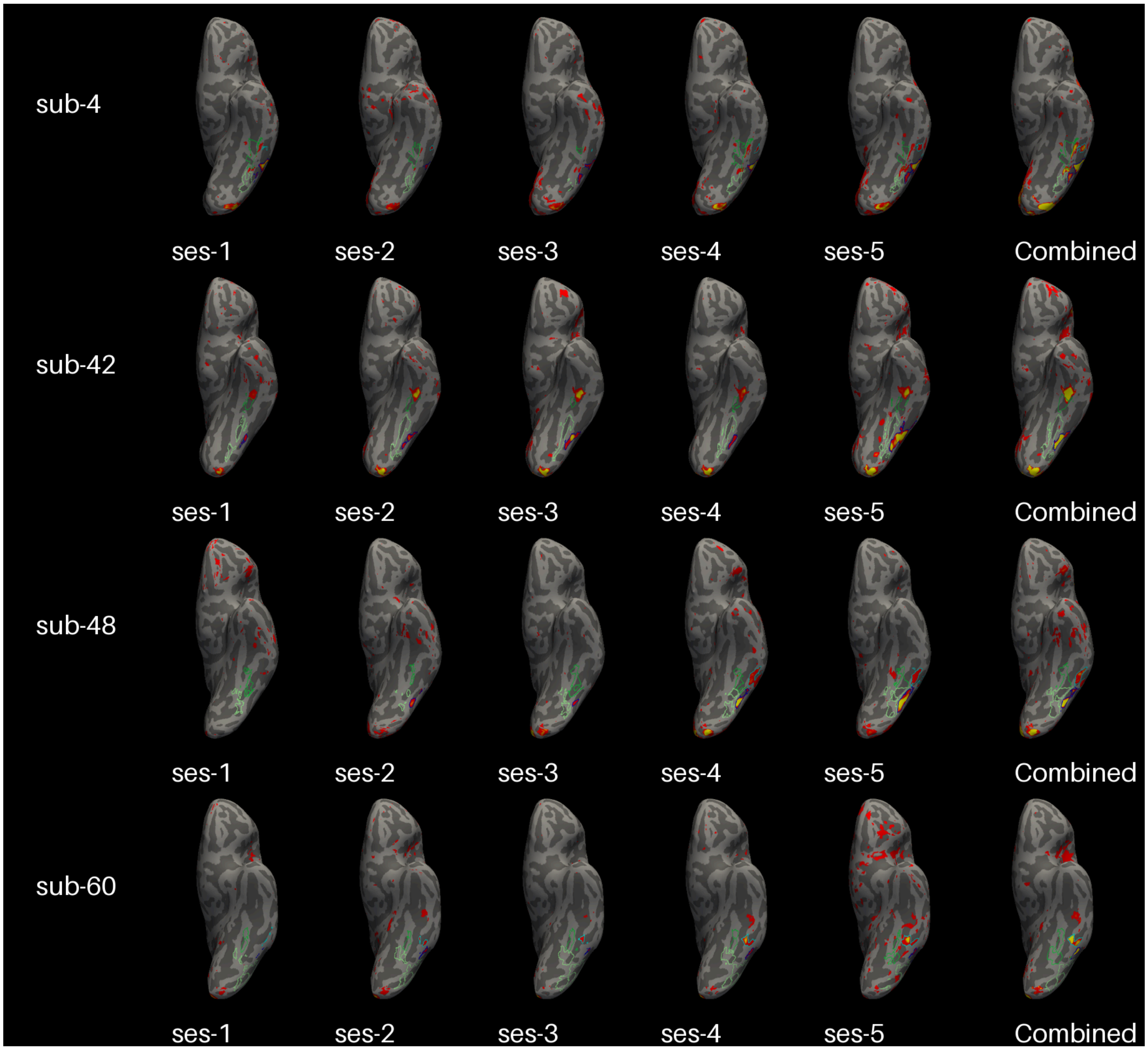


**Figure S6 | ROIs on Text > Other Statistical Maps in Sample Participants**

Regions of interest (ROIs) drawn on the native surface for 4 sample participants for each time point and averaged across time points (combined) projected over a contrast map of text > all other categories thresholded at a t of 3. VWFA-1 is dark blue, VWFA-2 is light blue, FFA-1 is light green, and FFA-2 is dark green. VWFA: Visual Word Form Area; FFA: Fusiform Face Area.

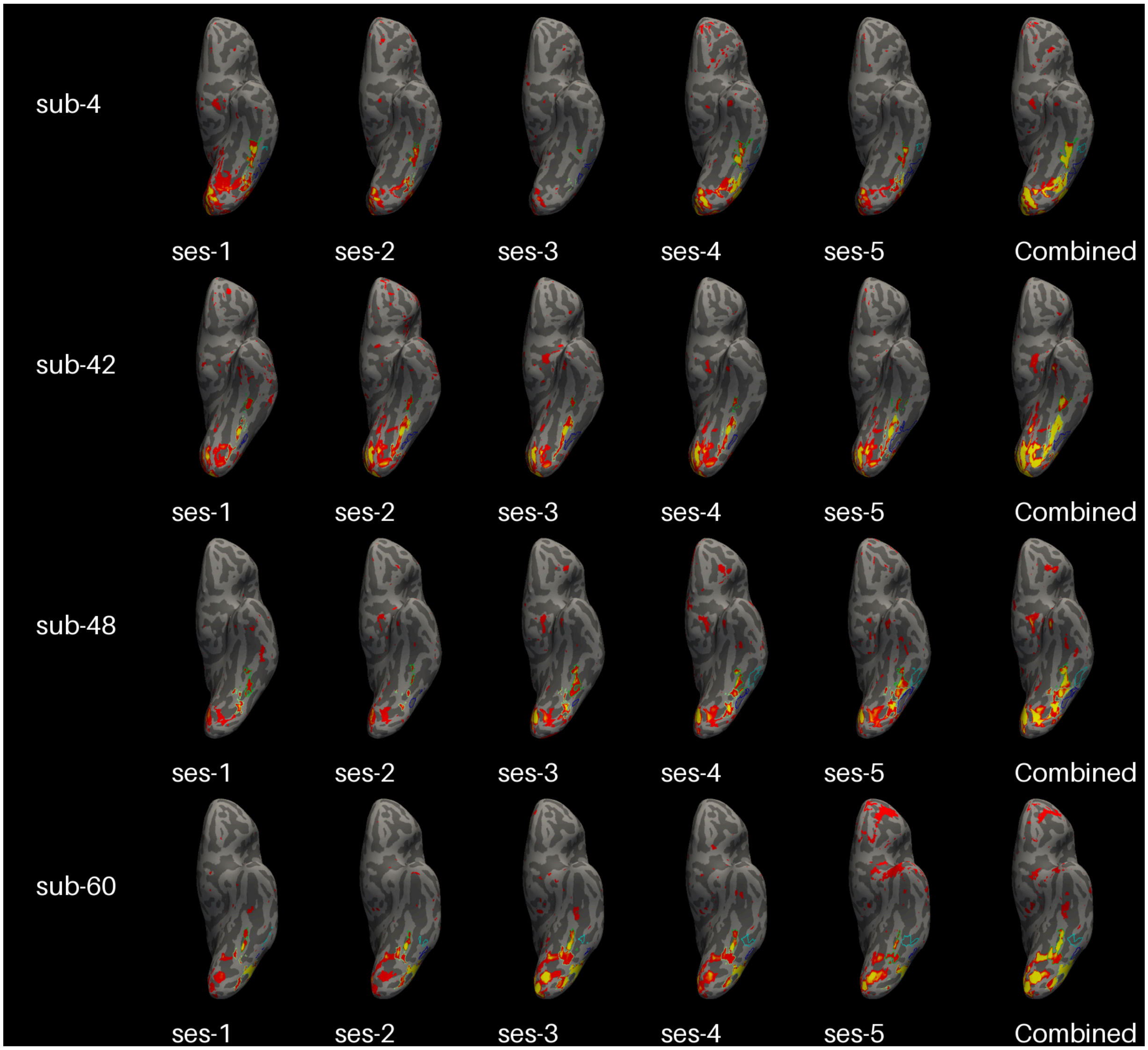


**Figure S7 | ROIs on Faces > Other Statistical Maps in Sample Participants**
Regions of interest (ROIs) drawn on the native surface for 4 sample participants for each time point and averaged across time points (combined) projected over a contrast map of faces > all other categories thresholded at a t of 3. VWFA-1 is dark blue, VWFA-2 is light blue, FFA-1 is light green, and FFA-2 is dark green. VWFA: Visual Word Form Area; FFA: Fusiform Face Area.
